# TMPO promotes cellular dissemination and metastasis in circulating tumor cells

**DOI:** 10.64898/2025.12.27.696673

**Authors:** Arianna Giacobbe, Aleksandar Z. Obradovic, Jinqiu Lu, Soonbum Park, Carlos Pedraz-Valdunciel, Giuseppe Nicolo’ Fanelli, Aunika Zheng, Jaime Y. Kim, Maya Stella Dixon, Jung Seung Nam, Florencia Picech, Caroline Laplaca, Renu K. Virk, Matteo Di Bernardo, Alexander Chui, Juan M. Arriaga, Stephanie Afari, Francisca Nunes de Almeida, Min Zou, Helen Garcia, Brian D. Robinson, Hongshan Guo, Shyamala Maheswaran, Daniel A. Haber, David T. Miyamoto, David M. Nanus, Scott T. Tagawa, Tian Zheng, Massimo Loda, Iok In Christine Chio, Michael Shen, Paraskevi Giannakakou, Andrea Califano, Peter A. Sims, Cory Abate-Shen

## Abstract

Metastasis—the process by which cancer cells spread beyond the primary tumor to distant organs—accounts for the vast majority of cancer-related deaths. To elucidate mechanisms underlying dissemination and metastasis in prostate cancer, we have investigated circulating tumor cells (CTCs) obtained from genetically engineered mouse models (GEMMs). The phenotypic and molecular properties of the CTCs, and organoids derived from these CTCs, closely model the tumor and metastatic phenotypes of their parental GEMMs. Moreover, organoids derived from individual CTCs exhibit molecular and morphological heterogeneity that is associated with distinct metabolic states as well as differences in human prostate cancer outcome. Using computational systems analyses, we have identified *TMPO*, encoding the nuclear membrane protein lamina-associated polypeptide 2 (Lap2), as a key driver of this heterogeneity. *TMPO* activity is upregulated in advanced human prostate tumors, metastases, and CTCs, and is associated with adverse clinical outcome. Our findings indicate that *TMPO* promotes dissemination and metastasis *in vivo* by enhancing survival in conditions of metabolic stress, and reveal a novel mechanistic link between CTC heterogeneity, stress adaptation, and metastatic potential.

## Main text

Although the past two decades have witnessed significant improvements in survival for patients with localized cancer, metastasis remains a leading cause of cancer-related death (*1*). This contrast in outcomes between localized and metastatic disease is particularly striking for prostate cancer—whereas locally-invasive disease rarely results in mortality, metastatic prostate cancer is invariably fatal (*2, 3*). Nowadays, the vast majority (∼90%) of men present with regionally-confined prostate adenocarcinoma having low to intermediate Gleason grade, a pathological assessment of prostate cancer aggressiveness (*4*). Men with low Gleason grade tumors are often managed by active surveillance (*i.e.,* active monitoring without treatment), while those with intermediate Gleason grade tumors can often be cured by surgery or radiation therapy (*2, 3*). For men with recurrent or advanced prostate cancer, the standard of care is androgen deprivation therapy (ADT), based on the dependence of both normal prostate epithelium and prostate cancer on androgen receptor (AR) signaling (*5, 6*). Although initially effective, ADT eventually results in the emergence of castration-resistant prostate cancer (CRPC), so named because of its continued reliance on AR signaling even in the absence of androgens (*6, 7*). Subsequent therapies include second generation anti-androgens, such as enzalutamide; while initially effective, their use can lead to more aggressive disease variants, including neuroendocrine prostate cancer (NEPC) (*8*). The primary site of prostate cancer metastasis is to bone (*9*), however metastasis to soft tissues, including lung and liver, is becoming increasingly prevalent, especially in aggressive prostate cancer variants (*2, 3, 10*). While metastasis is especially common in CRPC (referred to as mCRPC) and NEPC, not all metastatic disease arises at advanced stages. A subset of men present with *de novo* metastases prior to treatment, referred to as hormone sensitive oligometastatic prostate cancer (mHSPC) (*11, 12*).

A shared feature across many cancer types, metastasis represents the culmination of a complex series of events—collectively known as the metastatic cascade—that begins with the escape of tumor cells from the primary site, followed by their entry into the circulation, transit to distant organs, and ultimately colonization to form secondary tumors, known as metastatic lesions (*13–16*). This is a highly inefficient process, with numerous barriers that eliminate most disseminating tumor cells; indeed, only a small fraction of cells that escape the primary tumor successfully establish metastatic lesions (*17–19*). At each step in the cascade, tumor cells undergo phenotypic and molecular reprogramming associated with cellular plasticity, including differences in epithelia-to-mesenchymal transition (EMT) states as well as metabolic adaptations that allow them to survive in hostile microenvironments particularly during circulation (*20–24*).

In principle, investigating these dynamic cellular transitions should be attainable by studying the biological and molecular properties of tumor cells as they transit through the circulation—namely, circulating tumor cells (CTCs). Yet, in practice it has proven challenging to study CTCs, particularly in clinical settings, due to their rarity and inherent difficulties in their detection and isolation (*25*). Although analyses of circulating tumor DNA (ctDNA) offer an alternative means of monitoring disease progression and treatment response (*26–28*), analyses of CTCs provide the unique opportunity to study the molecular and biological characteristics of living tumor cells as they transit from tumors to colonize distant organs (*29*). Another key challenge is CTC complexity, which likely contributes to metastatic potential. In particular, CTCs do not only occur in advanced cancer stages but may also arise early in cancer progression, well before the occurrence of overt metastases (*30*), and they may transit in blood as singlets or multicellular aggregates with other CTCs and/or immune cells (*31, 32*). Yet, despite their heterogeneity, most strategies used to study CTCs are based on detection of specific properties, such as cell size or expression of surface markers (*33, 34*) potentially biasing analyses toward subsets of CTCs enriched for these features.

In the current study, we employed an unbiased approach to isolate and study the biology and molecular heterogeneity of CTCs as they arise *in vivo* across distinct tumor contexts and during temporal progression of metastasis. Specifically, we isolated CTCs from genetically-engineered mouse models (GEMMs) of metastatic prostate cancer by virtue of lineage-marking of prostatic tumor cells, which enabled the identification of a range of CTC phenotypes. We have found that the incidence and phenotype of CTCs, as well as organoids derived from them, faithfully model the tumor and metastatic phenotypes from which they originate. Analyses of individual CTCs (*i*CTCs) and individual organoids derived from these CTCs (*i*Organoids) revealed molecular and phenotypic heterogeneity associated with clinical outcome of human prostate cancer. A key driver of this molecular and morphological heterogeneity is *TMPO*, which encodes the nuclear membrane protein lamina-associated polypeptide 2 (Lap2). Up-regulation of TMPO activity is associated with adverse outcome in human prostate cancer, and promotes dissemination and metastasis *in vivo* through its ability to support cell survival under conditions of metabolic stress. Together these findings reveal a mechanistic link between CTC heterogeneity, stress adaptation, and metastatic potential mediated by TMPO activity.

## Results

### Prostate tumors from metastatic GEMMs display distinct molecular features

To study the relationship between tumorigenesis, cellular dissemination and metastasis, we have investigated a unique series of prostate cancer GEMMs that display a range of metastatic phenotypes, thereby enabling investigation of both the temporal and phenotypic evolution of metastasis *in vivo* (Fig. 1A-E; Fig. S1A-E; Table S1; Dataset 1A) (*35–39*). These GEMMs employ an inducible *Nkx3.1^CreERT2^* allele, which enables spatial and temporal control of gene recombination specifically in prostatic luminal cells (*40*), resulting in heterozygous loss-of-function of *Nkx3.1*, as is commonly observed in many human prostate cancers (*41, 42*). The baseline (control) model is the *NP* GEMM (for *Nkx3.1^CreERT2/+^; Pten^flox/flox^; R26R-CAG^LSL-EYFP/+^*), which has conditional loss-of-function of *Pten* (*39*), a frequent alteration in human prostate cancer (*42, 43*). *NP* mice display well-differentiated prostate adenocarcinoma that is non-lethal and non-metastatic (Fig. 1A, D, E; Fig. S1A-E; Table S1; Dataset 1A) (*37, 39*). Of particular importance for our studies, this GEMM series also has a conditional YFP reporter allele (*R26R-CAG^LSL-EYFP^*), allowing lineage marking of prostatic luminal cells following Cre-mediated recombination (*44*). The YFP lineage mark enables direct visualization and subsequent phenotypic and molecular characterization of tumor, disseminated, and metastatic cells as they arise *de novo* in the whole organism (Fig. 1B) (*36, 37*).

**Figure 1.**
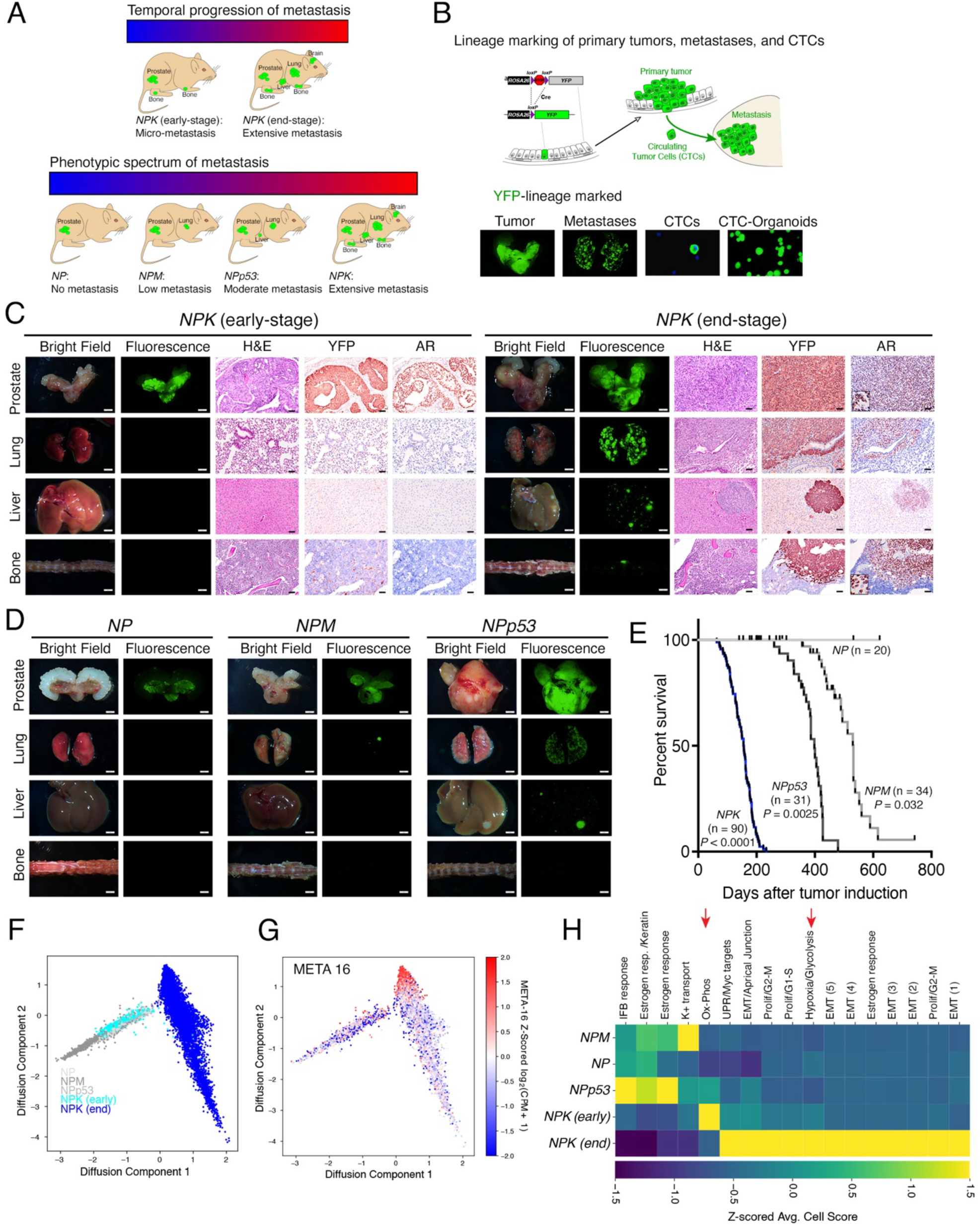
Prostate tumors from metastatic GEMMs display distinct molecular features. **(A)** Schematic showing the genetically-engineered mouse models (GEMMs) employed to study the temporal progression *(top)* or phenotypic spectrum *(bottom)* of prostate cancer metastasis. The relative metastatic phenotypes of the GEMMs are illustrated. **(B)** Strategy for lineage-marking of prostatic epithelium, which gives rise to yellow fluorescent protein (YFP)-marked tumor cells, and their descendant circulating tumor cells (CTCs) and metastases. **(C)** Representative images of prostate, lung, liver and bone from *NPK* early-stage and *NPK* end-stage mice. Shown are whole mount images of bright-field and *ex vivo* fluorescence, and sections of histology (H&E) and immunostaining (IHC) for YFP and AR (Androgen receptor), as indicated. (n = 3-4 mice/group for the IHC). **(D)** Representative bright-field and *ex vivo* fluorescence images of prostate, lung, liver and bone for the indicated GEMMs. In panels **C** and **D**, scale bars represent 2 mm for the whole mounts and 50 μm for the H&E and IHC. **(E)**. Kaplan-Meier survival curve showing overall survival for the indicated GEMMs. *P* values were calculated using a two-tailed log-rank test compared to *NP* mice (control). (**F)** Diffusion Component (DC) analysis of scRNA-sequencing data from prostate tumors derived from the indicated GEMMs. (**G)** Z-Score normalized expression of the META 16 gene signature (*36*) projected onto the DC plots. **(H)** Heatmap of hierarchical clustering showing average cell scores for top genes in each combination of GEMM signature, as derived from scHPF factor analysis (see *Methods*). The red arrows highlight key pathways of interest. See also Figure S1, Table S1, Dataset 1A, Dataset 2.

The principal focus of our study is the *NPK* GEMM (for *Nkx3.1^CreERT2/+^; Pten^flox/flox^; Kras^lsl-G12D/+^; R26R-CAG^LSL-EYFP/+^*) (*36*), which incorporates conditional activation of *Kras* via the widely-used *Kras^G12D^* allele (*45, 46*). Although *KRAS* mutations are relatively infrequent in human prostate cancer (*47*), *NPK* mice effectively model *RAS* pathway activation (*36*), which is prevalent in aggressive, poor-prognosis human prostate cancer (*36, 48, 49*). *NPK* mice develop lethal prostate cancer with fully penetrant metastases that arise in multiple distant sites, including lung, liver, brain, and bone (Fig. 1A, C, E; Fig. S1A-E; Table S1; Dataset 1A) (*36, 38*). These metastases are readily evident as visualized by *ex vivo* YFP fluorescence imaging of whole tissues, as well as histopathological analyses of immunostaining for the YFP lineage mark, and the androgen receptor (AR), a canonical marker of prostate cancer, both of which are prominently expressed in primary tumors and metastatic lesions, with nuclear AR expression particularly high in the bone lesions (Fig. 1C; Fig. S1A,C). While the *NPK* mice develop extensive metastases at terminal stages (approximately 5-7 months post-tumor induction; hereafter *NPK* end-stage mice), at earlier stages of disease progression (3 months post-tumor induction; hereafter *NPK* early-stage mice) these mice exhibit well-differentiated prostate adenocarcinoma with a low incidence of metastasis that occur as micro-metastases to bone (Fig. 1A, C; Fig. S1A-E; Table S1) (*36, 38*). For comparative analyses of their metastatic phenotypes, we examined two additional GEMMs: the *NPM* mice (for *Nkx3.1^CreERT2/+^;Pten^flox/flox^; Hi-MYC; R26R-CAG^LSL-EYFP/+^*) (*35*) and the *NPp53* mice (for *Nkx3.1^CreERT2/+^; Pten^flox/flox^; Trp53^flox/flox^; R26R-CAG^LSL-EYFP/+^*) (*37*). Both models develop metastatic prostate cancer, albeit with lower penetrance and distinct organ tropism compared with the *NPK* mice (Fig. 1A, D, E; Fig. S1A-E; Table S1; Dataset 1A). The *NPM* mice, which have prostate-specific gain-of-function of human c-MYC (*35, 50*), a hallmark of human prostate cancer (*51*), develop well-differentiated prostate adenocarcinoma with a low-incidence of metastasis, occurring primarily in the lung (Fig. 1A,D,E; Fig. S1A-E; Table S1; Dataset 1A). The *NPp53* mice, which have conditional loss-of-function of *Tp53* (*37*), another frequent alteration in human advanced prostate cancer (*42, 43*), develop poorly differentiated prostate adenocarcinoma with a moderate incidence of metastasis, predominantly to the lung and, less frequently, to the liver (Fig. 1A,D,E; Fig. S1A-E; Table S1; Dataset 1A) (*35, 37*). Notably, although they have a lower incidence of metastasis than either the *NPM* or the *NPp53* mice, metastases in the *NPK* early-stage mice occur exclusively as micro-metastasis to bone (Fig. S1D,E) (*36, 38*).

To uncover molecular differences underlying the distinct metastatic phenotypes of these GEMMs, we performed single-cell RNA sequencing (scRNA-seq) of their prostate tumors using a 10X Genomics platform (see *Methods* and Dataset 1A, B). Specifically, we compared *NPK* early-stage versus *NPK* end-stage tumors to identify transcriptional changes associated with temporal progression, and we analyzed tumors from the *NP, NPM*, *NPp53*, and *NPK* end-stage mice to assess phenotypic differences across the models (Fig. 1F-H; Fig. S1F-H; Dataset 1A, B; Dataset 2A, B). To focus on tumor cell–intrinsic programs associated with cellular dissemination and metastasis, we specifically analyzed the YFP-expressing tumor cells within the scRNA-seq datasets (see *Methods* and (*36*)). We found that tumor cells from *NPK* end-stage mice clustered separately from those of the *NPK* early-stage mice and all of the other GEMMs (*NP*, *NPM*, *NPp53*) (Fig. S1F, G). Notably, *NPK* end-stage tumor cells were distinguished by their high-level and widespread expression of vimentin (Fig. S1F, G), a marker of EMT (*52*).

To further define transcriptomic differences in tumor cells from *NPK* end-stage mice compared with those from the *NPK* early-stage mice and the other GEMMs, we applied single-cell hierarchical Poisson factorization (scHPF), an unsupervised probabilistic matrix factorization algorithm (*53*). scHPF identifies interpretable co-expression patterns—gene signatures that are expressed by the same cell—that reflect specific pathways or cell states. For each gene signature, scHPF assigns a score to each cell, reflecting the level at which a cell is implementing a given co-expression pattern. We visualized the resulting matrix of cell scores in two dimensions using diffusion component analysis (Fig. 1F, G; Dataset 2; and see *Methods*) (*54*). This analysis revealed that *NPK* end-stage tumor cells clustered distinctly from those of all the other GEMMs—namely the *NP, NPM*, *NPp53*, and *NPK* early-stage—which form overlapping clusters (Fig. 1F). Although the *NPK* end-stage tumor cells clustered separately and uniformly expressed the EMT marker vimentin (see Fig. S1F, G), only a subset exhibited strong enrichment for META-16 (Fig. 1G), which is a conserved gene signature predictive of poor prognosis for metastatic prostate cancer in human patients (*36*). These findings suggest that not all *NPK* end-stage tumor cells have equivalent metastatic potential.

The scHPF analysis identified 17 distinct gene signatures (i.e., factors) enriched in the *NPK* end-stage tumor cells compared with the other models (*NP, NPM*, *NPp53*, and *NPK* early-stage) (Dataset 2A). In addition to assigning a score to each cell for each scHPF signature, we also obtain a score for each gene. Thus, for each signature, we can rank all genes in order of their importance to the signature. We performed Hallmark pathway analysis on the top 100 scoring genes from each signature to interpret them in terms of pathway enrichment (see *Methods;* Dataset 2B). This revealed that five of these signatures were significantly enriched for EMT–associated genes (q<0.002 for all five EMT signatures; Fig. 1H). Consistent with this finding, *Vimentin, Foxo1, Zeb1,* and *Fibronectin* were markedly upregulated in the *NPK* end-stage tumor cells compared with those of the other GEMMs (Fig. S1H). Notably, the *NPK* end-stage tumor signatures show significant enrichment for genes associated with a hypoxia/glycolysis program, whereas the *NPK* early-stage tumors were enriched for genes associated with oxidative phosphorylation (q=7x10^-9^; Fig. 1H), suggesting that, in addition to EMT, temporal progression of metastasis involves metabolic reprogramming within the primary tumor.

### Circulating tumor cells recapitulate the metastatic phenotype of their parental GEMMs

To further investigate the relationship between tumorigenesis, dissemination, and metastasis we studied the molecular and biological characteristics of circulating tumor cells (CTCs) isolated from the blood of the GEMMs (Fig. 2A-E; Fig. S2A-H; Table S2A, S2B; Dataset 1A). We identified CTCs by virtue of their YFP lineage mark (see Fig. 1B), thus providing an unbiased isolation criterion independent of any prior molecular or phenotypic assumptions about the putative CTCs. Using the *NPK* mice to establish experimental parameters, we isolated YFP-marked cells from blood following red blood cell (RBC) depletion, yielding samples comprised of YFP-positive cells and un-marked white blood cells (see *Methods*). The RBC-depleted blood samples were plated and imaged by live fluorescence microscopy to visualize YFP-expressing cells (Fig. S2A). To confirm the tumor origin of the YFP-marked cells, we collected RBC-depleted blood samples by Cytospin and performed immunofluorescence staining (Fig. 2B). We found that the YFP-marked cells expressed canonical prostate tumor markers, including AR and the luminal cytokeratin 8 (Ck8) (Fig. 2B), confirming their identity as *bona fide* CTCs.

**Figure 2.**
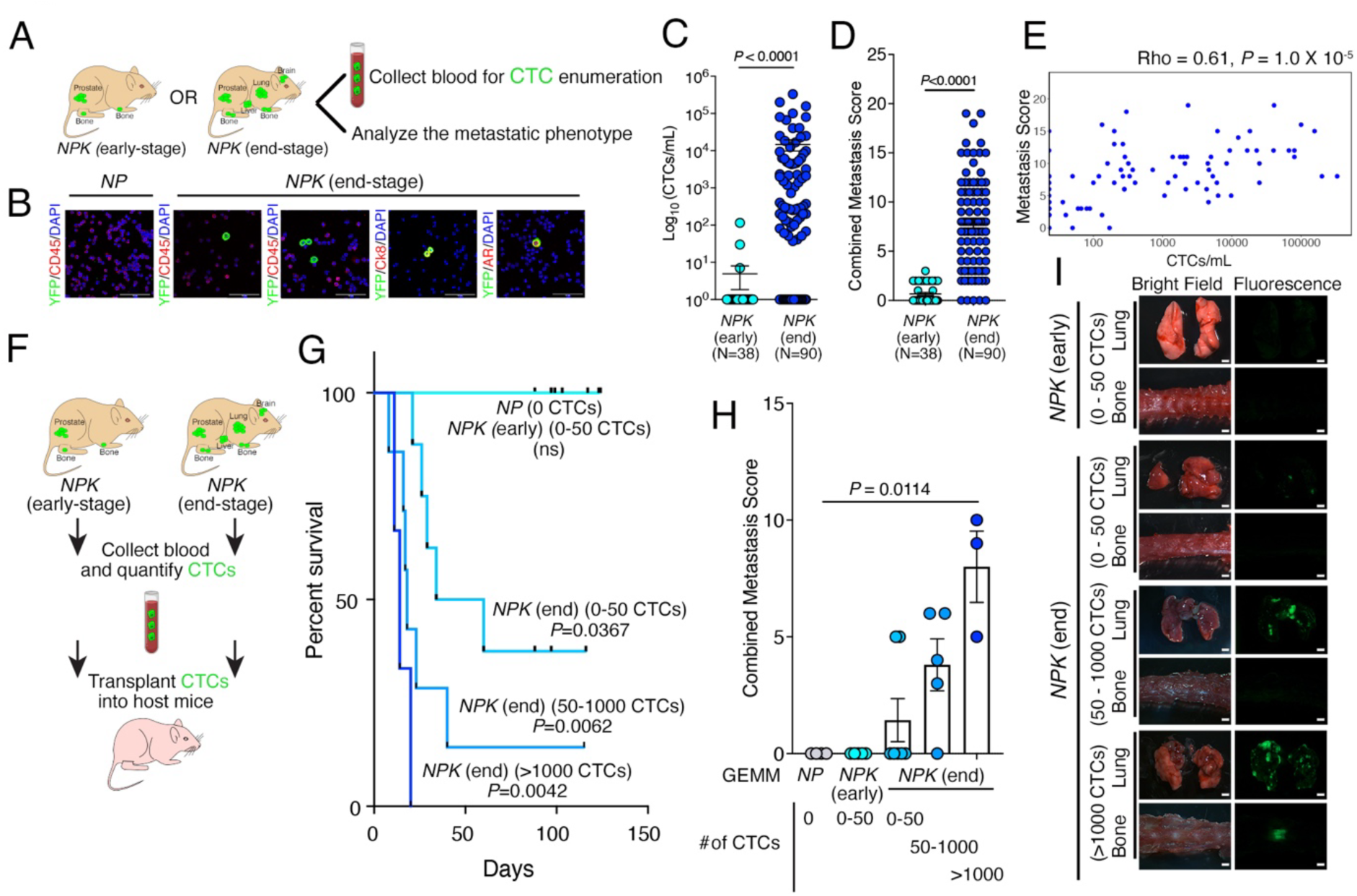
Circulating tumor cells (CTCs) recapitulate the metastatic phenotype of their parental GEMMs. **(A).** Schematic illustrating the strategy for isolation and analyses of CTCs from the indicated GEMMs. **(B)** Immunofluorescence staining of CTCs from *NPK* end-stage mice with the indicated markers. *NP* derived blood samples served as a negative control. Scale bars, 50 μm. (**C)** Scatter dot plot showing the number of CTCs/mouse reported on a logarithmic scale as CTCs/mL, for the *NPK* early-stage and *NPK* end-stage mice, as indicated. CTCs were quantified by counting live YFP-marked cells which were visualized by fluorescence microscopy. **(D)** Scatter dot plot showing the combined metastasis score for each *NPK* early-stage and *NPK* end-stage mouse, as indicated. A semi-quantitative metastasis score was assigned to each mouse based on visualization of metastases and micro-metastases using epifluorescence imaging (see *Methods*). In panels **C** and **D**, data represent mean ± SEM. *P* values were calculated using the two-tailed Mann-Whitney U test. **(E)** Scatter plot depicting the Spearman correlation between metastasis score and CTC count (CTCs/mL) for each *NPK* end-stage mouse. *P* values were calculated from a two-sided permutation test. **(F-I)** Transplantation of CTCs from *NPK* early-stage and *NPK* end-stage mice *in vivo.* **(F)** Strategy: Following quantification of CTCs in the blood of *NPK* early-stage and *NPK* end-stage mice, the corresponding CTC samples were transplanted into non-tumor-bearing recipient host mice via intracardiac injection and the recipient hosts were monitored for survival. **(G)** Kaplan-Meier survival curves showing overall survival of recipient hosts following CTC transplantation from the indicated groups (*n* = 4 for *NP* (0 CTCs); *n* = 5 for *NPK* early-stage (0-50 CTCs); for the *NPK* end-stage, *n* = 8 for (0-50 CTCs); *n* = 7 for (50-1000 CTCs); *n* = 3 for (>1000 CTCs)). *P* values were calculated using a two-tailed log-rank test compared to the *NPK* early-stage group. **(H)** Bar-dot plot showing combined metastasis scores for the indicated host mice. *P* values were calculated using the Kruskal-Wallis test with Dunn’s multiple comparison correction compared to *NP* group. Data represent mean ± SEM. (**I)** Representative bright-field and *ex vivo* fluorescence images of lung and bone from recipient host mice from the indicated groups. Scale bar, 8 mm. See also Figure S2, Table S2, Dataset 1A and 1C.

We quantified CTC incidence (*i.e.,* the number of mice with CTCs) and enumeration (*i.e.,* number of CTCs/mL) using two independent approaches, namely, direct cell counting by fluorescence microscopy and FACS (Fig. S2A-E; Table S2A, S2B). Additionally, we compared CTC samples collected from either intracardiac or submandibular bleeds; while not significantly different the former tended to yield higher CTCs counts (Fig. S2D; Table S2B). Unless otherwise indicated, all subsequent studies used RBC-depleted YFP-marked blood samples from intracardiac bleeds, hereafter referred to as “CTC samples”.

Using these parameters, we analyzed CTC samples from the various GEMMs by direct plating followed by visualization of live YFP fluorescence, which revealed striking differences in incidence and enumeration of CTCs across the models (Fig. 2C; Fig. S2A-E; Table S2A; Dataset 1A). Specifically, we did not detect CTCs in the *NP* mice (0 of 20) while only a small number of *NPK* early-stage mice harbored CTCs (5%, n=2/38), each having relatively few YFP-positive cells (<10 cells/mL). *NPM* mice exhibited CTCs in approximately one-fourth of cases (24%, 8 of 34; mean=116 cells/mL) and *NPp53* mice in more than one-third of cases (39%, 12 of 31; mean=613 cells/mL). The *NPK* end-stage mice had the highest CTC burden, with the majority of mice harboring CTCs (74%, 67 of 90) and their enumeration revealing a much broader range of YFP-positive cells (from 10^2^ to 10^6^/mL; mean = 14638 cells/mL) (Fig. 2C, Fig. S2A, B, E; Table S2A; Dataset 1A). Notably, CTCs from *NPK* early-stage, *NPM*, and *NPp53* mice appeared almost exclusively as singlets, with rare cases of CTC doublets or multicellular aggregates (Fig. S2A; Table S2A). In contrast, multicellular aggregates were observed in approximately 12% (11 out of 90) of CTCs from the *NPK* end-stage mice (Fig. S2A; Table S2A).

To investigate the relationship between CTC incidence and enumeration and the metastatic phenotype of their parental GEMMs, we used a semi-quantitative metastasis score, that incorporates the presence, tissue distribution, and burden of metastatic lesions (see *Methods* and (*36*)). This comparison between CTC incidence and enumeration with the metastasis score revealed that the CTC phenotype mirrored the metastatic phenotype of their parental GEMM. Specifically, *NP* mice exhibited neither CTCs nor metastases, while *NPK* early-stage mice had low CTC incidence and correspondingly low metastasis scores (Fig. 2C, D; Fig. S2E, F). Likewise, *NPM* and *NPp53* mice, which have low to moderate CTC incidence have proportionally low to moderate metastasis scores (Fig. S2E, F). In contrast, the *NPK* end stage mice, which exhibited the highest CTC incidence and enumeration had the highest metastasis scores (Fig. 2C, D; Fig. S2E, F). Importantly, metastasis score was significantly correlated with both CTC incidence and enumeration, particularly for the *NPK* end-stage mice (Rho = 0.61, *P*= 1.0 X 10^-5^) (Fig. 2E; Fig. S2G, H). These findings demonstrate that CTC incidence and abundance closely parallel the metastatic potential of the parental GEMMs from which they originate.

To directly assess the metastatic potential of the CTCs we transplanted blood from the GEMMs into healthy, tumor-free recipient hosts (Fig. 2F-I; Fig. S2I, J; Dataset 1A,1C). Following euthanasia, we obtained blood (minimum of 200 μl) from *NPK* end-stage mice, which have a wide range in the number of CTCs per mouse, *NPK* early-stage mice, which consistently have low numbers of CTC, or as controls *NP* mice, which do not have CTCs (see Fig. 2C; Fig. S2E; Table S2A). For each donor mouse, the number of CTCs was determined in half of the CTC sample (100 μl) as described above. The remainder of the CTC sample (100 μl) was transplanted via intracardiac injection into recipient host mice, which were monitored for survival over 150 days (Fig. 2F, G).

All host mice transplanted with CTC samples from *NPK* end-stage donors died by 120 days, whereas none of those transplanted with CTC samples from *NPK* early-stage or *NP* donors exhibited lethality (composite *P* value = 0.0022; Fig. 2G). Notably, while host mice transplanted with CTCs from *NPK* end-stage mice displayed lethality over a wide range of CTCs (from <50 to >1000), those receiving higher numbers succumbed most rapidly (*P* = 0.0042; Fig. 2G). Moreover, whereas even the lowest numbers of CTCs (<50) from *NPK* end-stage donors resulted in lethality, equivalent numbers of CTCs (<50) from *NPK* early-stage donors displayed no lethality (*P* = 0.0367; Fig. 2G).

Post-mortem analyses revealed that host mice transplanted with CTCs from *NPK* end-stage, but not early-stage, donors developed metastases to multiple organs, including lung and bone, as determined by gross examination and histological analyses (Fig. 2H, I; Fig. S2J). Notably, the metastasis scores of transplanted host mice were strongly anticorrelated with their survival (Rho = —0.7804, *P* <0.0001; Fig. S2I), indicating that lethality resulted from metastatic disease. Taken together these findings reveal that the incidence, abundance, and metastatic phenotype of CTCs closely mirror the metastatic phenotype of their corresponding parental tumors including the capability of generating lethal metastases when transplanted into otherwise tumor-free hosts.

### CTC-organoids retain the tumor and metastatic phenotypes of their parental GEMMs

To further investigate the biological and molecular properties of the CTCs, we generated organoids from CTC samples obtained from the GEMMs (Fig. 3A-F; Fig. S3A, B; Fig. S4A-I; Table S3; S4; Dataset 1A, 1D, 1E). Starting from established methods for generating mouse prostate organoids (*55*), we optimized culture conditions for CTC-derived organoids using samples from *NPK* end-stage mice. We assessed the incidence of CTC-organoid generation (i.e., the number of CTC samples that yield CTC-derived organoids) using three independent methods: (*1*) direct plating of CTC samples in organoid media; (*2*) depletion of CD45-positive immune cells prior to plating; or (*3*) FACS isolation of live YFP-positive cells from CTC samples followed by plating (Fig. S3A; Table S3; Dataset 1D). All three methods yielded comparably high CTC-organoid incidence (> 70%) and the resulting CTC-organoids generated from each method displayed similar histological phenotypes, as shown by immunostaining for AR, cytokeratins (Pan-Ck), vimentin, and Ki67 (Fig. S3A).

**Figure 3.**
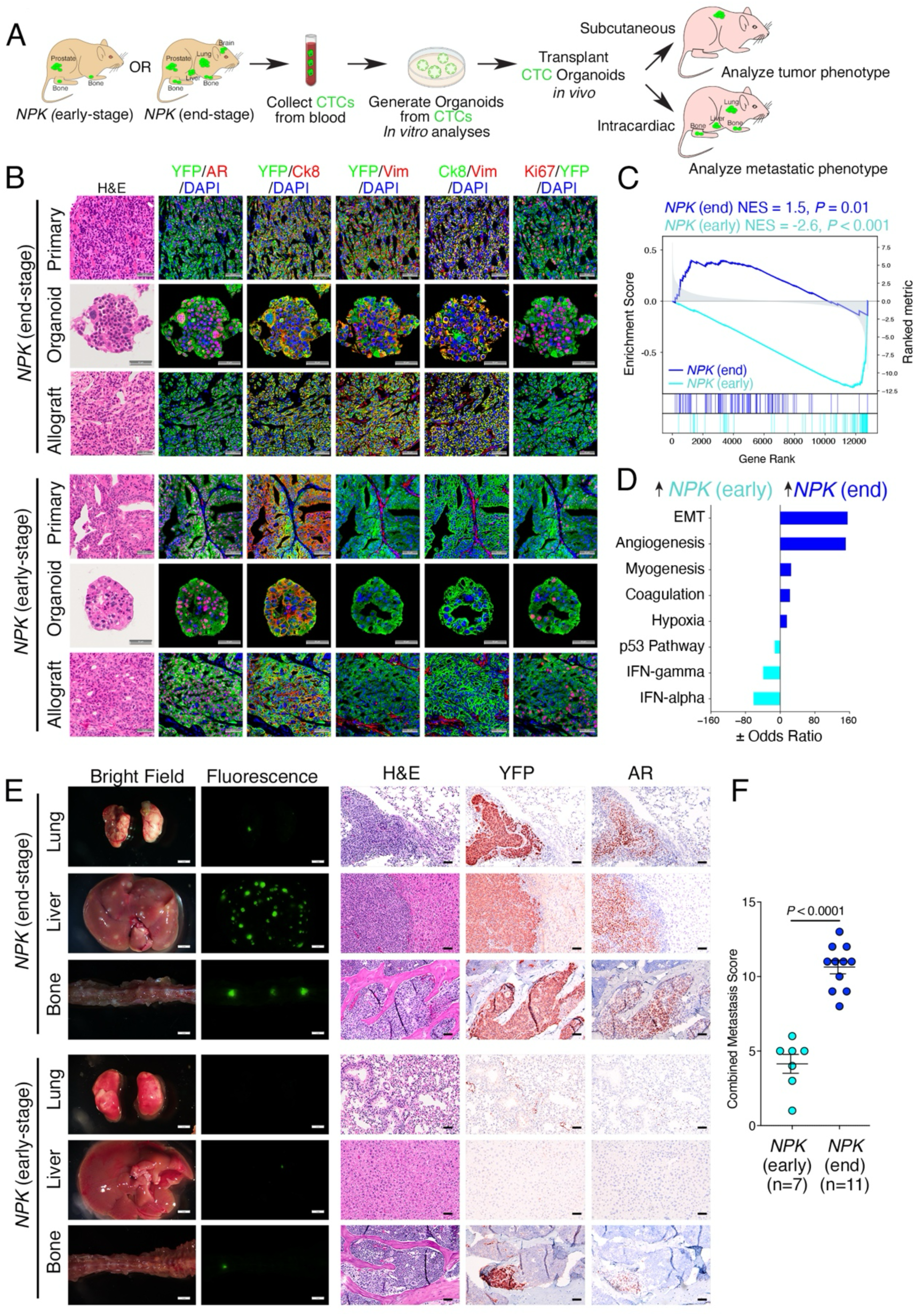
CTC-organoids retain the tumor and metastatic phenotypes of their parental GEMMs. **(A)** Experimental strategy: Blood samples were collected from *NPK* early-stage and *NPK* end-stage mice and red blood cell (RBC)-depleted samples were plated directly in organoid culture conditions (see *Methods*). CTC-derived organoids were analyzed *in vitro* or implanted into non-tumor-bearing recipient host mice via subcutaneous or intracardiac injection for *in vivo* analyses. **(B)** Representative images of matched pairs from the parental tumor (primary) and the corresponding CTC-derived organoid lines propagated *in vitro* (organoid) or grown as allografts *in vivo* (allograft). The images show H&E or immunofluorescence staining for: DAPI (4,6-diaminidino-2-phenylindole), AR, Ck8 (Cytokeratin 8), Vim (Vimentin), YFP and Ki67. (n = 4 for *NPK* early-stage; n = 5 for *NPK* end-stage). Scale bar, 50 mm. **(C)** Gene set enrichment analyses (GSEA) of differentially expressed genes from scRNA-sequencing comparing *NPK* early-stage versus *NPK* end-stage primary tumors with their corresponding CTC-derived organoids. Normalized enrichment scores (NES) and *P* values were estimated using 1,000 gene permutations. **(D)** Hallmark pathway enrichment analysis of the leading-edge genes from the GSEA in **C**. **(E)** Metastatic potential of CTC-derived organoids from *NPK* early-stage and *NPK* end-stage following intracardiac injection into recipient hosts. Representative images of lung, liver and bone, showing whole mount images of bright-field and *ex vivo* fluorescence, and sections of H&E and IHC for YFP and AR, as indicated. (n = 3-4 mice/group for the IHC). Scale bars represent 2 mm for the whole mount images, and 50 μm for the H&E and IHC images. **(F)** Dot plot showing the combined metastasis score of recipient host mice from panel **E**. For panels **E** and **F**, *n* = 7 and *n* = 11 recipient host mice were analyzed from the *NPK* early-stage and *NPK* end-stage organoids, respectively. *P* value in panel **F** was calculated using a two-tailed Mann-Whitney U test. Data are presented as mean ± SEM. See also Figure S3, S4, Table S3, S4 Dataset 1A, 1D-1F, Dataset 3.

We further evaluated organoid incidence using CTC samples isolated from intracardiac (arterial) versus submandibular (venous) blood. Although the occurence of CTCs and the histological features of the resulting CTC-organoids were similar between the two blood sources, the incidence of organoid generation was significantly lower for submandibular samples (*P* < 0.0001; Fig. S3B; Table S3; Dataset 1D), consistent with enhanced survival of CTCs in arterial relative to venous blood (*56*). Therefore, for most subsequent studies, CTC-organoids were generated by direct plating of CTC samples from intracardiac bleeds, which provided the highest incidence while minimizing sample manipulation.

We attempted to generate CTC-derived organoids from each of the GEMMs (Fig. S4A) and observed that organoid incidence was significantly correlated with the corresponding incidence of CTCs (Rho = 0.4457; *P* = 0.0073; Fig. S4B; Table S4; Dataset 1E). Specifically, samples from *NP* mice which had no CTCs yielded no CTC-organoids (0 of 4), the *NPK* early-stage mice which have few cases of CTCs also had few CTC-derived organoids (6%, 2 of 33), and we observed low to moderate efficiencies for the *NPM* (20%, 2 of 10) and *NPp53* (56%, 5 of 9) models. In contrast, *NPK* end-stage mice, which have a high incidence of CTCs, displayed a similarly high incidence of CTC organoids (75%, 40 of 53; Fig. S4A, Table S4; Dataset 1A).

To investigate their biological and molecular properties, we established CTC-derived organoid lines from *NPK* early-stage, *NPK* end-stage, and *NPp53* mice and analyzed their growth, histological, and molecular phenotypes *in vitro* and *in vivo* (Fig. 3A-F; Fig. S4C–I). To investigate their molecular properties, we performed scRNA-seq on CTC-organoid lines from *NPK* early-stage and *NPK* end-stage mice (Fig. 3C, D; Dataset 3A-C). Gene set enrichment analysis (GSEA) revealed significant overlap between genes differentially expressed in the tumor cells and those differentially expressed in the corresponding CTC-organoid lines (NES = 1.5, *P* = 0.01 for the *NPK* end-stage; NES = - 2.6, *P* < 0.001 for the *NPK* early-stage; Fig. 3C), indicating that key transcriptomic differences between early-stage and end-stage tumor cells are preserved in their CTC-organoid counterparts. Notably, pathway analysis of leading-edge genes showed significant enrichment of EMT and hypoxia programs in CTC-organoid lines derived from *NPK* end-stage mice relative to those from *NPK* early-stage mice (Fig. 3D), consistent with the enrichment of these programs in the *NPK* end-stage primary tumor cells (see Fig. 1H).

The CTC-organoid lines also recapitulated key features of their respective parental tumors, while retaining key differences that distinguish the tumors from each other. Consistent with their parental tumors, CTC-organoid lines from *NPK* end-stage—but not from *NPK* early-stage or *NPp53* mice—exhibited robust expression of vimentin (Fig. 3B; Fig. S4H, I). Moreover, Ki67 expression was highest in the *NPK* end-stage tumors and their corresponding CTC-organoids, intermediate in *NPp53* tumors and organoids, and lowest in *NPK* early-stage tumors and organoids (Fig. 3B; Fig. S4H, I).

Given these findings, we asked whether CTC-organoid lines grown *in vivo* recapitulate the tumor and metastatic phenotypes of their corresponding GEMMs. To assess tumor growth, intact CTC-organoids (i.e., without dissociation) from *NPK* early-stage, *NPK* end-stage, and *NPp53* mice were implanted subcutaneously into the flanks of tumor-free recipient hosts. Tumors derived from the CTC-organoids displayed similar histological features as their parental tumors, including expression of key markers such as YFP, Ki67, AR, luminal cytokeratin (Ck8), and vimentin (Fig. 3B, Fig. S4E-I). To evaluate metastatic potential, CTC-organoid lines from *NPK* early-stage and *NPK* end-stage mice were implanted intracardially into tumor-free recipient hosts (Fig. 3E, F). Both gross examination and histological analyses revealed that host mice implanted with NPK end-stage, not early-stage CTC-organoids had a high propensity to develop metastases, including to lung, liver, and bone, as determined by gross examination, histological analyses, and analyses of metastasis score (Fig. 3E, F; Dataset 1F). Taken together, these results show that CTC-organoids faithfully recapitulate the molecular, tumor, and metastatic phenotypes of the GEMMs from which they originate.

### Individual organoids (*i*Organoids) display morphological and molecular heterogeneity

Although the *NPK* end-stage mice exhibit a high propensity to metastasize, only a subset of their tumor cells express the META-16 signature associated with metastasis (see Fig. 1G), suggesting that not all of their tumor cells have similar metastatic potential. Moreover, *NPK* end-stage mice display considerable variability in the extent of their metastatic phenotype, as reflected by the wide range of CTC numbers and metastasis scores observed across individual mice (Fig. 2C, D; Fig. S2E, F). Therefore, to investigate their potential heterogeneity, we generated clonally derived organoids from individual CTCs (hereafter referred to as *i*Organoids) and analyzed their morphological features and transcriptomic profiles (Fig. 4A).

**Figure 4.**
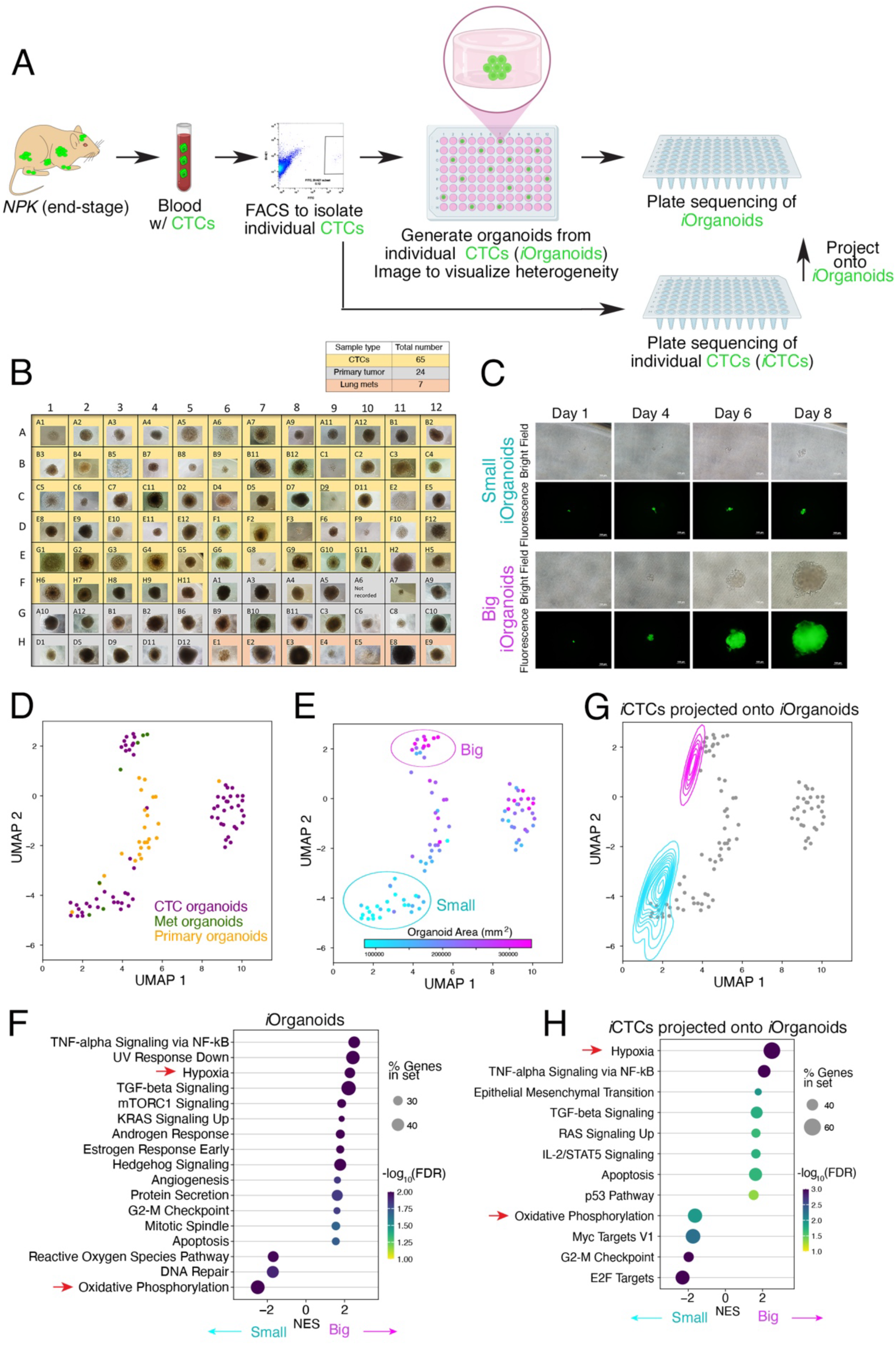
Individual organoids (*i*Organoids) display morphological and molecular heterogeneity. **(A)** Strategy for transcriptional profiling individual organoids (*i*Organoids) and individual CTC (*i*CTCs) from *NPK* end-stage mice. To generate *i*Organoids, individual CTCs were isolated via FACS and plated directly into a 96-well dish containing organoid culture media, grown for one week and then subjected to RNA sequencing (see *Methods*). Alternatively, individual CTC (*i*CTCs) were collected one cell per well onto a 96-well plate and directly subjected to scRNA sequencing. Controls for the *i*Organoids and *i*CTCs included individual cells from the primary tumor and/or lung metastases isolated by FACS. The numbers of *i*Organoids sequenced were: CTC n= 65; primary tumor n = 24; and lung metastatic n = 7. The numbers of *i*CTCs sequenced were: CTC n= 576; primary tumor n = 192. **(B)** Representative bright-field images of *i*Organoids following 1 week of growth in culture showing the 96-well plate design. **(C)** Bright-field and fluorescence images of *i*Organoids grown for the indicated time points (days). Scale bar, 100 μm **(D)** UMAP embedding of *i*Organoids color-coded by origin: CTCs, purple; primary tumor, yellow; or metastases, blue. **(E)** UMAP embedding of *i*Organoids color-coded by Organoid Area (mm^2^) displayed as a continuous variable. The circles indicate the relative positions of the “big” and “small” *i*Organoids. **(F)** Dot plot showing the enrichment of hallmark pathways differentially correlated with organoid size across *i*Organoids. **(G)** UMAP embedding of scRNA seq data from the *i*CTCs projected as colored contours onto the *i*Organoids (represented as gray dots). *i*CTCs projecting onto “big” *i*Organoids are shown as purple contours, while those projecting onto “small” *i*Organoids are shown in light blue. **(H)** Dot plot showing enrichment analysis of hallmark pathways differentially expressed between *i*CTCs projected onto small versus large *i*Organoids. In panels **F** and **H**, the red arrows highlight key pathways of interest. See also Figure S5, Table S5, Dataset 1B and 4.

Specifically, we performed FACS to isolate live YFP-marked cells from CTC samples of *NPK* end-stage mice or, for comparison, from dissociated primary tumors or lung metastases from these mice and plated individual cells (one per well) in 96-well plates containing organoid culture media as above (Fig. S5A). After one week of growth, visual inspection revealed striking heterogeneity in *i*Organoid morphology (Fig. 4B). Despite being plated at the same time and cultured under identical conditions, *i*Organoids displayed markedly variable sizes after 7 days (Fig. 4B). Differences between “Big” and “Small” *i*Organoids became increasingly apparent over time (Fig. 4C) and were more pronounced in *i*Organoids derived from CTCs than in those derived from primary tumors or lung metastases (Fig. S5B), despite high organoid-forming efficiencies across all sources (Fig. S5C).

After one week of growth, *i*Organoids were transferred to a second 96-well plate and subjected to low-input RNA-sequencing analysis using a plate-sequencing platform (Fig. 4A, D) (*57*). Transcriptomic analysis revealed distinct subpopulations among the CTC-derived *i*Organoids, most of which were transcriptionally distinct from the primary tumor-derived *i*Organoids, whereas a subset more closely resembled those derived from lung metastases (Fig. 4D).

Strikingly, we observed a strong concordance between the transcriptional phenotypes of CTC-derived *i*Organoids and their physical size. UMAP embedding revealed a clear separation between “Big” and “Small” *i*Organoids (Fig. 4E), which corresponded to distinct gene expression programs (Fig. S5D, Dataset 4A, 4B). In particular, prostate basal cell markers, such as *Cd44* (Rho = 0.77; FDR adjusted *P* value = 3.8X 10^-16^) and *Itga6* (Rho = 0.71; FDR adjusted *P* value = 1.89 X 10^-13^), were strongly correlated with “Big” *i*Organoids, whereas genes involved in mitochondrial metabolism and cellular respiration, such as *Slc25a39* (Rho = −0.71; FDR adjusted *P* value = 1.89 X 10^-13^) and *Coa3,* (Rho = −0.71; FDR adjusted *P* value = 1.86 X 10^-13^) were correlated with “Small” *i*Organoids (Fig. S5D).

Notably, “*i*Organoid size’’ was associated with distinct metabolic programs, wherein “Big” *i*Organoids were enriched for a hypoxia-associated expression program whereas “Small” *i*Organoids size were enriched for oxidative phosphorylation (Fig. 4F), paralleling the relationship observed between *NPK* end-stage and *NPK* early-stage tumors (see Fig. 1H). Therefore, we asked whether organoid lines derived from *NPK* early-stage and *NPK* end-stage mice would display similar morphological heterogeneity when grown as clonally-derived organoids. To address this, the established *NPK* end-stage and *NPK* early-stage organoid lines (see Fig. 3) were dissociated, subjected to FACS to isolate single cells, and plated individually (one cell per well) in 96-well plates containing organoid culture media. After one week of growth, *i*Organoids derived from *NPK* end-stage and *NPK* early-stage mice displayed distinct “Big” or “Small” phenotypes, respectively, which correlated with cell number per organoid rather than cell size (Fig. S5F-J).

We next asked whether the molecular heterogeneity observed in the CTC-derived *i*Organoids recapitulates that of CTCs themselves. To address this, individual CTCs (*i*CTCs) were isolated by FACS onto 96-well plates (1 CTC per well) and sequenced using a plate-sequencing platform (Fig 4A; Dataset 1B; Dataset 5; see *Methods*) (*57*). Projection of *i*CTC cell density onto the *i*Organoid data revealed that probability density peaks representing distinct *i*CTC subpopulations closely corresponded to the “Big” and “Small” *i*Organoids (Fig 4G). Pathway analysis of these two *i*CTC subsets mirrored observations from *i*Organoids; namely, enrichment for EMT, hypoxia, TNFα signaling, and TGFβ signaling were associated with *i*CTCs projecting onto “Big” *i*Organoids whereas oxidative phosphorylation was enriched in CTCs projecting onto “Small” *i*Organoids (Fig. 4F, H). Collectively, these findings indicate that the morphological and molecular heterogeneity of individual CTC-derived organoids reflect intrinsic heterogeneity among CTCs, potentially linked to differences in their metabolic programs.

### Molecular and morphological heterogeneity of *i*CTCs is associated with clinical outcome

Consistent with the transcriptomic analyses of the *i*Organoids, expression profiling of the *i*CTC scRNA-seq data revealed several distinct cell clusters (Fig. S6A), indicative of substantial underlying heterogeneity. To identify molecular drivers of this heterogeneity, we performed master regulator (MR) analyses using the *i*CTC scRNA-seq dataset, which was processed to minimize the effects of read depth and cell-cycle–associated variation (see *Methods*) (Fig. 5A–C). First, we applied the ARACNe algorithm (*58*) to infer genome-wide transcriptional regulatory networks from scRNA-seq data derived from mouse prostate tumor cells (as in Fig. 1). We then interrogated these inferred networks using MetaVIPER (*59*), the single cell extension of the VIPER algorithm (*60*), which infers protein activity by assessing the enrichment of each protein’s regulon within differentially expressed genes. This approach nominates candidate master regulators (MRs) as proteins whose inferred activities are most differentially distributed across cells. Using this framework, we identified four distinct *i*CTC MR modules (M_1_–M_4_), each corresponding to transcriptionally and functionally distinct CTC cell states (Fig. 5B,C; Fig. S6B; Dataset 5A–C). Notably, differentially active MR proteins in modules M_2_ and M_3_ were significantly enriched for the *i*Organoid size signature, with M_3_ exhibiting the strongest enrichment (*P* = 2.2 × 10⁻⁹; Fig. 5D).

**Figure 5.**
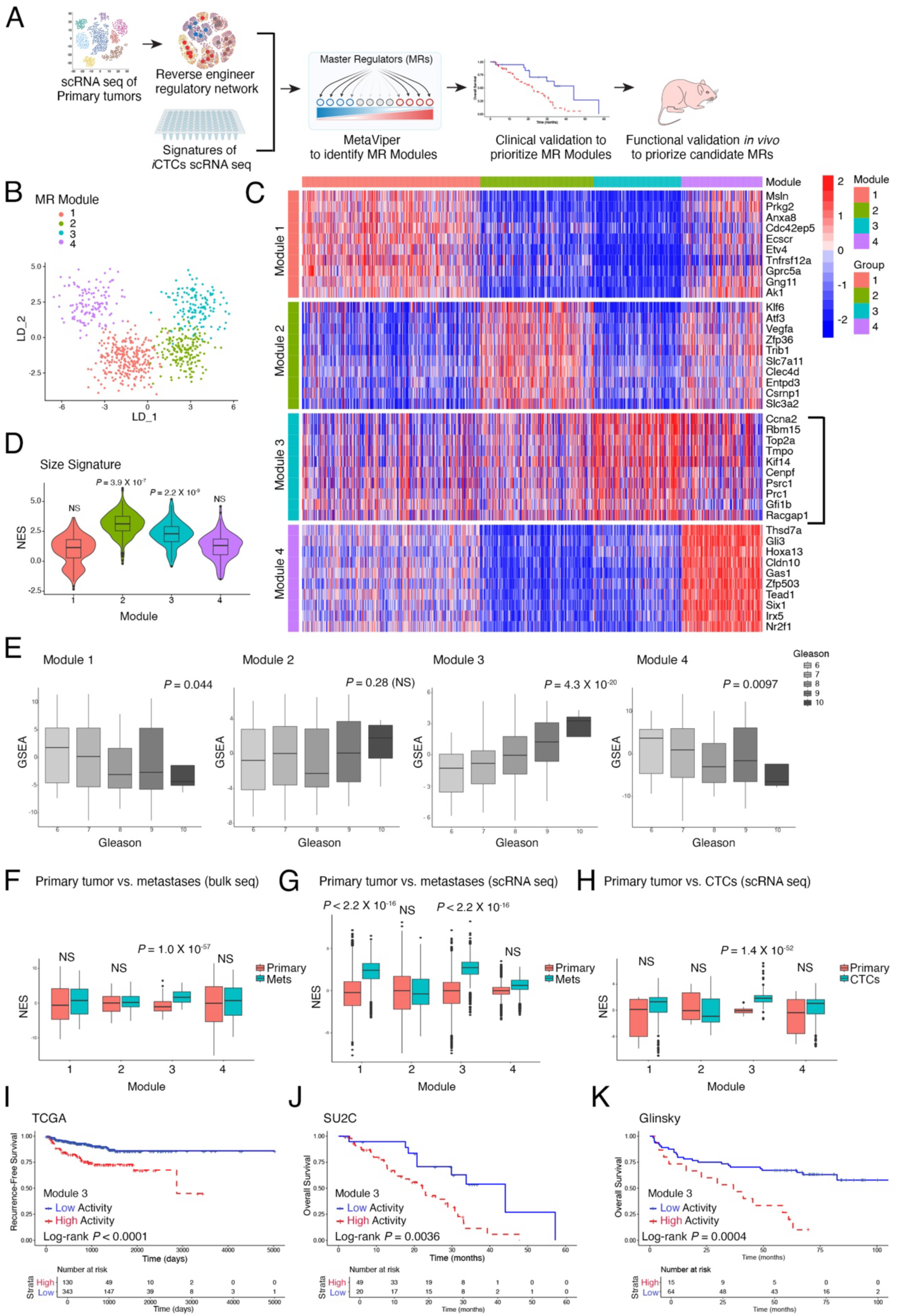
Molecular and morphological heterogeneity of *i*CTCs is associated with clinical outcome. **(A)** Strategy for identification of master regulator (MR) modules. The scRNA-seq data from the mouse primary tumors (as in Fig. 1) were used to reverse engineer whole genome regulatory networks. These networks were then interrogated with the scRNA-seq data from the *i*CTCs to transform differential gene expression signatures into differential protein activity profiles. This identified 4 distinct master regulator (MR) Modules (MR_1_-MR_4_). Prioritization of MR modules was done by correlative clinical validation using human patient cohorts to identify MR modules associated with prostate cancer progression and adverse disease outcome. Prioritization of individual candidate MRs within the MR modules was done by *in vivo* functional analyses (see *Methods*)**. (B)** Linear Discriminant (LD) projection plot showing *i*CTCs color-coded by the MRs-associated modules (MR modules MR_1_-MR_4_), as indicated. **(C)** Heatmap showing differential protein activity across the MR modules. Differential protein activity was computed by t-test, and top MRs for each module were ranked by statistical significance. **(D)** Violin plot showing the NES values for enrichment of the indicated MR modules (MR_1_-MR_4_) with the organoid size signature (see Fig. 4). NES values were determined by 1000 random permutations of gene ranking, with distribution of resulting values for each cell plotted by module and compared for each Module against entire dataset by t-test. **(E-K)** Prioritization of MR modules based on human patient cohorts. **(E)** Box plot showing the distribution of module enrichment scores plotted against Gleason score in the TCGA dataset. *P* values were computed using Pearson correlation of MR module enrichment score versus Pearson score, with Benjamini-Hochberg correction for multiple hypothesis testing. **(F-H)** Box plots showing NES for the indicated MR modules (MR_1_-MR_4_). Panel **F** compares bulk RNA seq data from primary tumors (n = 558, TCGA cohort) versus metastases (n = 270, SU2C cohort). Panel **G** compares scRNA-seq data from primary tumors (*n* = 7, NeoRED cohort) versus metastases (*n* = 8, PRIME-CUT cohort). **Panel H** compares scRNA seq data from primary tumors (*n* = 12) and individual CTCs (*n* = 77). In panels **F-H**, Statistical comparisons of enrichment scores were performed by t-test with Benjamini-Hochberg multiple testing correction. **(I-K)** Kaplan-Meier curves showing the association of MR Module MR_3_ with recurrence-free survival (TCGA cohort, **I**) and overall survival (SU2C cohort, **J**; Glinsky cohort, **K**). *P* values were calculated using a two-tailed log-rank test. See also Figure S6-8, Table S5, Datasets 5-7.

To assess conservation of the mouse-derived CTC MR modules with human prostate cancer, we examined their enrichment using diverse clinical criteria across multiple independent patient cohorts. Specifically, we evaluated: *(i)* association with prostate cancer aggressiveness, based on Gleason score (*4*); *(ii)* relative enrichment in advanced disease, comparing primary tumors versus metastases and primary tumors versus CTCs; and *(iii)* association with adverse clinical outcome, based on biochemical recurrence and overall survival. Toward this end, we leveraged publicly-available transcriptomic datasets including bulk-seq cohorts of primary tumors and metastases from: *(i)* The Cancer Genome Atlas (TCGA), consisting of primary prostate tumors of untreated patients (*n* = 558) (*42*); *(ii)* the Stand Up To Cancer (SU2C), comprised of metastatic biopsies from patients that had been heavily treated (*n* = 270) (*43*); and *(iii)* the Glinsky cohort, which includes primary prostate tumors obtained from radical prostatectomies in patients with recurrent and non-recurrent disease (*n* = 79) (*61*). Additionally, we queried scRNA-seq of primary tumors, metastases, and CTCs from *(iv)* the NeoRED-P cohort, comprised of primary tumors from patients with high risk localized prostate cancer (N = 7 patients; n = 232 tumor cells) (*62*); *(v)* the PRIME-CUT cohort, comprised of metastatic tumor biopsies, including pre-treatment biopsies (N = 8 patients; n = 783 tumor cells) (*63*)); and *(vi)* a cohort of CTCs isolated from patients with metastatic prostate cancer (N = 13 patients; n = 77 CTCs), and primary tumors from 12 patients (*64*), combined with a second cohort of CTCs (N = 5 patients; n = 44 CTCs) (*65*). These cohorts are described in Table S5.

Among the four MR modules, M_3,_ which was the module most significantly associated with the size signature (see Fig. 5D), was also most significantly associated with prostate cancer outcome across these multiple clinical criteria and independent patient cohorts (Fig. 5E-K; Fig. S6C-G). Specifically, M_3_ activity was strongly associated with high Gleason score in the TCGA cohort (*P* = 4.3 X 10^-20^; Fig. 5E). Additionally, M_3_ activity was highly enriched in metastatic biopsies relative to primary tumors, as evident by comparing the SU2C versus TCGA cohorts (*P* = 1.0 X 10^-57^), as well as the PRIME-CUT versus NeoRED-P cohorts (*P* < 2.2 X 10^-16^; Fig. 5F, G). Notably, M_3_ activity was also highly enriched in CTCs compared to primary tumors (*P* = 1.4 X 10^-52^; Fig. 5H).

Among the MR modules, M_3_ activity was also most significantly associated with adverse clinical outcome, including biochemical recurrence in the TCGA cohort (log-rank *P* value < 0.0001) and overall survival in both the SU2C cohort (log-rank *P* value = 0.0036) and Glinsky cohort (log-rank *P* value = 0.0004) (Fig. 5I-K). The prognostic significance of M_3_ was further evident by multivariate analyses based on both the TCGA and SU2C cohorts, wherein M_3_ activity was significantly associated with poor prognosis, independent of Gleason score for primary tumors in the TCGA cohort (TCGA: hazard ratio 1.4, *P* < 0.001) and even improved prognosis relative to Gleason alone for metastatic biopsies in the SU2C cohort (hazard ratio 1.5, *P* = 0.005; Fig. S6F, G). Interestingly, M_4_, which was associated with the “Small” size organoids, was enriched in low Gleason score tumors (Fig. 5E) and tended toward inverse association with survival, although not statistically significant (Fig. S6C-E), suggesting it may be associated with favorable outcome. Taken together, these findings suggest that morphological and molecular heterogeneity of CTCs reflect biologically relevant tumor cell states that are predictive of human prostate cancer progression and clinical outcome.

Given the association of the M_3_ MR module with advanced prostate cancer and adverse clinical outcome, we next investigated the functional contribution of top candidate MRs within this module for dissemination and metastasis *in vivo* (Figs. S7, S8). We first established independent 2D cell lines from non-clonal NPK end-stage CTC organoid lines (as in Fig. 3) and confirmed that these 2D derivatives faithfully recapitulate the tumorigenic and metastatic phenotypes of their parental 3D organoid lines (Fig. S7A–G; Dataset 6A). We then transduced the 2D cell lines with lentiviral shRNAs targeting each of the top 10 candidate MRs in M_3_ and assessed the effects on colony formation *in vitro*, as well as tumorigenicity, cellular dissemination, and metastasis *in vivo* (Fig. S8A–J; Dataset 6B; Dataset 7). Although MR silencing minimally affected cell viability and primary tumorigenicity (Fig. S8B–E), depletion of several candidate MRs—including *Prc1*, *Hsp90b1*, and *Tmpo*—resulted in a reduced incidence of CTCs (Fig. S8G). Notably, silencing of *Tmpo* also led to a marked reduction in metastasis (Fig. S8F). Based on these findings, we focused subsequent analyses on *Tmpo*, also known as lamina-associated protein 2 or Lap2 (*66*).

### TMPO expression associated with adverse disease outcome

We first validated *Tmpo* expression in the mouse prostate cancer models. Analysis of *Tmpo* expression in the scRNA-seq data from GEMM primary tumors (as in Fig. 1) revealed high expression in a subset of vimentin-positive cells within NPK end-stage tumors, which notably overlapped with the META16 metastasis signature (Fig. 6A; see Fig. 1G). Consistent with these findings, we detected TMPO protein expression in CTCs from *NPK* end-stage mice (Fig. 6B; Fig. S9A).

**Figure 6.**
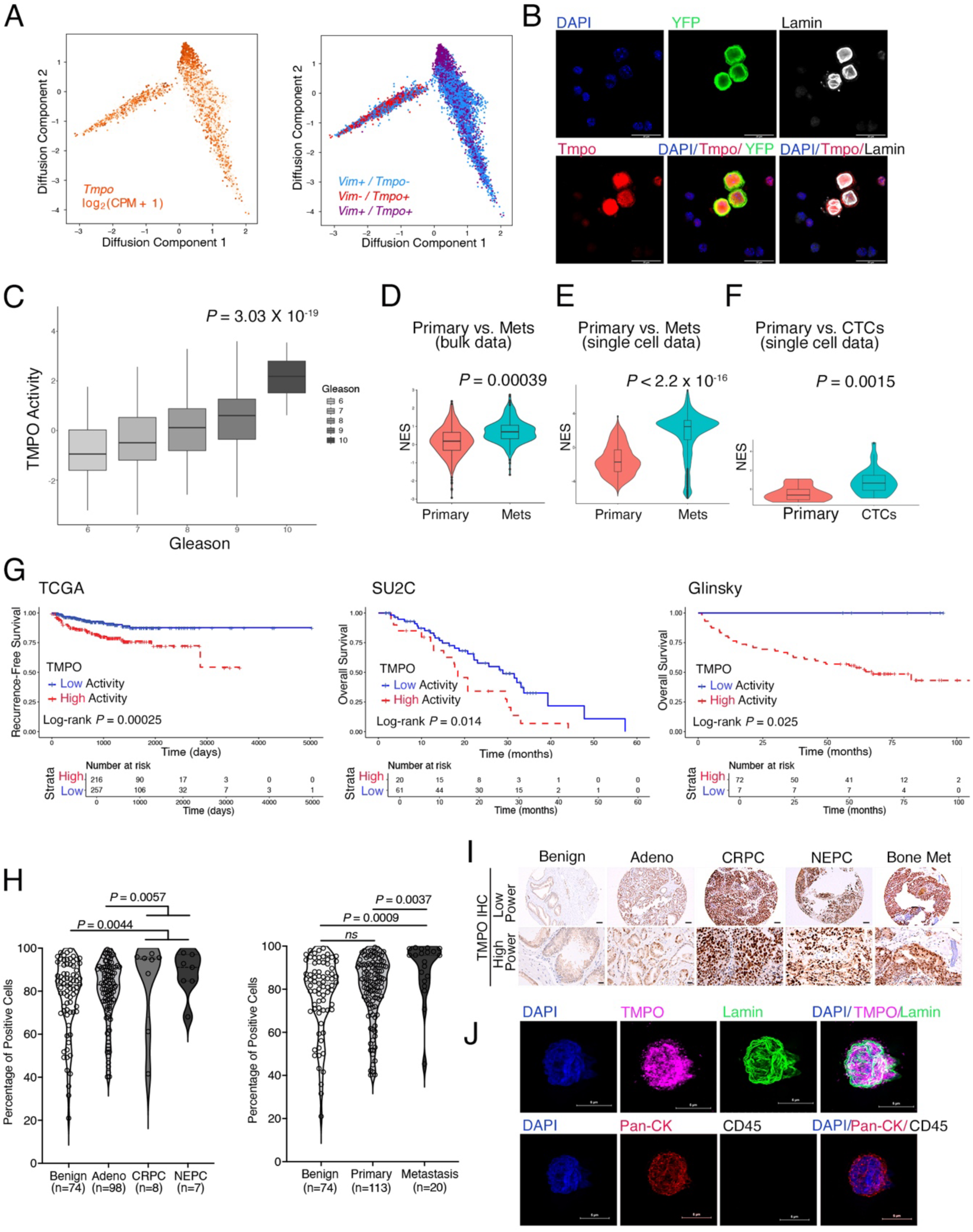
TMPO expression associated with adverse disease outcome. **<u>Panels A and B: Analyses of mouse prostate tumors and CTCs</u>: (A)** Diffusion Component (DC) projection of scRNA-seq data from mouse prostate tumors (see Figure 1) showing expression levels of *Tmpo* alone (left) or *Tmpo* together with *Vimentin* (right). **(B)** Immunofluorescence staining of CTCs obtained from *NPK* end-stage mice. DAPI; YFP, Tmpo and Lamin (Lamin A/C). Scale bars, 20 μm. **<u>Panels C-G: Transcriptomic analyses of human prostate cancer patient cohorts</u>: (C)** Box plot showing the distribution of TMPO activity depicted by enrichment scores plotted against Gleason score in the TCGA dataset. *P* values were computed by Pearson correlation of TMPO activity NES versus Pearson score. **(D-F)** Violin plots depicting distribution of the NES (y axis), which reflect activity levels of TMPO comparing: in panel **D**, bulk seq data of primary tumors from TCGA (*n* = 558) versus metastases from SU2C cohort (*n* = 270); in panel **E**, scRNA-seq data from primary tumors (*n* = 7, NeoRED cohort) versus metastases (*n* = 8, PRIME-CUT cohort); and in panel **F,** scRNA-seq data from primary tumors (*n* = 12) versus CTCs (*n* = 77) from the Myamoto *et al.* dataset. In panels **D-F**, *P* values for each comparison of enrichment scores were computed by Wilcoxon rank-sum test, with Benjamini-Hochberg multiple-testing correction. **(G)** Kaplan-Meier survival curves showing the association of TMPO activity with Recurrence-free survival (TCGA cohort, left), and Overall survival (SU2C and Glinsky cohorts, middle and right, respectively). *P* values were calculated using a two-tailed log-rank test. **<u>Panels H-J: Protein expression analysis of human prostate cancer patient cohorts</u>: (H-I)** Immunostaining of TMPO on tissue microarrays (TMA) of human prostate tumors and metastases. Analysis of TMPO expression was performed on digitized whole-slide images using HALO and HALO AI module (see *Methods*). **(H)** (Left) Violin plot showing the percentage of TMPO positive cells across tumor categories: benign prostate (*n* = 74), adenocarcinoma (Adeno, *n* = 98), castration-resistant prostate cancer (CRPC, *n* = 8), neuroendocrine prostate cancer (NEPC, *n* = 7). (Right) Violin plot showing the percentage of TMPO positive cells across benign tissues (*n* = 74), primary tumors (*n* = 113), and metastases (*n* = 20). For comparisons between more than two groups, *P* values were calculated using the Kruskal-Wallis test with Dunn’s multiple comparison correction. **(I)** Representative images from the TMAs in panel **H** showing TMPO immunostaining of benign prostate, adenocarcinoma (Adeno), CRPC, NEPC, and bone metastases (Bone met). Low power, scale bar: 100 μm; high power, scale bar: 20 μm). **(J)** Representative high-resolution confocal images of pan-CK^+^ CTCs enriched from patients with mCRPC and co-stained for Lamin A/C (Lamin) and TMPO expression (top). CTCs were identified as DAPI⁺/pan-CK⁺/CD45⁻ cells (bottom). Scale bar; 5 μm See also Figure S9, Table S6, S7, Datasets 8 and 9.

Given the limited number of studies that have investigated TMPO expression in human cancer, especially in human prostate cancer (*67, 68*), we assessed TMPO expression and MR activity at the transcriptomic level across the human prostate cancer patient cohorts and associated clinical parameters, introduced in Figure 5. We found that *TMPO* MR activity and expression were significantly upregulated in high Gleason score tumors in the TCGA cohort (*P* = 3.03 X 10^-19^ and *P* = 6.3 X10^-19^, respectively; Fig. 6C; Fig. S9C), comparable to that of the M_3_ MR Module (see Fig. 5E). Moreover, *TMPO* activity was up-regulated in metastases relative to primary tumors when comparing the SU2C versus TCGA cohorts (*P* = 0.00039, Fig. 6D; Fig. S9B), and the PRIME-CUT versus NEO-RED cohorts (*P* < 2.2 X 10^-16^; Fig. 6E). *TMPO* activity was also up-regulated in CTCs versus primary tumor cells (*P* = 0.0015; Fig. 6F). Most notably, *TMPO* activity and, to a lesser extent expression, were also associated with adverse disease outcome, including biochemical recurrence in the TCGA cohort (log-Rank *P* = 0.00025), and overall survival in the SU2C (log-Rank *P* = 0.014) and Glinsky cohorts (log-Rank *P* = 0.025) (Fig. 6G; Fig S9D). Beyond prostate cancer, analysis of three independent TCGA cohorts revealed that *TMPO* expression was significantly increased with tumor stage in breast, lung, and colon cancers (Fig. S9E), further supporting the clinical relevance of *TMPO*.

Given these findings, we examined TMPO protein expression in human prostate cancer using a series of tissue microarrays (TMAs) representing a spectrum of disease stages, including benign prostate, prostate adenocarcinoma, castration-resistant prostate cancer (CRPC) and neuroendocrine prostate cancer (NEPC), and encompassing both primary tumors and metastases, including to bone (n=141 total patients and 207 samples, Fig. 6H,I: Fig. S9F; Table S6, Dataset 8) (*69–73*). We found that TMPO protein expression was significantly upregulated in the most aggressive forms of prostate cancer (*P* = 0.004) as well as in metastases compared with primary tumors (*P* = 0.003) (Fig. 6H, I; Fig, S9F).

We next examined TMPO expression in CTCs obtained from patients with metastatic castration-resistant prostate cancer (mCRPC), including individuals receiving active systemic therapy (n = 5; Table S7). CTCs were isolated from peripheral blood using a size-based microfluidic platform and collected onto slides for multiplex immunofluorescence staining; CTCs were defined as nucleated, pan-CK⁺ or PSMA⁺, CD45⁻ cells (see *Methods*; Figs. 6J, S9G). Using these criteria, CTCs were detected in 4 of the 5 patients analyzed (Dataset 9A). Slides were subsequently co-stained with antibodies against TMPO and Lamin A/C, a nuclear structural marker, and imaged by high-resolution confocal microscopy. Strikingly, TMPO expression was detected in nearly all (>99%) CTCs from these mCRPC patients (Fig. 6J; Datasets 9A, 9B). TMPO localized predominantly to the nucleus and co-localized with Lamin A/C. Collectively, these findings identify TMPO as a clinically relevant determinant of prostate cancer progression, dissemination, and metastasis, and of adverse patient outcome.

### TMPO promotes cellular dissemination and metastasis via metabolic reprogramming

Given the association of TMPO with disease progression and clinical outcome, we next examined its functional role in cellular dissemination and metastasis *in vivo*, as well as the relationship of these activities to molecular heterogeneity and metabolic reprogramming (Fig. 7A–O; Fig. S10A–L). We silenced *Tmpo*, along with two additional top candidate MRs from the M_3_ module (*Hsp90b1* and *Prc1*), or a control, in CTC-derived cell lines and assessed the effects on cellular dissemination and metastasis following subcutaneous or intracardiac transplantation (Fig. 7A–D; Fig. S10A–E; Datasets 6C and 6D). *Tmpo* silencing resulted in a significant reduction in both the incidence and number of circulating tumor cells (CTCs) (*P* = 0.0006; Fig. 7B; Fig. S10D; Dataset 6C), as well as a marked decrease in metastatic burden, including lung, liver, and bone metastases, as confirmed by *ex vivo* imaging and immunohistochemical analyses (*P* = 0.0064; Fig. 7C,D; Fig. S10B,E; Dataset 6C and 6D). These findings demonstrate that *Tmpo* is required for efficient cellular dissemination and metastasis *in vivo*.

**Figure 7.**
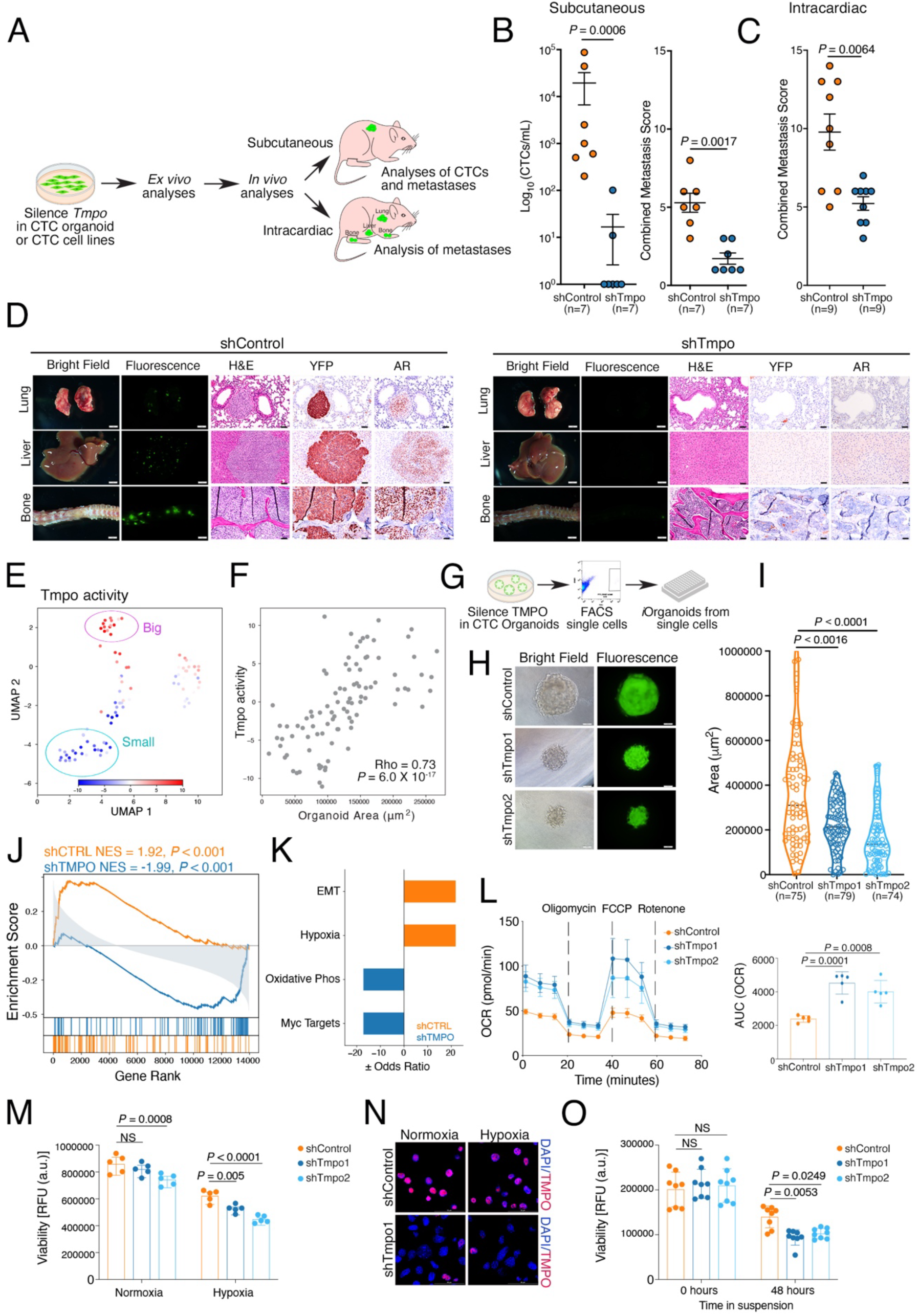
TMPO promotes cellular dissemination and metastasis via metabolic reprogramming. **(A)** <u>Overall Strategy</u>: Non-clonal CTC-derived cell lines were engineered with two independent doxycycline-inducible shRNAs targeting *Tmpo* or a control shRNA. The *Tmpo*-silenced or control CTC cell lines were analyzed *in vitro* (panels **E-O**) or grown *in vivo* following subcutaneous or intracardial injection (panels **B-D**). **(B-D)** Functional validation of Tmpo for dissemination and metastasis. **(B)** Subcutaneous engraftment of *Tmpo*-silenced or a control CTC cell lines (*n* = 7 for *shControl*; *n* = 7 for *shTmpo*). (Left) Scatter dot plot showing CTC counts per mouse reported on a logarithmic scale as CTCs/mL for the indicated groups. (Right) Dot plot showing the combined metastasis score for the indicated groups of mice. Data represent mean ± SEM. *P* values were calculated using a two-tailed Mann-Whitney U test. **(C, D)** Intracardiac engraftment of *Tmpo*-silenced or a control CTC cell lines (*n* = 9 for *shControl*; *n* = 9 for *shTmpo*). **C**. Dot plot showing the combined metastasis score for the indicated groups of mice. Data represent mean ± SEM. *P* values were calculated using a two-tailed Mann-Whitney U test. **(D)** Representative images of lung, liver and bone from recipient mice showing the bright-field and *ex vivo* fluorescence images H&E and IHC for YFP and AR, as indicated. Scale bars represent 2 mm for the whole mounts and 50 μm for the H&E and IHC. **(E-F)** <u>Association of Tmpo with organoid size</u>. **(E)** UMAP embedding of Tmpo MR activity (as described in Fig. 6) projected onto *i*Organoids. The circles indicate the relative positions of the “big” and “small” *i*Organoids (see Fig. 4). **(F)** Scatter dot plot depicting the relationship between Tmpo MR activity and organoid area (μm^2^). **(G-I)** <u>Analyses of *Tmpo i*Organoids</u>**. (G)** Strategy: Non-clonal *Tmpo-*silenced or control CTC-derived cell lines were subjected to FACS and seeded as individual cells in 96-well plate directly in organoid culture conditions to obtain *Tmpo-*silenced and control *i*Organoids. The resulting *i*Organoids were grown for seven days following which the size (area) was calculated for each individual *i*Organoid. (*i*Organoids analyzed: n = 75 control; n =79 *shTmpo#1;* n =74 *shTmpo#2*). **(A) (H)** Representative bright-field and fluorescence images of sh*Tmpo* and control *i*Organoids, as indicated. Scale bar, 100 μm**. (I)** Violin plot showing area measurements (μm^2^) of *i*Organoids from the indicated groups. *P* values were calculated using the Kruskal-Wallis test with Dunn’s multiple comparison correction relative to the shControl group. **(J)** GSEA comparing differentially expressed genes from scRNA-sequencing of the non-clonal *shControl* and *shTmpo* CTC-organoids (as in Panel A) with the organoid size signature (as in Fig. 4). NES and *P* values were estimated using 1,000 gene permutations. **(K)** Hallmark pathway enrichment analysis for the leading-edge genes obtained from the GSEA in panel J. **(L-O)** <u>Association of *Tmpo* with metabolic activity</u>. **(L)** (Left) Seahorse analysis of mitochondrial respiration *Tmpo*-silenced or a control CTC cell lines. For OCR measurements, basal-, ATP-linked, maximal, and reserve respiration rates were quantified, and values normalized to the total intracellular ATP level determined by luminescence. (Right) Bar graph with overlaid dots showing the area under the curve (AUC) analysis of the overall quantification of the total oxygen consumption rate (OCR) from the seahorse OCR assay. Statistical significance for each group comparison was assessed using an unpaired t-test against the control group. **(M)** Bar graph with overlaid dots showing cell viability measurements (RFU, relative fluorescence units) of *Tmpo*-silenced or a control CTC cell lines under normal or hypoxic conditions at 48 hours. *P* values were calculated using two-way ANOVA with Dunnett’s’ multiple comparison test against the control group. **(N)** Representative immunofluorescence images from *Tmpo*-silenced or a control CTC cell lines under normal or hypoxic conditions showing the indicated markers. Scale bars, 50 μm. **(O)** Bar graph with overlaid plots showing cell viability measurements of *Tmpo*-silenced or a control CTC cell lines following detachment-induced stress at the indicated time points (0 and 48 hours). *P* values were calculated using two-way ANOVA with Dunnett’s multiple comparison test against the control group. See also Figure S10, Datasets 4B, 10, 11A and 11B.

To investigate its association with heterogeneity, we examined *Tmpo* activity in the *i*Organoid expression profiling data (as in Fig. 4). We found that *Tmpo* activity was strongly enriched in the “Big” versus “Small” *i*Organoids (Fig. 7E), and that both its activity and expression levels were strongly correlated with organoid size (Rho = 0.73, *P* = 6.0 X 10^-17^ and Rho = 0.49, *P* = 4 X 10^-7^, respectively; Fig. 7F, Fig. S10F).

To directly ask whether *Tmpo* is associated with organoid size, we silenced *Tmpo* in non-clonal *NPK* end-stage CTC organoid lines (Fig. 7G-I; Fig. S10I,J). Following organoid dissociation, single cells were isolated from *Tmpo*-silenced or control organoids by FACS, followed by plating individual cells on 96-well plates (1 cell per well) in organoid media (as in Fig. 4). Although *Tmpo*-silenced and control *i*Organoids exhibited comparable organoid forming-efficiency (Fig S10J), *i*Organoids derived from control cells preferentially formed “Big” organoids whereas those derived from *Tmpo*-silenced cells preferentially formed “Small” orgnaoids (*P* < 0.0016 for sh*Tmpo1*; *P* < 0.0001 for sh*Tmpo2*) (Fig. 7G-I). These results indicate that *Tmpo* contributes to organoid size and associated morphological features associated with CTC heterogeneity.

To investigate the underlying molecular basis of this phenotype, we performed scRNA-seq on the non-clonal *Tmpo*-silenced versus control organoid lines. GSEA revealed significant overlap between genes differentially expressed in *Tmpo*-silenced versus control organoids and the organoid size signature (for *shControl* NES = 1.92, *P* < 0.001; for *shTmpo* NES = -1.99, *P* < 0.001; Fig. 7J; Dataset 10). Pathway analysis of leading-edge genes showed enrichment of hypoxia and EMT programs in control organoids, similar to the “Big” *i*Organoids, whereas *Tmpo*-silenced organoids were enriched for oxidative phosphorylation, similar to the “Small” *i*Organoids (Fig. 7K, and see Fig. 4F). Consistent with these observations, *Tmpo* silencing increased oxidative phosphorylation, as measured by oxygen consumption rate (OCR) (Fig. 7L; Dataset 11A), indicating that *Tmpo* regulates metabolic state as well as organoid morphology.

To further investigate these activities, we silenced *Tmpo* in the non-clonal 2D CTC-derived cells and examined the effects on cell viability under stress conditions encountered by CTCs in circulation, including hypoxia and mechanical stress (Fig. 7M-O). We found that *Tmpo* silencing significantly impaired CTC survival under hypoxic conditions and reduced viability following detachment-induced mechanical stress (Fig. 7M-O, Dataset 11B). In contrast, *Tmpo* silencing had no significant effect on either cell migration or invasion (Fig. S10G,H,K,L; Dataset 11C). Collectively, these findings support a model in which *TMPO* activity in CTCs promotes metabolic reprogramming that enhances CTC survival under conditions of cellular stress, thereby facilitating dissemination and metastasis.

## Discussion

Despite the fact that metastasis is the leading cause of cancer-related mortality, it is an extraordinarily inefficient process, since only a small fraction of tumor cells that escape the primary site are capable of surviving in circulation and ultimately seeding metastatic outgrowths. Understanding what distinguishes circulating tumor cells (CTCs) that survive from those that do not—and elucidating the mechanisms that confer a selective advantage for dissemination and metastasis—is therefore of paramount importance. By interrogating the molecular and morphological heterogeneity of individual CTCs (*i*CTCs) and organoids derived from individual CTCs (*i*Organoids) we identified key drivers of CTC heterogeneity that are associated with disease progression and adverse clinical outcomes in human prostate cancer. Most notably, we have identified TMPO as a key regulator of dissemination and metastasis and our analyses indicate that it functions by promoting survival of CTCs under conditions of metabolic and mechanical stress. Together, these findings provide new insights into the biological basis of CTC heterogeneity, particularly with respect to how a subset of CTCs acquires a selective advantage for survival under stress. These results may ultimately inform new strategies for prognostic stratification of tumors at high risk for metastasis, as well as identify potential therapeutic opportunities to prevent or limit tumor cell dissemination and metastatic progression.

*TMPO* encodes lamina-associated polypeptide 2 (LAP2), a nuclear membrane–associated protein containing a LAP2–Emerin–MAN1 (LEM) domain, for which several isoforms have been described (*66*). Although LAP2 has been extensively studied as a structural component of the nuclear envelope and for its interactions with other nuclear membrane proteins (*66, 74, 75*), significantly less is known about the expression or functions of TMPO in cancer, particularly in prostate cancer (*67*). In particular, TMPO has been implicated in chromatin organization and transcriptional regulation, influencing pathways associated with proliferation, survival, and migration by modulating the activity of several transcription factors and signaling pathways, including the cell-cycle regulator retinoblastoma protein (*76*), MRTF-A (*77*), and Gli1 (*74*). However, specific roles for TMPO in promoting metastasis have not been previously apparent.

Our data uncover a previously unrecognized link between TMPO and metabolic reprogramming during metastatic dissemination. TMPO silencing results in deregulation of oxidative phosphorylation, hypoxia-associated pathways, and epithelial–mesenchymal transition (EMT) programs, suggesting that TMPO promotes cell survival by coordinating multiple stress-adaptation mechanisms. This is consistent with the now widely appreciated concept that metabolic heterogeneity within tumors contributes to differences in metastatic efficiency, with highly metastatic cells relying on mechanisms that buffer oxidative stress (*22–24*). Thus, we propose a model in which TMPO becomes activated in lethal prostate cancer, and potentially in other cancers, and enables tumor cells that enter the circulation to overcome the hypoxic and mechanical stresses they encounter.

Our studies reveal a striking upregulation of *TMPO* in advanced prostate cancer, including in metastases and CTCs, and demonstrate a strong association with adverse clinical outcomes. Moreover, our functional studies establish a direct role for TMPO in promoting cellular dissemination and metastasis in prostate cancer models *in vivo.* Notably, these TMPO-associated functions may extend beyond prostate cancer, as we have also observed increased TMPO expression at advanced stages of tumor progression in breast, lung, and colon cancers. Our correlative findings based on human patient cohorts suggests that TMPO may be a *bona fide* marker of cellular dissemination in primary tumors, as well as a biomarker that distinguishes a subpopulation of CTCs with enhanced metastatic potential. Further investigation will be required to fully elucidate how TMPO expression is regulated in these specific contexts in tumor progression, as well as to define the precise mechanistic relationships among TMPO activity, metabolic reprogramming, and metastatic competence.

## Author Contributions

Conceptualization, A.G., P.A.S. and C.A-S.; methodology, A.G., P.G, P.A.S. and C.A-S.; formal analysis, A.G., A.Z.O., S.P., C.P-V. G.N.F., A.Z., M.D.B., M.L., I.I.C.C.,, P.G., A.C. and P.A.S.; investigation, A.G, J.L., S.P., C.P-V., G.N.F, J.Y.K., M.S.D., J.S.N., F.P., C.L., R.K.V., J.M.A. S.A. F.N.D.A., M.Z. H.G.; data curation, A.G., A.Z.O., G.N.F., A.Z., M.D.B., A.C., H.G., S.M., I.I.C.C., P.G., P.A.S. and C. A-S.; resources, B. D. R., H.G., S.M., D.A.H., D.T.M., D.M.N., S.T.T., M.L., I.I.C.C., P.G., A.C., P.A.S. and C.A-S.; software, A.Z.O., G.N.F., A.Z., M. D.B., A.C. and P.A.S.; writing- original draft, A.G., A.Z.O., C.P-V., G.N.F., I.I.C.C., M.S., P.G., A.C., PA.S. and C. A-S.; writing- review & editing, A.G., A.Z.O., S.P., C.P-V, G.N.F., D.A.H., D.T.M., D.M.N., I.I.C.C., M.S., P.G., A.C., P.A.S., and C.A-S.; visualization, A.G., A.Z.O., P.A.S. and C.A-S; supervision, C. A-S.; funding acquisition, C.A-S.

All authors participated in writing and approved the manuscript.

## Acknowledgments

Some figure panels (as indicated) were created with BioRender.com using an institutional license sponsored by Columbia University’s VP&S Office for Research. These studies were supported by shared resources funded by the Herbert Irving Comprehensive Cancer Center Grant (P30CA013696): Molecular Pathology, Flow Cytometry, and the Sulzberger Columbia Genome Center core facilities.

This work was supported by grants R01 CA173481 and R01 CA183929 (to C.A.S.), P01 CA265768 (to C.A.S., M.M.S. and M.L.), U54CA274506 (to P.A.S.), R35CA197745 and S10OD032433 (to A.C.), R01 CA266704 (to P.G.), P50CA211024, P01CA265768, R01 CA259200, the Department of Defense grant HT9425-25-1-0385 (to M.L.); R01 CA259007 and R01 CA302774 (to D.T.M.); R01 CA129933 (to D.A.H.); U01 CA214297 and R01 CA255602 (to D.A.H., S.M.).

This work was supported by funding from the Prostate Cancer Foundation, 22CHAL05 (to M.L) and 23CHAL15 (to M.M.S and A.C.) Pershing Square Sohn Research Alliance (to I.I.C.C.); the TJ Martell Foundation for Cancer Research (to C.A.S. and M.M.S); the Howard Hughes Medical Institute (to D.A.H.); the Breast Cancer Research Foundation (to D.A.H.); and the National Foundation for Cancer Research (to D.A.H). A.G. was supported by a Prostate Cancer Foundation (PCF) Young Investigator Award; a U.S. Department of Defense Prostate Cancer Research Program Award (W81XWH-18-1-0193); and an International Cancer Research Fellowship Outgoing iCARE funded by the Associazione Italiana per la Ricerca sul Cancro (AIRC) and the Marie Curie Actions COFUND. J.L. was supported by a U.S. Department of Defense Prostate Cancer Research Program Early Investigator Award (W81XWH-22-1-0078), and a Prostate Cancer Foundation (PCF) Young Investigator Award. S.P. was supported by a Bladder Cancer Advocacy Network Young Investigator Award (1068480), a Herbert Irving Comprehensive Cancer Center (HICCC) postdoctoral Pilot Award, and a fellowship from the American Cancer Society (ACS) (PF23-1142485). C.P-V. was supported by Pangaea Oncology, Spain. G. N.F. was supported by a fellowship from the American-Italian Cancer Foundation (AICF), and by the Italian Ministry of University and Research—PON “Research and Innovation” (PON R&I) Actions IV.4 “PhD programs and research contracts on innovation topics.” F.P. was supported by a Department of Defense Early Investigator Research Award (W81XWH-22-1-0054) and an HICCC Precision Oncology System Biology (POSB) program Postdoctoral Pilot Award. J.M. Arriaga was supported by the Dean’s Precision Medicine Research Fellowship from the Irving Institute for Clinical and Translational Research at Columbia University Irving Medical Center (CUIMC) (UL1TR001873) and a Prostate Cancer Foundation Young Investigator Award.

## Index of all Supplementary Materials

**Resource Table R1**. List of all reagents and resources used in this study

**Resource Table R2.** Description of the relevant mouse alleles

**Resource Table R3.** List of antibodies used in this study

**Resource Table R4**. Description of shRNAs used in this study

**Resource Table R5.** List of primers used in this study

**Supplementary Datasets** (*provided separately*)

**Dataset 1. Detailed description of the phenotypic characterization of individual mice included in this study and summary analysis performed on each** [related to all Figures]

**1A.** Complete description of individual mice and all analyses [related to all Figures]

**1B.** Description of individual mice used for RNA sequencing [related to Fig. 1 and S1].

**1C.** Description of mice analyzed from transplantation studies [related to Figs. 2 and S2].

**1D.** Description of mice used to derive CTC organoids [related to Figs. 3 and S3]

**1E.** Incidence of CTC organoids from GEMMs [related to Figs. 3 and S4]

**1F.** Description of mice used for intracardiac allografts [related to Figure 3]

**Dataset 2. scRNA-seq of mouse primary tumors** [related to Figures 1 and S1]

**2A**. scHPF Gene Score Matrix from scRNA-seq of primary tumors

**2B.** Z-scored average cell scores for scHPF factors

**Dataset 3. scRNA-seq data for early and late CTC organoids** [related to Figure 3]

**3A.** Ranked gene list for GSEA from differential expression analysis of *NPK* early-stage and *NPK* end-stage organoids

**3B**. Gene sets for GSEA from differential expression analysis of *NPK* early-stage and *NPK* end-stage primary tumors

**3C**. Leading-edge gene signatures

**Dataset 4. RNA seq analyses of *i*Organoids** [related to Figure 4]

**4A.** scHPF Gene Score Matrix from scRNA-seq of *i*Organoids

**4B.** Ranked gene list from correlation between *i*Organoid gene expression and area (e.g., the size signature).

**Dataset 5. VIPER analysis of *i*CTCs** [related to Figure 5]

**5A.** CTC VIPER matrix

**5B**. Matrix Metadata

**5C.** CTC VIPER markers

**Dataset 6. In vivo functional validation of MRs** [related to Figs. 5,7, S7 and S8, S10]

**6A.** Subcutaneous implantation of CTC cell lines

**6B**. In vivo functional validation of all candidate MRs via subcutaneous injection

**6C.** In vivo functional validation of 3 top MRs via subcutaneous injection

**6D.** In vivo validation of Tmpo function via intracardiac injection

**Dataset 7. In vitro MRs functional validation** [related to Figure 5 and S8]

**7A.** Quantitative real-time PCR (qPCR) validation of MRs expression

**7B.** Colony-forming assays

**Dataset 8: Quantification of TMPO immunostaining in human prostate cancer TMAs**

[related to Figs. 6 and S9]

**Dataset 9. Enumeration and phenotype of human CTCs** [related to Figs. 6 and S9]

**9A.** Enumeration and phenotypic profiling of CTCs in patient samples

**9B**. Enumeration and phenotypic profiling of PSMA+ CTCs in patient samples

**Dataset 10. GSEA from DE analysis comparing shControl to shTMPO** [related to Figure 7]

**10A.** Gene sets for GSEA from differential expression comparing shControl to shTMPO

**10B.** Leading-edge signatures from GSEA

**Dataset 11. Functional analyses of *Tmpo*-silenced CTC lines in vitro** [related to Figs. 7 and S10]

**11A.** Oxygen Consumption Rate (OCR) analysis

**11B.** Cell viability analysis under hypoxia and mechanical stress

**11C.** Migration and invasion assay

## Supplementary Figures

**Figure S1:**
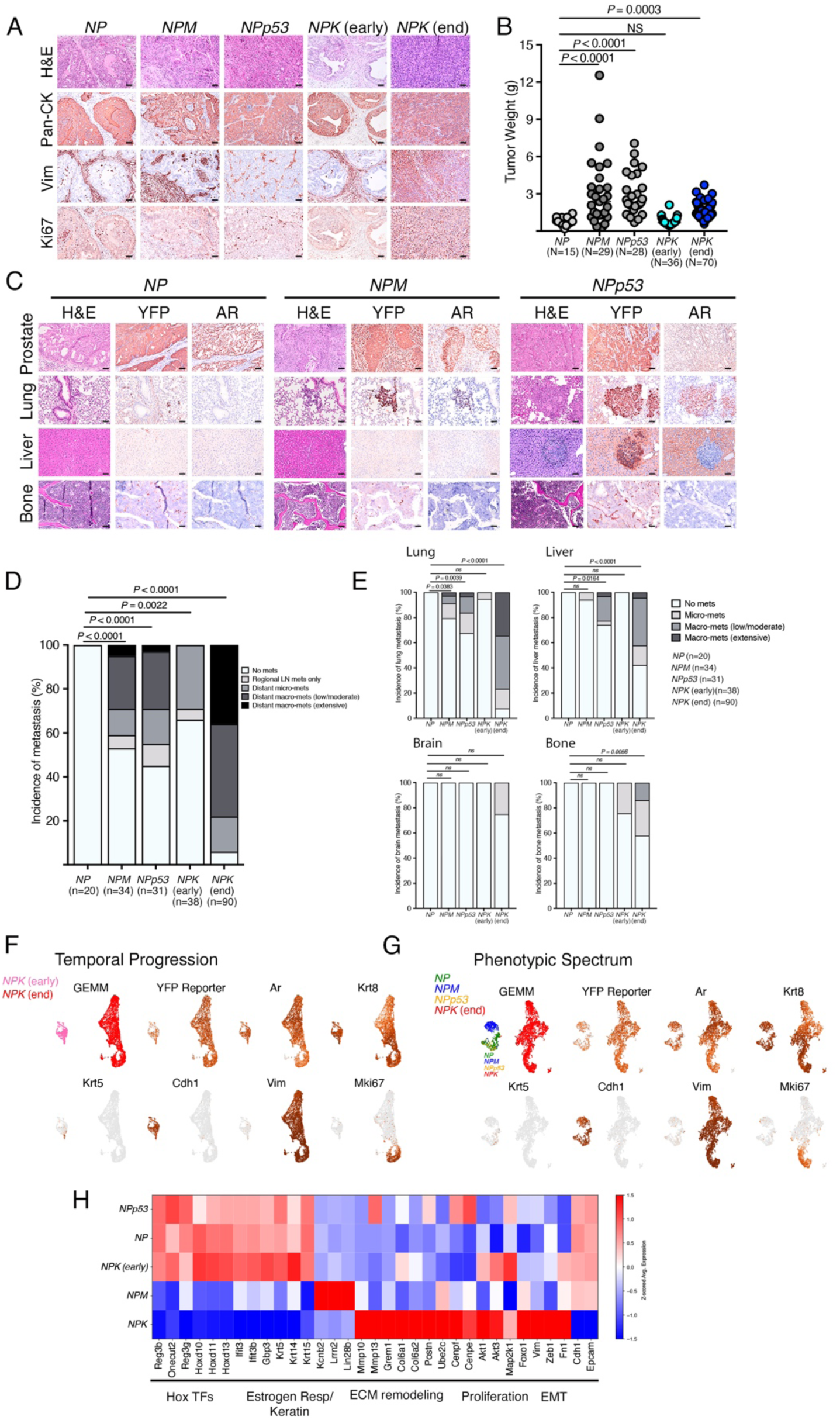
Additional analyses of the GEMMs [related to Figure 1]. **(A)** Representative images of prostate tumors of the indicated GEMMs showing histology (H&E) and immunostaining for Pan-CK (pan-cytokeratin), Vim (Vimentin), and Ki67, as indicated. (IHC was done on n = 3 mice per group) Scale bars, 50 μm **(B)** Dot plot showing tumor weight (g) for the indicated GEMMs. *P* values were calculated using the Kruskal-Wallis test with Dunn’s multiple comparison correction relative to control NP group. **(C)** Representative images of prostate tumors, and lung, liver, and bone, showing histology (H&E) and immunostaining for YFP, AR (Androgen Receptor), as indicated. (IHC was done on n = 3 mice per group) Scale bar, 50 μm. (**D, E)** Stacked bar graph showing the percentage of mice with metastasis across GEMMs. Panel **D** shows a composite across all sites, and panel **E** shows incidence of metastases to lung, liver, brain and bone, as indicated. (n = 20 for *NP;* n = 34 for *NPM;* n = 31 *for NPp53;* n = 38 for *NPK* early-stage; n = 90 for *NPK* end-stage mice). *P* values were calculated by performing pairwise comparisons against *NP* control group using the Fisher’s exact test. **(F, G)** Single-cell RNA-seq analysis of primary tumors from the indicated GEMMs. UMAP visualization of primary tumors cells from the indicated GEMMs. Shown are the scaled expression levels (DESeq2 normalized values) of YFP, Ar (Androgen receptor), Krt8 (Keratin 8), Krt5 (Keratin 5), Cdh1 (Cadherin 1), Vim (Vimentin), and Mki67. **(H)** Heat map showing Z-scores of average expression levels of the indicated genes across GEMMs.

**Figure S2:**
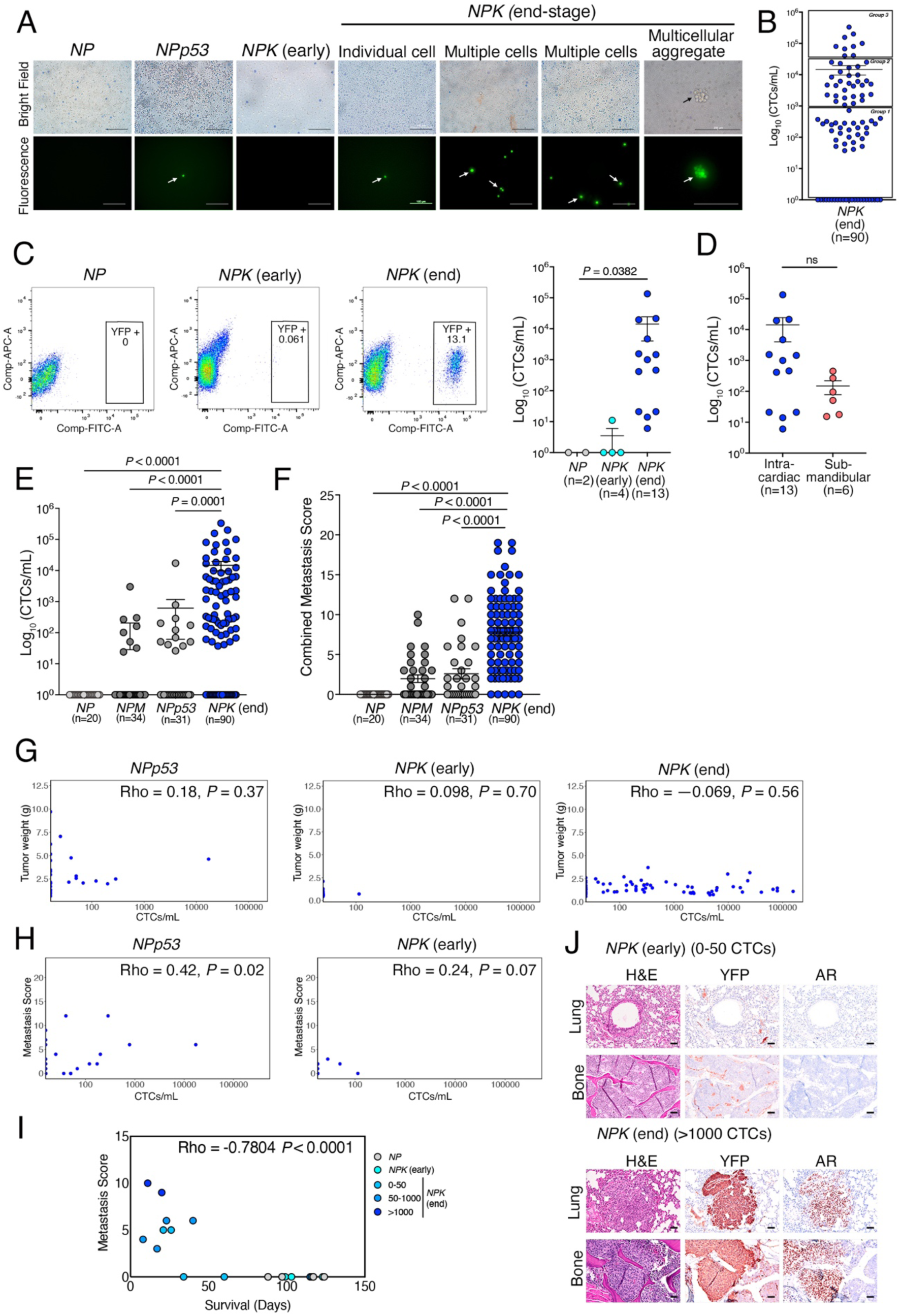
Additional characterization of CTCs from the GEMMs [related to Figure 2]. **(A)** Representative images of RBC-depleted blood samples from the indicated GEMMs stained with Trypan blue, showing live YFP-expressing CTCs visualized by fluorescence microscopy. Scale bar, 100 μm. **(B)** Scatter dot plot showing CTC counts per mouse (reported on a logarithmic scale as CTCs/mL) for the *NPK* end-stage mice. CTCs were quantified by visual counting of YFP-labeled cells under a microscope. Data represent mean ± SEM. **(C**) (Left) Fluorescence-activated cell sorting (FACS) of YFP-marked CTCs from blood samples collected from the indicated GEMMs. Representative FACS plots are shown, with the percentage of YFP-labeled cells indicated. Axes display fluorescent signals for the Fluorescein Isothiocyanate (FITC-A) and Allophycocyanin (APC-A) channels. (Right) Dot plot showing CTC counts (reported on a logarithmic scale as CTCs/mL) of FACS-analyzed CTCs for the indicated GEMMs. Data represent mean ± SEM. *P* values were calculated using the Kruskal-Wallis test with Dunn’s multiple comparison correction relative to control NP group. **(D)** Dot plot showing the CTC counts, reported on a logarithmic scale as CTCs/mL, obtained following FACS of CTC samples from *NPK* end-stage mice using blood from the indicated collection sites (intracardiac or submandibular). *P* values were calculated using the two-tailed Mann-Whitney U test. (**E)** Scatter dot plot showing the number of CTCs/per mouse reported on a logarithmic scale as CTCs/mL for the GEMMs as indicated. CTCs were quantified by counting live YFP-marked cells which were visualized by fluorescence microscopy. **(F)** Scatter dot plot showing the combined metastasis score for each mouse from the genotypes indicated. In panels **E** and **F**, data represent mean ± SEM. *P* values were calculated using the Kruskal-Wallis test with Dunn’s multiple comparison correction. **(G)** Scatter plot depicting the Spearman correlation between tumor weight (g) and CTC counts (CTCs/mL) for the indicated GEMMs. **(H)** Scatter plot depicting the relationship between metastasis scores and CTC counts (CTCs/mL) for the indicated GEMMs. In panel **G** and **H**, *P* values were calculated from a two-sided permutation test. **(I)** Scatter plot depicting the relationship between metastasis scores and survival (days) for the indicated groups. *P* values were calculated using Spearman correlation analysis. **(J)** Representative images of lung and bone from recipient hosts mice (from Fig. 2I) showing H&E and IHC for YFP and AR. Scale bars, 50 μm.

**Figure S3:**
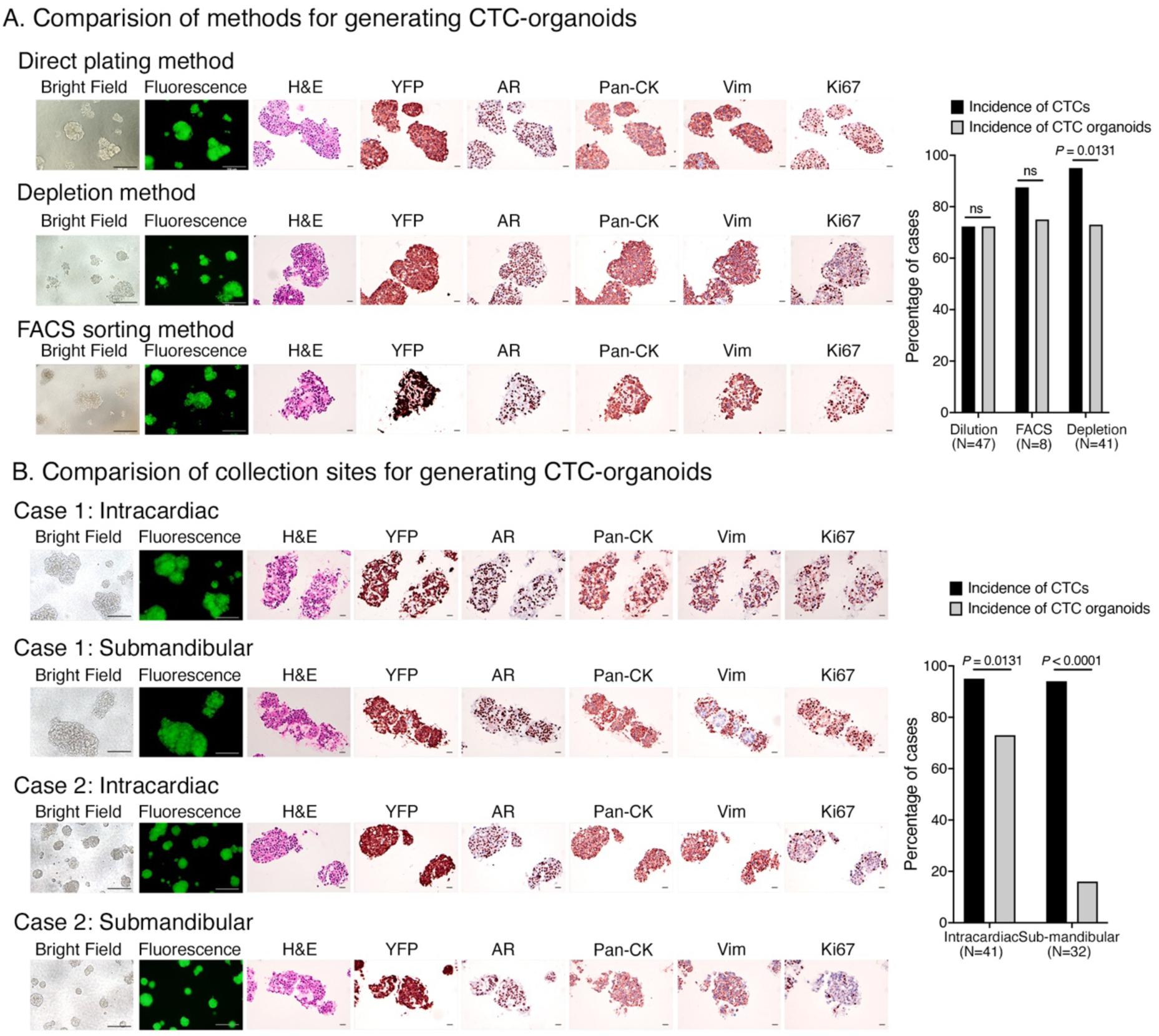
Methodology for generating CTC-derived organoids [related to Figure 3]. **(A** and **B)** (Left) Representative images from organoids generated by different methodologies (**A**) or the different blood collection sites (**B**). Panel **B** shows two independent matched cases. Shown are representative images of bright-field or fluorescence, H&E, or IHC for YFP, AR, pan-CK, Vim, and Ki67, as indicated. Scale bars for bright field and fluorescence images are 200 μm, and for the H&E and IHC images are 20 μm. (Right) Bar graph showing the incidence of CTCs and CTC organoids obtained by the indicated methodologies (**A**) or collection sites (**B**). *P* values were calculated using Fisher’s exact test.

**Figure S4:**
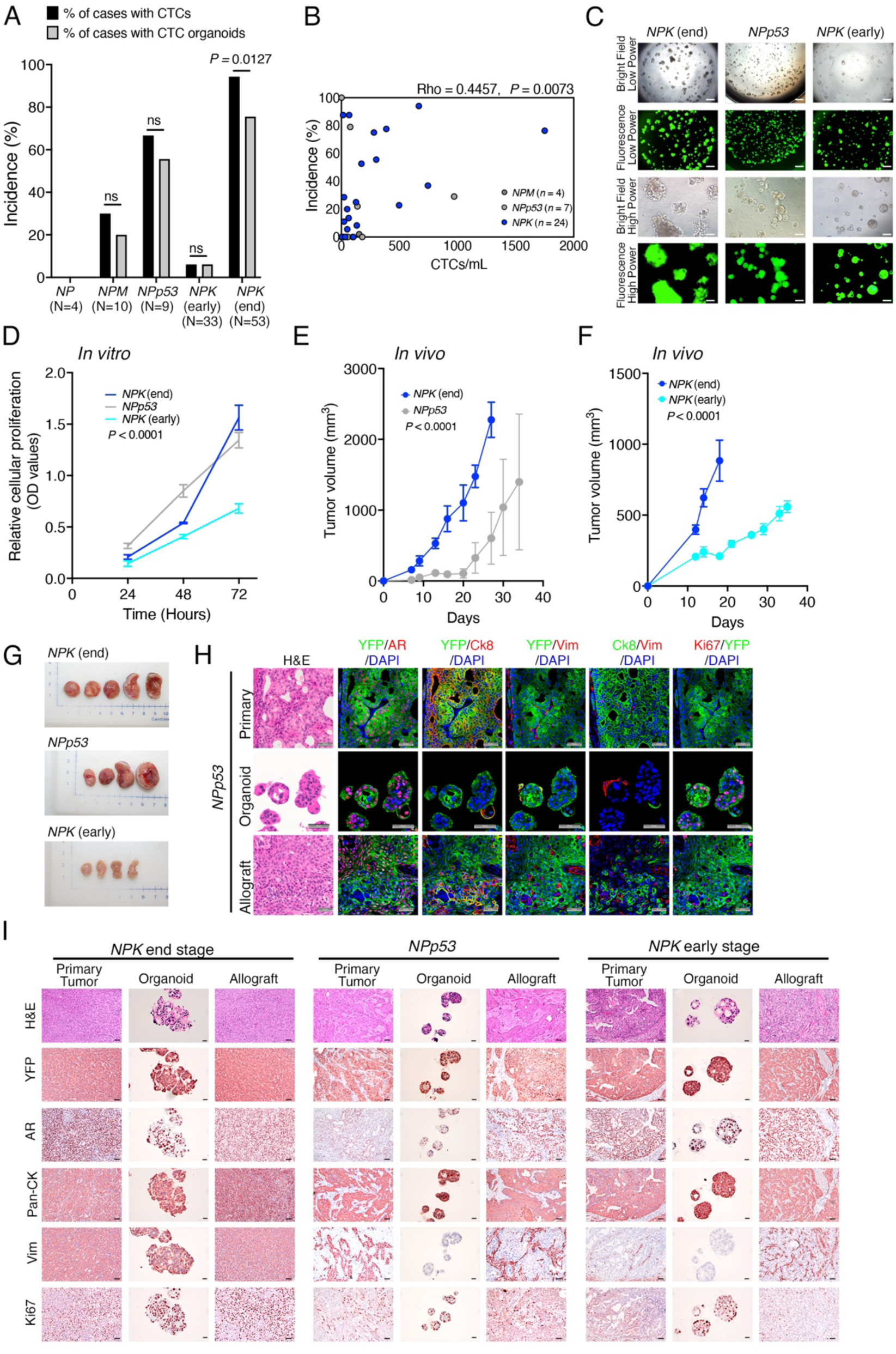
Additional phenotypic characterization of CTC-organoids [related to Figure 3]. **A)** Bar graph showing the incidence of CTCs and CTC-derived organoids in the indicated mice. (n = 4 for *NP;* n = 10 for *NPM;* n = 9 *for NPp53;* n = 33 for *NPK* early-stage; n = 53 for *NPK* end-stage mice) *P* values were calculated using Fisher’s exact test. **(B)** Scatter plot depicting the relationship between organoid incidence and CTC counts (CTCs/mL) for the indicated GEMMs. *P* values were calculated using Spearman correlation analysis. **(C)** Representative bright-field and fluorescence images of CTC-derived organoids from the indicated GEMMs. Scale bars 500 μm for low power and 200 μm for high power. (**D)** Growth curve showing cell proliferation quantified by the MTT assay (3-(4,5-dimethylthiazol-2-yl)-2,5-diphenyltetrazolium bromide) of the indicated CTC-organoid lines. Absorbance was measured at 560 nanometers at the indicated time points (hours). OD, optical density. *P* values were calculated using a two-way ANOVA test. (**E-F)** Tumor volume measurements for the indicated groups. Organoid lines from the indicated GEMMs were implanted into the flank of *nude* host mice, and tumor growth was monitored by caliper measurements using the formula [Volume = (width)^2^ x length/2]. *P* values between the indicated groups were calculated using a two-way ANOVA test. **(G)** Shown are images of allografts tumors from CTC organoids from the indicated mice that were grown in non-tumor-bearing host mice and imaged at the time of euthanasia. (**H, I)** Representative images of matched pairs from the parental tumor (primary) and the corresponding CTC-derived organoid lines propagated *in vitro* (organoid) or grown as allografts *in vivo* (allograft). **H**. Representative H&E or immunofluorescence staining for the indicated markers. (n = 5). Scale bar: 50 μm. **(I)** Representative H&E and IHC staining. Scale bar for tissue images, 50 μm. Scale bar for organoid images, 20 μm.

**Figure S5:**
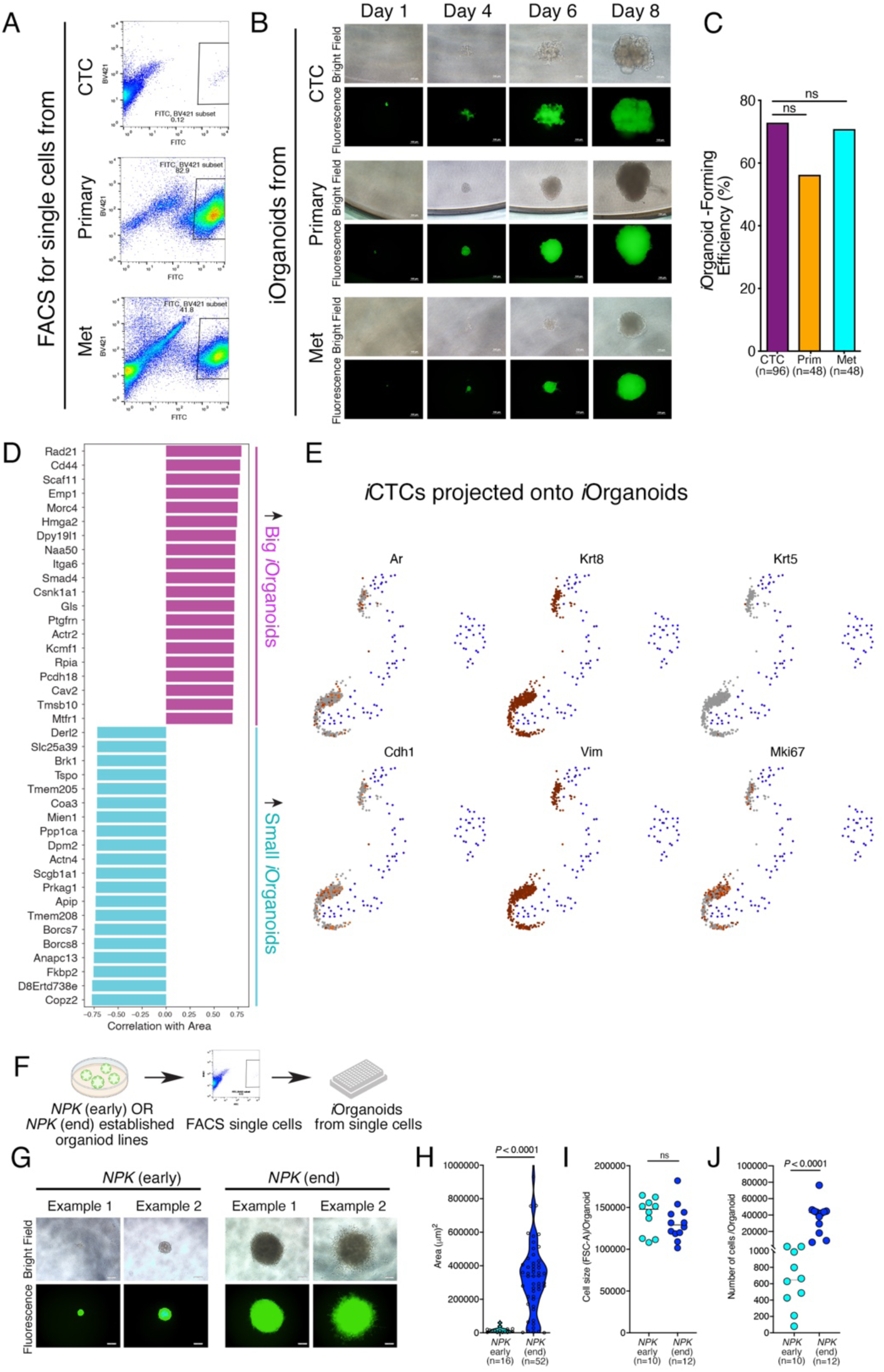
Additional characterization of *i*Organoids [related to Figure 4]. **(A)** FACS of YFP-marked CTCs from blood or dissociated tissue samples collected from *NPK* end-stage mice. Representative FACS dot plots are shown, with the percentage of YFP-labeled cells indicated. Axes display the fluorescent signal of the Fluorescein Isothiocyanate (FITC) and Brilliant Violet 421 (BV421) channels, used to detect YFP-marked live cells. **(B)** Representative bright-field and fluorescence images of individual organoids from CTCs (CTC), primary tumor (Primary) and lung metastases (Met), each grown from an individual sorted cell, shown at the indicated time points (days). Scale bar, 100 μm. **(C)** Bar graph showing the percentage of organoid-forming efficiency from the individual cells for the indicated groups. *P* values were calculated using Fisher’s exact test by performing pairwise comparisons against CTC group. **(D)** Bar chart showing genes correlated or anti-correlated with organoid area, ranked by Spearman correlation coefficient values. Genes positively correlated with organoid area are labeled in purple; negatively correlated genes are labeled in light blue. **(E)** UMAP embedding of scRNA seq data from the *i*CTCs projected tours onto the *i*Organoids (represented as gray dots) showing expression levels of the genes as indicated. **(F-J)** <u>Analyses of *i*Organoids from the established *NPK* early-stage and *NPK* end-stage organoid lines</u>**. (F)** Strategy: Non-clonal *NPK* early-stage or *NPK* end-stage organoid lines were subjected to FACS and seeded as individual cells in 96-well plate directly in organoid culture conditions to obtain *NPK* early-stage or *NPK* end-stage *i*Organoids. The resulting *i*Organoids were grown for seven days following which the size (area) was calculated for each individual *i*Organoid. **(G)** Representative bright-field and fluorescence images of *NPK* early-stage or *NPK* end-stage *i*Organoids, as indicated. Scale bar, 100 μm**. (H)** Violin plot showing area measurements (μm^2^) of *i*Organoids from the indicated groups. *P* values were calculated using a two-tailed Mann-Whitney U test. **(I-J)** (I) Dot plot showing cell size for the indicated groups. Cell size was measured using FSC-A values (forward scatter area). **(J)** Dot plot showing the number of cells per organoid for the indicated groups. *P* value was calculated using a two-tailed Mann-Whitney U test.

**Figure S6:**
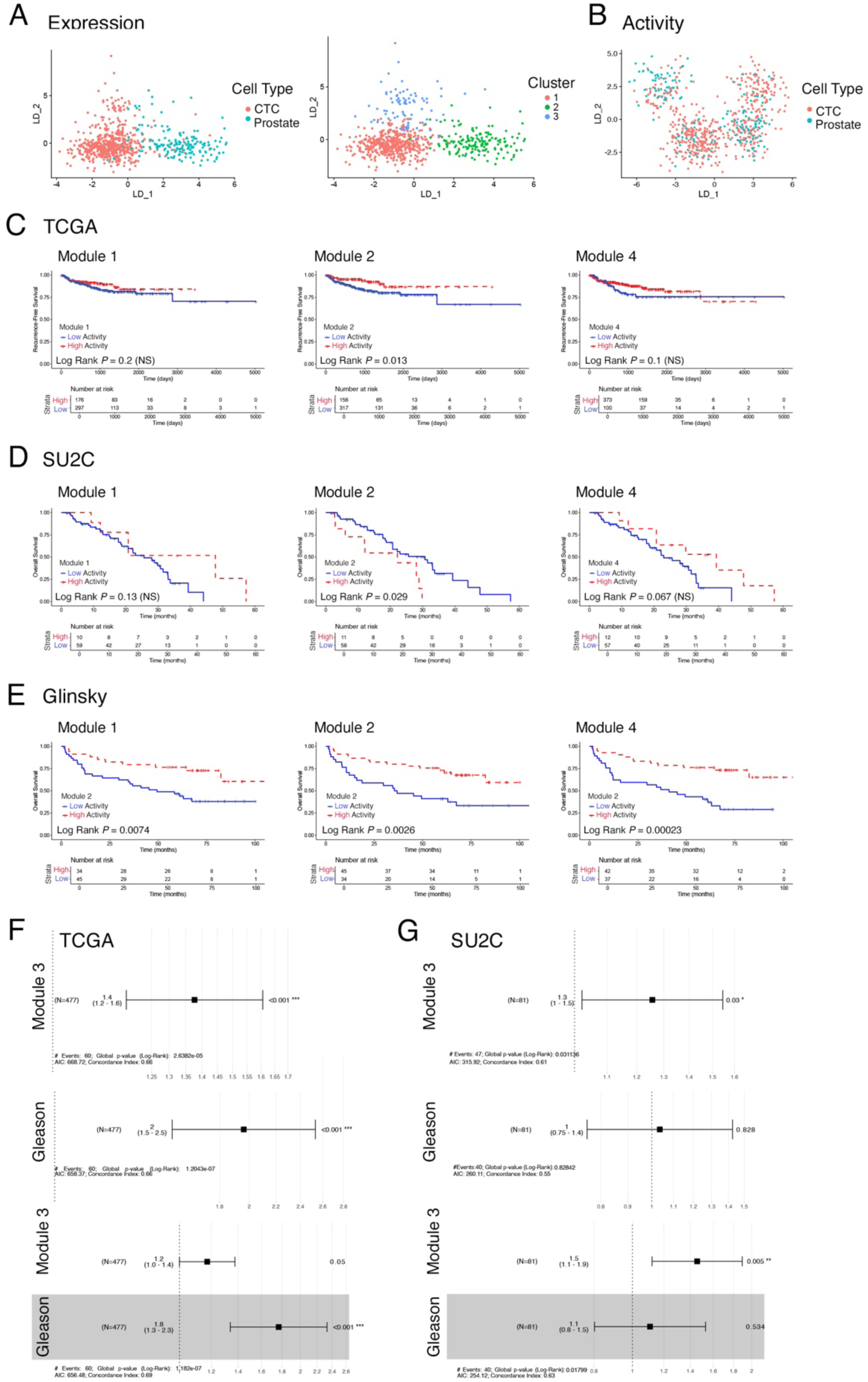
Additional clinical validation of MR modules [related to Figure 5]. **(A)** Linear Discriminant (LD) projection plot showing the *i*CTCs and *i*Primary tumor cells color-coded by cell type (Left) or cluster (Right) based on gene expression. **(B)** Linear Discriminant (LD) projection plot showing the *i*CTCs and *i*Primary tumor cells color-coded by cell type based on protein activity. **(C-E)** Kaplan-Meier curves showing the association of the indicated MRs’s Modules (M_1_, M_2_, or M_4_) with recurrence-free survival (TCGA in panel **C**) or overall survival (SU2C in panel **D** and Glinsky in panel **E**). *P* values were calculated using a two-tailed log-rank test. **(F, G)** Forest plots showing single-variable (MR Module 3 or Gleason), and multivariate Cox regression hazard ratios by testing the association of patient-by -patient normalized enrichment score for the MR Module 3 and Gleason Score with recurrent free survival in the TCGA **(F)** and overall survival in the SU2C **(G)** datasets.

**Figure S7:**
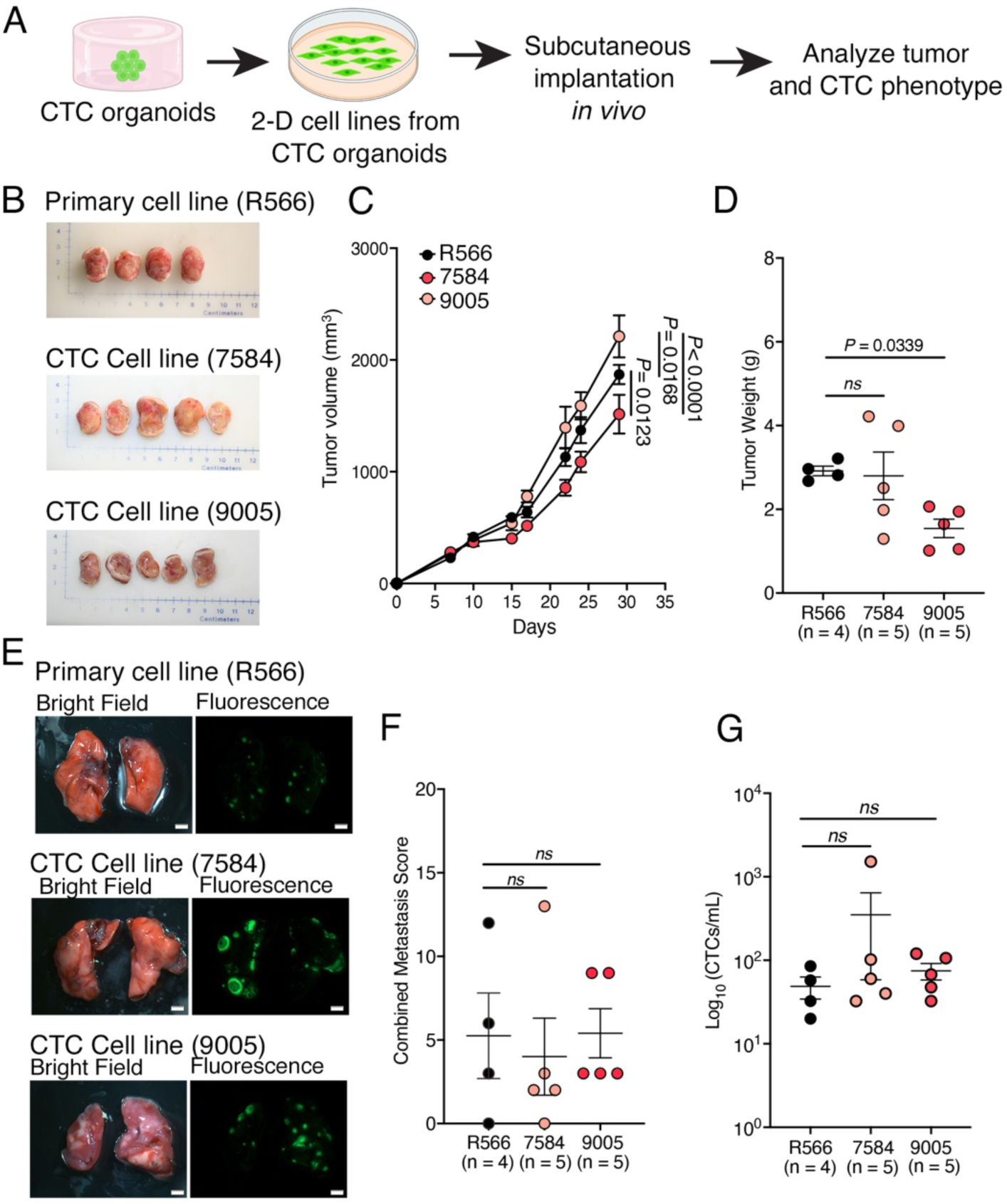
Establishment of 2D cell lines from CTC-organoids [related to Figure 5]. **(A)** Strategy to derive and validate 2-D cell lines from 3-D CTC organoid lines. Cell lines were derived from non-clonal established *NPK* end-stage CTC organoids. Once established the 2D lines were engrafted into recipient mouse hosts to assess the tumor, CTC incidence/enumeration, and metastatic phenotypes. n = 4 for control line R566; n = 5 mice for the CTC cell lines). We established 2 independent cell lines and performed studies to compare their activities with a previously established cell line from primary tumors reported in (*36*). CTC cell line 7584 and 9005 had similar phenotypes; for most subsequent analyses we used the 9005 cell line. **(B)** Shown are images of allografts tumors from the primary tumor-derived or CTC-organoid derived cell lines that were grown in non-tumor-bearing host mice and imaged at the time of euthanasia. **(C)** Tumor volume measurements for the indicated groups. Organoid lines from the indicated GEMMs were implanted into the flank of *nude* host mice, and tumor growth was monitored by caliper measurements using the formula [Volume = (width)^2^ x length/2]. *P* values between the indicated groups were calculated using a two-way ANOVA test. **(D)** Dot plot showing tumor weight (g) for the indicated groups. *P* values were calculated using the Kruskal-Wallis test with Dunn’s multiple comparison correction relative to the primary cell line (R566). Data represent mean ± SEM. **(E)** Representative bright-field and *ex vivo* fluorescence images of lungs from non-tumor bearing host mice engrafted with the indicated cell lines. Scale bar, 8 mm. **(F)** Dot plots showing the combined metastasis scores for the indicated groups. Data represent mean ± SEM*. P* values were calculated using the Kruskal-Wallis test with Dunn’s multiple comparison correction relative to the primary cell line (R566). **(G)** Dot plots showing the number of CTCs determined by visual counting under a microscope. CTC counts are reported on a logarithmic scale as CTCs/mL. *P* values were calculated using the Kruskal-Wallis test with Dunn’s multiple comparison correction relative to the primary cell line (R566). Data represent mean ± SEM.

**Figure S8:**
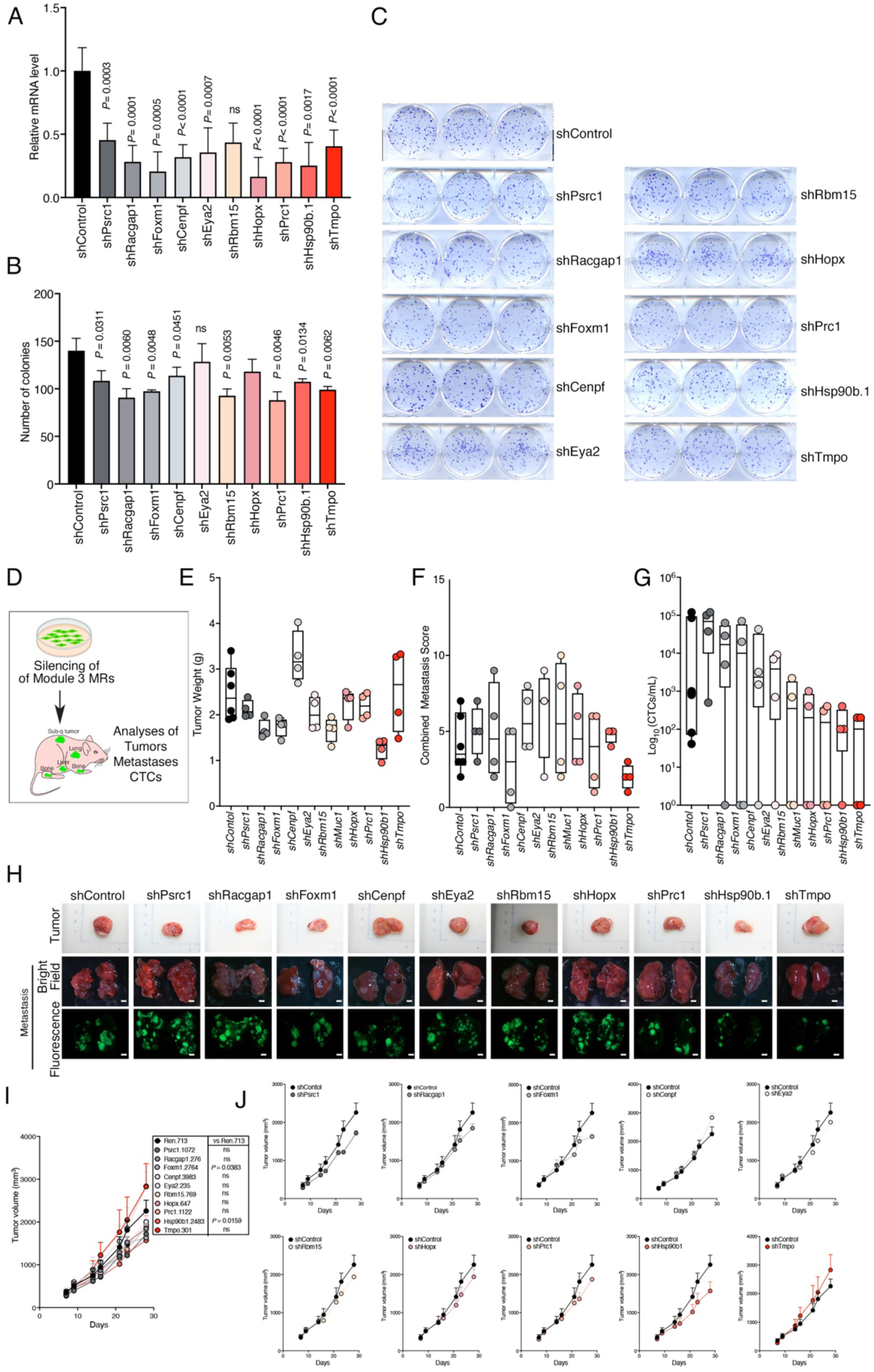
Functional validation of MRs from module M_3_ [related to Figure 5]. Functional analyses of top-ranked candidate MRs from Module 3 (see Fig. 5) was done by silencing the relevant genes using shRNAs as indicated in CTC cell line 9005 (see Fig. S7) and analyzed *in vitro* or *in vivo*. **(A-C)** <u>Analyses *in vitro*</u>. **(A)** Bar graph of quantitative real-time PCR analysis of the indicated genes showing relative expression levels represented as mean ± SD. *P* values were calculated using a two-tailed Student’s *t* test. **(B, C)** Colony forming assays. **(B)** Bar graph showing quantification of colonies. Data are represented as mean ± SD. *P* values were calculated using a two-tailed Student’s *t-* test. **(C)** Representative images showing crystal violet staining of colony formation assays. **(D-J)** <u>Analyses *in vivo*</u>. **(D)** MR-silenced or control CTC cells were grown in non-tumor bearing recipient mice and analyzed for tumor growth (**E, H-J**), metastatic potential (**F**), and CTC dissemination (**G**). (n = 6 for shControl group; n = 4 for each shRNA against the indicated MRs) **(E-G)** Box plots overlaid with individual data points showing tumor weight in grams (**E**), combined metastasis scores (**F**), and CTC counts, reported on a logarithmic scale as CTCs/mL (**G**)**. (H)** Representative images of tumors, and bright field and fluorescence images of lungs. Scale bar 8 mm (for *ex vivo* bright field and fluorescence images). **(I)** Tumor volume measurements of allograft tumors. *P* values between the indicated groups were calculated using a two-way ANOVA test relative to the control group. **(J)** Tumor volume measurements for each group with the indicated shRNA and the corresponding shControl group.

**Figure S9:**
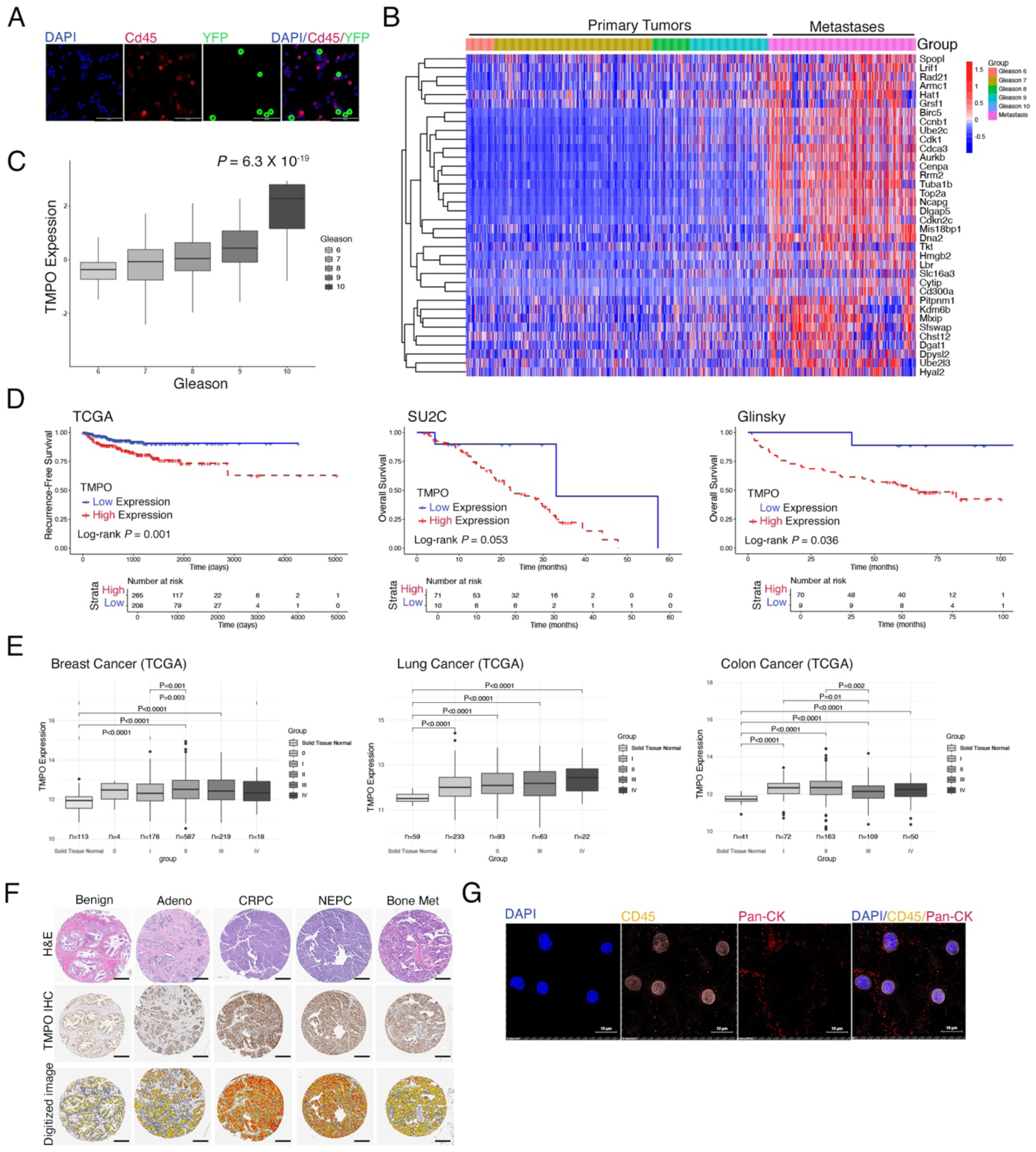
Additional validation of TMPO expression human cancer [related to Figure 6]. **(A)** Immunofluorescence staining of CTCs derived from *NPK* mice with the indicated markers. DAPI (4’,6-diamidino-2-phenylindole), Cd45 (Protein tyrosine phosphatase receptor type C, PTPRC), YFP. Scale bar, 50 μm. **(B)** Heatmap showing the expression of inferred upregulated target genes from the TMPO regulon in primary tumors (TCGA) and metastases (SU2C)**. (C)** Box plot showing the distribution of *TMPO* expression depicted by enrichment scores plotted against Gleason score in the TCGA dataset. *P* values were computed by Pearson correlation of *TMPO* expression normalized enrichment score versus Pearson score. **(D)** Kaplan-Meier survival curves showing the association of *TMPO* expression with Recurrence-free survival (TCGA cohort, left), Overall survival (SU2C, and Glinsky cohorts, middle and right, respectively). *P* values were calculated using a two-tailed log-rank test. **(E)** Box plots showing *TMPO* expression in Breast (Left), Lung (middle), and Colon (Right) cancer primary tumors from TCGA dataset. **(F)** Immunostaining of TMPO on tissue microarrays (TMA) comprised of human prostate tumors and metastases. Shown are H&E and immunostaining images of representative cores from TMAs, with digitalized images of TMPO staining for the indicated cases. Scale bar, 300 μm. **(G)** Representative images of immunofluorescent staining of peripheral blood mononuclear cells (PBMCs) from mCRPC. Cells were fixed and immune-stained for the indicated markers. PBMCs exhibited the expected hematopoietic profile (CD45⁺/pan-CK⁻). Scale bar: 10 μm.

**Figure S10:**
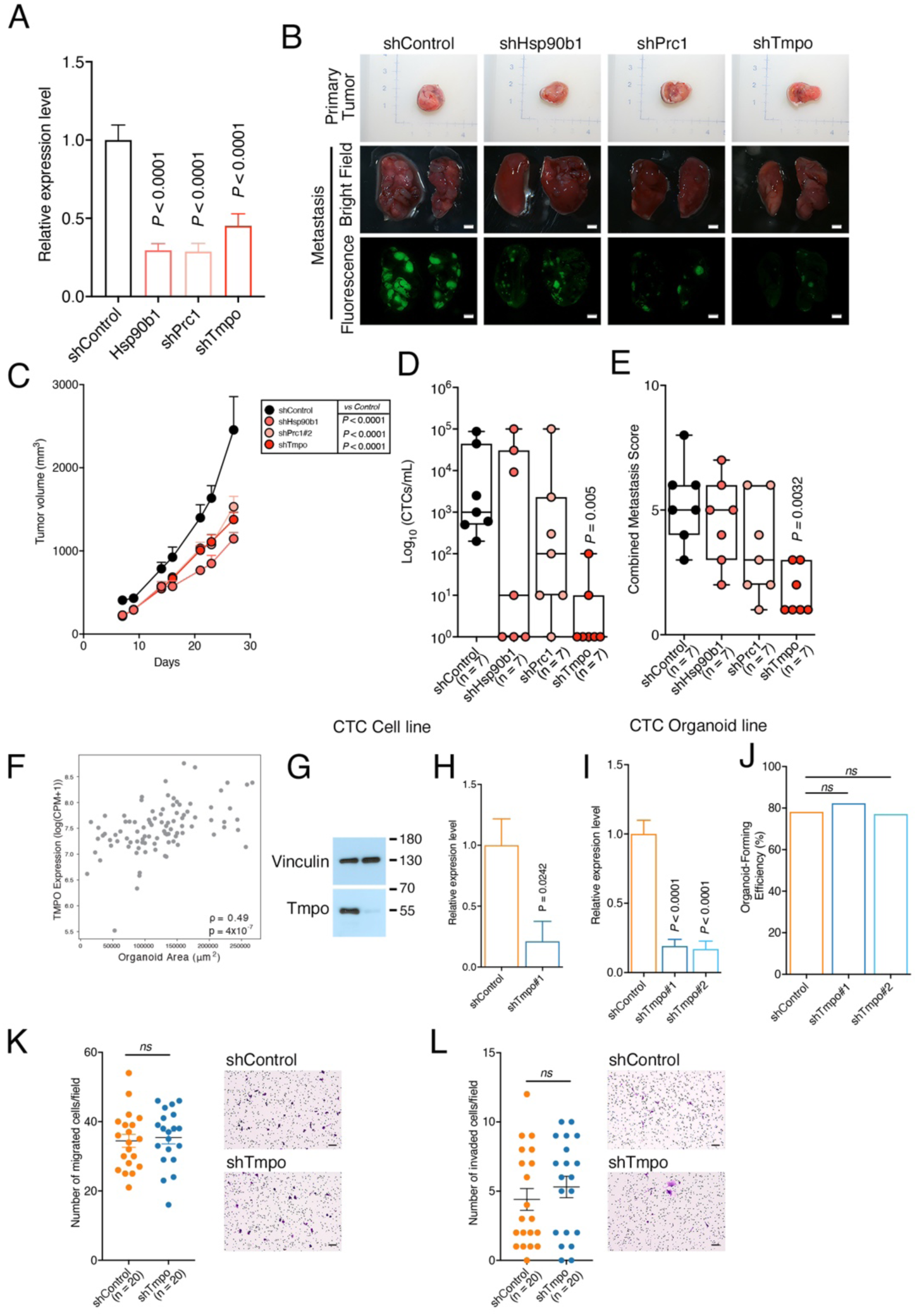
Additional functional validation of TMPO function [related to Figure 7]. **(A-E)** Analyses *in vivo*. (**A**) Bar graph showing quantitative real-time PCR analysis. Shown are relative expression levels for the indicated genes represented as mean ± SD. *P* values were calculated using a two-tailed Student’s *t* test. **(B)** Comparison of tumors and lungs collected at the time of euthanasia from nude mouse hosts implanted with CTC-derived cell line (as in Fig. S7) engineered with shRNAs targeting the indicated MRs. Representative images of tumors, and bright field and fluorescence images of lungs are shown. Scale bar 8 mm. **(C)** Tumor volume measurements for the indicated groups. Two-way ANOVA test was used to calculate the significance *(P* value) of the difference between the indicated groups relative to the control group. **(D, E)** Box plots with individual data points (n = 7 in each group) showing CTC counts (reported on a logarithmic scale as CTCs/mL) (D), and combined metastasis scores (E) from host mice implanted with the indicated MR-depleted CTC cell lines relative to shControl group. *P* values were calculated using the Kruskal-Wallis test with Dunn’s multiple comparison correction relative to the shControl group. (**F-L**) Analyses *in vitro.* **(F)** Scatter dot plot depicting the relationship between *Tmpo* expression and organoid area (μm^2^). **(G)** Western blot analyses of shControl or shTmpo-silenced cells for the indicated antibodies. Vinculin is a control for protein loading. **(H, I)** Bar graphs of quantitative real-time PCR analysis for CTC cell line (H) or organoid lines (I) engineered with shRNA targeting *Tmpo* or control shRNA. Shown are relative expression levels for the indicated gene represented as mean ± SD. *P* values were calculated using a two-tailed Student’s *t* test. **(J)** Bar graph showing the percentage of organoid-forming efficiency from the indicated groups. *P* values were calculated using Fisher’s exact test by performing pairwise comparisons against shControl group. **(K-L)** Migration and invasion assay. Dot plots showing the number of migrated cells per field (K) or invaded cells per filed (L) (n = 20) for the indicated groups. *P* values were calculated using a two-tailed Mann-Whitney U test. Shown are representative images of the migration and invasion assay for the indicated groups.

**Table S1.** Summary of the phenotypes of the GEMMs used in this study. [Related to Figures 1 and S1].

| GEMM | Total mice | Survival (days) |  | GU weight (g) |  | Metastasis Incidence |  |  |  |  |  |  |  |  |  |  |  | CTC incidence |  |  |  |  |  |
| --- | --- | --- | --- | --- | --- | --- | --- | --- | --- | --- | --- | --- | --- | --- | --- | --- | --- | --- | --- | --- | --- | --- | --- |
|  |  | Median Survival (days) | Log-rank test (relative to NP group) | Median GU weight (g) | IQR | Combined Metastasis Score |  | Overall |  |  | Lung |  | Liver |  | Brain |  | Bone |  | N | % | Mean CTC count/mL | Median CTC count/mL | IQR |
|  |  |  |  |  |  | Median Metastasis Score | IQR | N | % | Fisher's exact test (relative to NP group) | N | % | N | % | N | % | N | % |  |  |  |  |  |
| NP | 20 | >500 | / | 0.70 | 0.54 - 1.09 | 0 | 0 — 0 | 0 (20) | 0 | / | 0 (20) | 0 | 0 (20) | 0 | 0 (12) | 0 | 0 (12) | 0 | 0 (20) | 0 | 0 | 0 | 0 - 0 |
| NPM | 34 | 532 | P = 0.0322 | 2.37 | 1.21- 3.90 | 0 | 0 — 3 | 14 (34) | 41 | P = 0.0001 | 7 (34) | 21 | 2 (34) | 6 | 0 (23) | 0 | 0 (23) | 0 | 8(34) | 24 | 116 | 0 | 0 - 6 |
| NPp53 | 31 | 398 | P = 0.0025 | 2.25 | 1.99 - 4.23 | 1 | 0 — 4 | 14 (31) | 45 | P < 0.0001 | 10 (31) | 32 | 8 (31) | 26 | 0 (07) | 0 | 0 (07) | 0 | 12(31) | 39 | 613 | 0 | 0 - 50 |
| NPK (early) | 38 | NA | NS | 0.65 | 0.57 - 0.80 | 0 | 0 — 2 | 11 (38) | 29 | P = 0.0022 | 2 (38) | 5 | 0 (38) | 0 | 0 (37) | 0 | 9 (37) | 24 | 2(38) | 5 | 4 | 0 | 0 - 0 |
| NPK (end) | 90 | 156 | P < 0.0001 | 1.45 | 1.07 - 1.87 | 8 | 4 — 11 | 85 (90) | 94 | P < 0.0001 | 83 (90) | 92 | 52 (90) | 58 | 15 (60) | 25 | 24 (57) | 42 | 67(90) | 74 | 14638 | 283 | 0 - 5075 |

**Table S2A.** Summary of the incidence and enumeration of CTCs determined by direct counting [Related to Figures 2 and S2].

| GEMM | Total mice | GU Tumor weight |  | Incidence of urinary occlusion | Combined Metastasis Score |  | Blood collection site | Blood volume |  | Incidence of CTCs |  |  |  |  | Incidence of CTC clusters |  |
| --- | --- | --- | --- | --- | --- | --- | --- | --- | --- | --- | --- | --- | --- | --- | --- | --- |
|  |  | Median (g) | IQR |  | Median Metastasis Score | IQR |  | Median blood volume (μL)/sample | IQR | # of cases | % | Mean CTCs/mL | Median CTCs/mL | IQR | # of cases | % |
| Phenotypic spectrum of metastasis progression |  |  |  |  |  |  |  |  |  |  |  |  |  |  |  |  |
| NP | 20 | 0.70 | 0.54 - 1.09 | 0 (20) | 0 | 0 — 0 | Intracardiac | 350 | 300 - 488 | 0 (20) | 0 | 0 | 0 | 0 - 0 | 0 | 0 |
| NPM | 34 | 2.37 | 1.21- 3.90 | 1 (34) | 0 | 0 — 3 | Intracardiac | 475 | 288 - 525 | 8 (34) | 24 | 116 | 0 | 0 - 6 | 1 (34) | 2.9 |
| NPp53 | 31 | 2.25 | 1.99 - 4.23 | 5 (31) | 1 | 0 — 4 | Intracardiac | 500 | 400 - 700 | 12 (31) | 39 | 613 | 0 | 0 - 50 | 1 (31) | 3.2 |
| Temporal progression of metastasis |  |  |  |  |  |  |  |  |  |  |  |  |  |  |  |  |
| NPK (early) | 38 | 0.65 | 0.57 - 0.80 | 0 (38) | 0 | 0 — 2 | Intracardiac | 500 | 388 - 550 | 2 (38) | 5 | 4 | 0 | 0 - 0 | 1 (38) | 3.0 |
| Phenotypic heterogeneity within the NPK model |  |  |  |  |  |  |  |  |  |  |  |  |  |  |  |  |
| NPK (end) total | 90 | 1.45 | 1.07 - 1.87 | 29 (90) | 8 | 4 — 11 | Intracardiac | 300 | 200 - 400 | 67 (90) | 74 | 14638 | 283 | 0 - 5075 | 11 (90) | 12 |
| Group 1 (low) | 53 | 1.51 | 1.21 - 1.92 | 23 (53) | 6 | 3 — 9 | Intracardiac | 350 | 225 - 500 | 30 (53) | 57 | 122 | 50 | 0 - 225 | 0 (53) | 0.0 |
| Group 2 (medium) | 30 | 1.13 | 0.97 - 1.64 | 7 (30) | 10 | 7 —12 | Intracardiac | 250 | 138 - 400 | 30 (30) | 100 | 9809 | 5150 | 2228 - 13435 | 7 (30) | 23 |
| Group 3 (high) | 7 | 1.26 | 1.14 - 1.45 | 0 (07) | 12 | 10 — 15 | Intracardiac | 200 | 50 - 250 | 7 (07) | 100 | 145009 | 100000 | 80000 - 200000 | 4 (07) | 11 |

**Table S2B.**
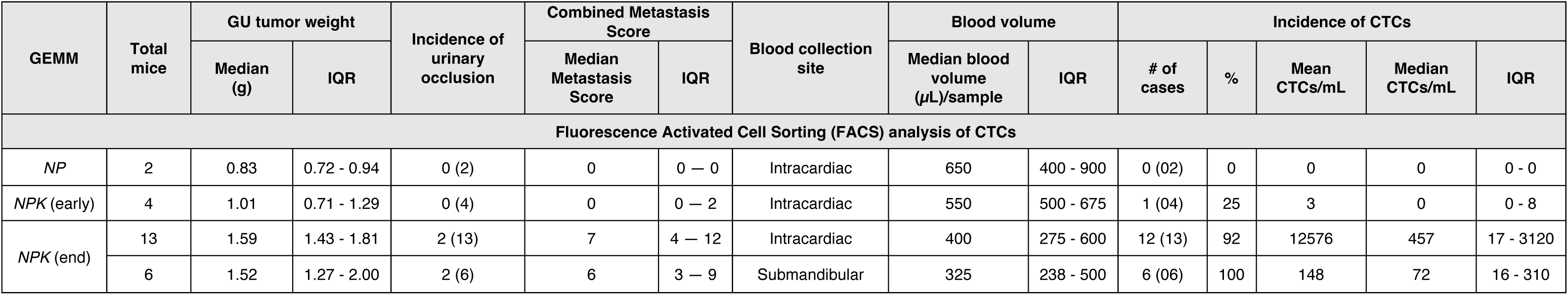
Summary of the incidence and enumeration of CTCs determined by FACS analysis. [Related to Figures 2 and S2].

**Table S3.** Summary of results from different methods used to establish CTC organoids. [Related to Figures 3 and S3].

| GEMM | Total samples analyzed | Blood collection site | Blood volume |  | Methodology | Incidence of CTCs |  | Incidence of CTC organoids |  |
| --- | --- | --- | --- | --- | --- | --- | --- | --- | --- |
| | | | Median blood (μL)/sample | IQR | | # of cases | % | # of cases | % |
| NPK | 47 | Intracardiac | 300 | 250 - 450 | Dilution | 34(47) | 72 | 34(47) | 72 |
|  | 8 | Intracardiac | 450 | 163 - 650 | FACS sorting | 7(8) | 88 | 6 (8) | 75 |
|  | 41 | Intracardiac | 300 | 190 - 475 | CD45 - negative depletion | 39(41) | 95 | 30(41) | 73 |
|  | 32 | Sub-mandibular | 200 | 100 - 338 |  | 30(32) | 94 | 5(32) | 16 |

**Table S4.** Summary of CTC organoid generation. [Related to Figures 3 and S4].

| GEMM | Incidence of CTCs |  |  |  | Incidence of organoids derived from CTCs |  |  | Organoid-forming efficiency |  |  |  |  |
| --- | --- | --- | --- | --- | --- | --- | --- | --- | --- | --- | --- | --- |
|  | Total samples | CTCs observed (Yes/No) | # of cases | % | Total samples | # of cases with CTC organoids | % | Total samples | # of cases with CTCs | Mean CTCs/mL | # of cases with CTC organoids | Mean organoid-forming efficiency (%) |
| *NP* | 4 | No | 0 (04) | 0 | 4 | 0 (04) | 0 | NA | NA | NA | NA | NA |
| *NPM* | 10 | Yes | 3 (10) | 30 | 10 | 2 (10) | 20 | 4 | 4 (4) | 59 | 1 (4) | 0 |
| *NPp53* | 9 | Yes | 6 (09) | 67 | 9 | 5 (09) | 56 | 7 | 7 (7) | 204 | 4 (7) | 33 |
| *NPK (early)* | 33 | Yes | 3 (33) | 9 | 33 | 2 (33) | 6 | NA | NA | NA | NA | NA |
| *NPK* | 53 | Yes | 50 (53) | 94 | 53 | 40 (53) | 75 | 24 | 24 (24) | 234 | 17(24) | 32 |

NA – not applicable

**Table S5.** Description of human transcriptomic datasets. [Related to Figures 5,6, S6 and S9].

| Ref/name | Description and use in this study | n | Platform | Geo/Link | Reference |
| --- | --- | --- | --- | --- | --- |
| TCGA cohort | <u>Description:</u><br>Surgical resection biospecimens from patients with prostate adenocarcinoma without prior treatment<br><u>Use in this study:</u><br>Discovery and validation of CTC MRs-associated modules with tumor progression and biochemical recurrence-free survival outcome. Validation of the expression and activity levels of TMPO in primary tumors | 558 | Illumina HiSeq 2000 W | [TCGA Data Portal: https://portal.gdc.cancer.gov/](https://portal.gdc.cancer.gov/) | (42) |
| SU2C cohort | <u>Description:</u><br>Bone or soft tissue tumor biopsies from post treatment metastatic castration-resistant prostate cancer (mCRPC). Only samples made with polyA libraries were used for mRNA expression analysis.<br><u>Use in this study:</u><br>Discovery and validation of the clinical relevance of the CTC MRs modules. Validation of the expression and activity levels of TMPO in metastases | 270 | Illumina HiSeq 2500 | [www.cbioportal.org GitHub portal: https://github.com/cBioPortal/datahub/tree/master/public/prad_su2c_2019](https://github.com/cBioPortal/datahub/tree/master/public/prad_su2c_2019) | (43) |
| Glinsky cohort | <u>Description:</u><br>Primary prostate tumors from radical prostatectomies from patients with recurrent and non-recurrent disease<br><u>Use in this study:</u><br>Discovery and validation of the clinical relevance of the CTC MRs-associated modules. Validation of the expression and activity levels of TMPO in metastases | 79 | Affymetrix Human Genome U133A | NA | (61) |
| CTC cohort 1 | <u>Description:</u><br>Bulk primary tumors from 13 patients with localized prostate cancer (n=12); Circulating tumor cells isolated with CTC-iChip from patients with metastatic prostate cancer (n = 77)<br><u>Use in this study:</u><br>Comparative analysis of CTC MRs- associated modules in CTCs versus primary tumors. Validation of TMPO protein activity in CTCs. | 77 CTCs<br>12 primary tumors | RNA sequencing of isolated CTCs<br>Bulk RNA sequencing of primary tumors | GEO: GSE67980 | (64) |
| CTC cohort 2 | <u>Description:</u><br>Circulating tumor cells isolated (n = 44) from 5 patients with metastatic castration-resistant prostate cancer (mCRPC) post treatment<br><u>Use in this study:</u><br>Comparative analysis of CTC MRs- associated modules in CTCs versus primary tumors (from CTC cohort 1) | 44 | HiSeq X | GEO:<br>GSE208449 | (65) |
| PRIME-CUT cohort | <u>Description:</u><br>Bone, lymph node lung, and liver tumor biopsies from patients with metastatic castration sensitive prostate cancer (mCSPC) at baseline from 8 patients (NCT03951831)<br><u>Use in this study:</u><br>Comparative analysis of CTC MRs-associated modules and TMPO protein activity in metastases (PRIME-CUT) versus primary tumors (NeoRED-P cohort) | 8 | Illumina<br>NovaSeq 6000 | Mendeley:<br>https://doi.org/10.17632/5nnw8xrh5m.1 | (63) |
| NeoRED-P cohort | <u>Description:</u><br>Surgical resection of treatment-naïve biospecimens (radical prostatectomies) from 7 patients with high risk localized prostate cancer<br><u>Use in this study:</u><br>Comparative analysis of CTC MRs-associated modules and TMPO protein activity in metastases (PRIME-CUT) versus primary tumors (NeoRED-P cohort) | 7 | Illumina<br>NovaSeq 6000 | NA | (62) |

**Table S6.**
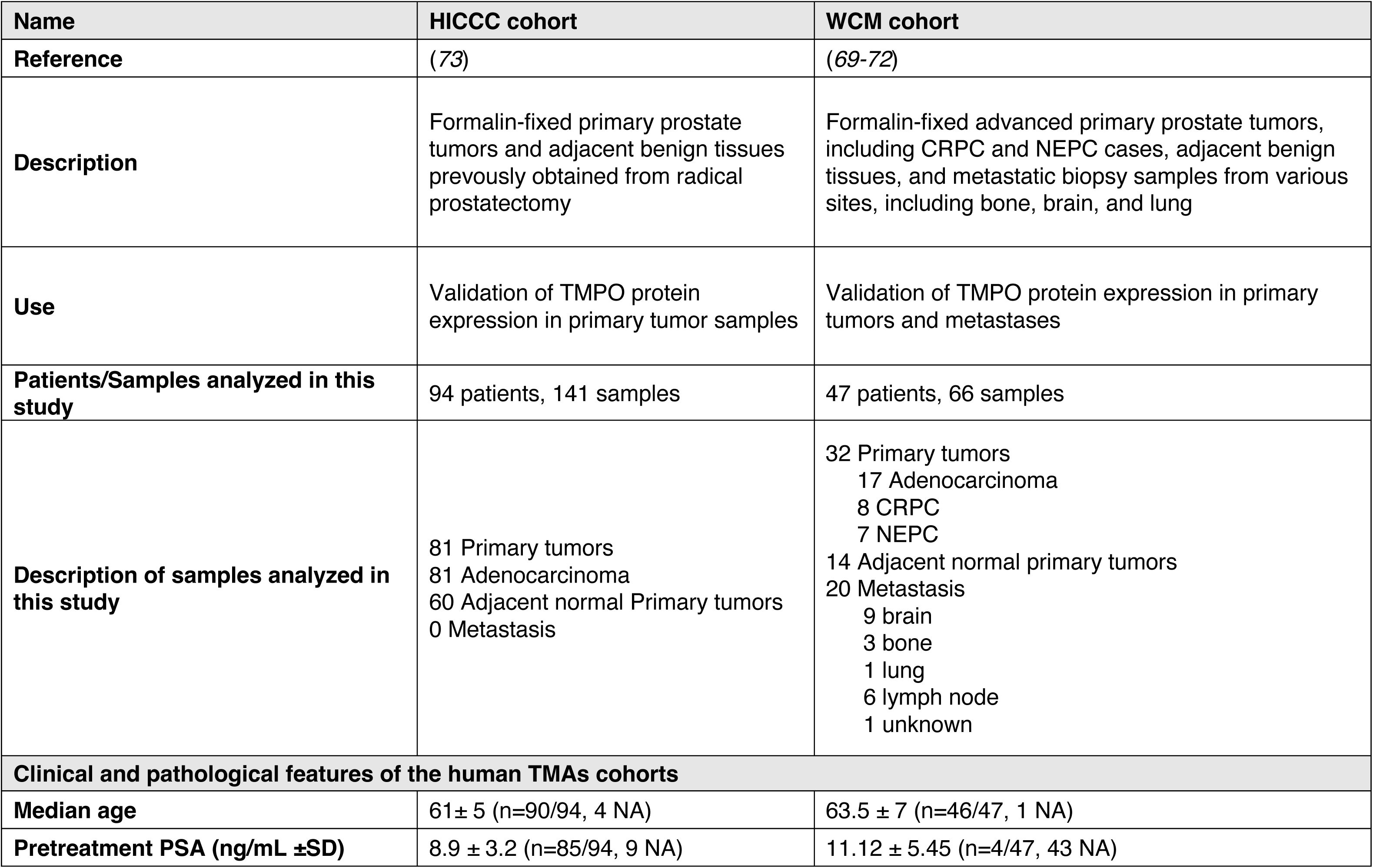
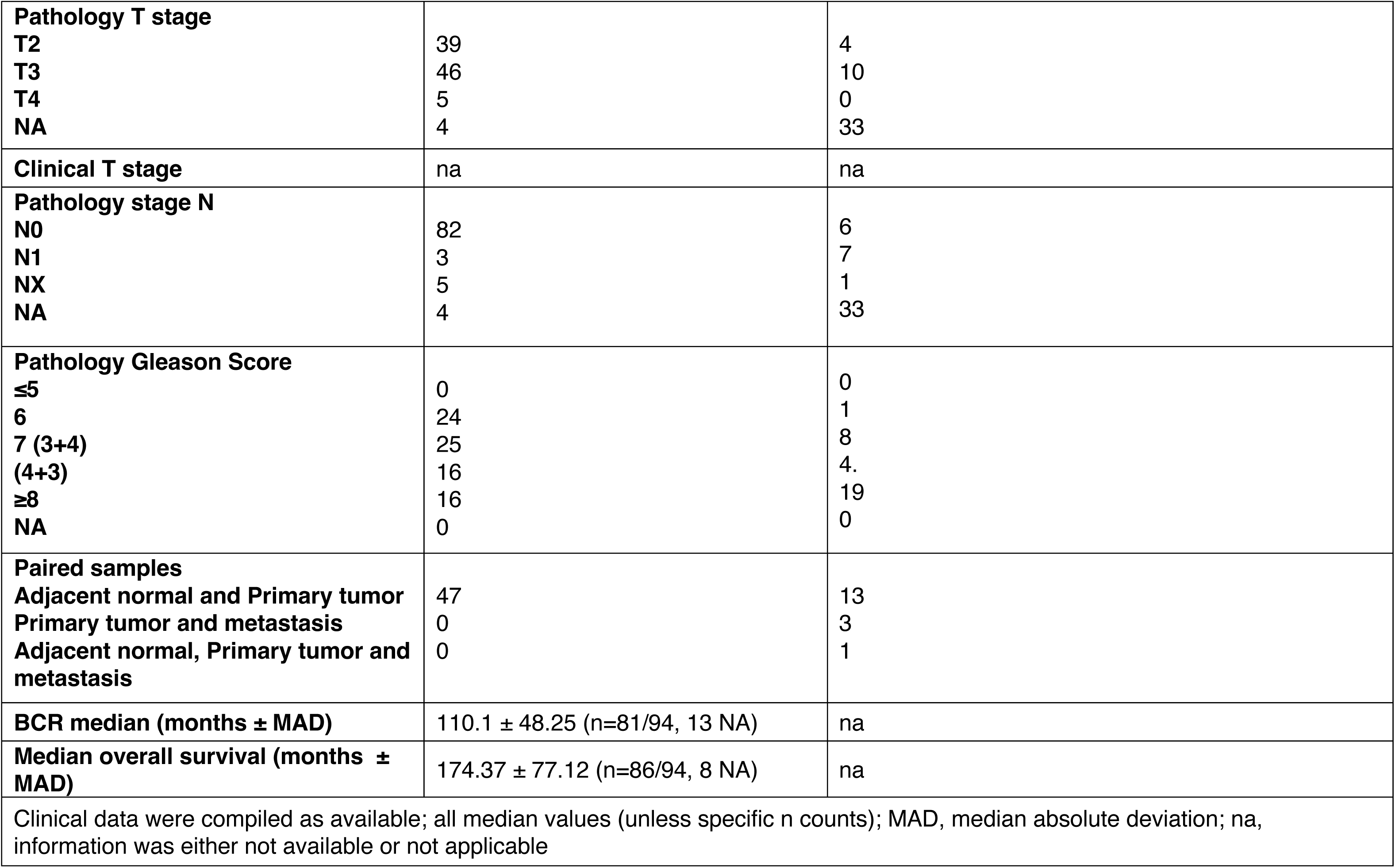
Clinical and pathological features of the human TMAs cohorts. [Related to Figures 6 and S9].

| Name | HICCC cohort | WCM cohort |
| --- | --- | --- |
| Reference | (73) | (69-72) |
| Description | Formalin-fixed primary prostate tumors and adjacent benign tissues previously obtained from radical prostatectomy | Formalin-fixed advanced primary prostate tumors, including CRPC and NEPC cases, adjacent benign tissues, and metastatic biopsy samples from various sites, including bone, brain, and lung |
| Use | Validation of TMPO protein expression in primary tumor samples | Validation of TMPO protein expression in primary tumors and metastases |
| Patients/Samples analyzed in this study | 94 patients, 141 samples | 47 patients, 66 samples |
| Description of samples analyzed in this study | 81 Primary tumors<br>81 Adenocarcinoma<br>60 Adjacent normal Primary tumors<br>0 Metastasis | 32 Primary tumors<br>17 Adenocarcinoma<br>8 CRPC<br>7 NEPC<br>14 Adjacent normal primary tumors<br>20 Metastasis<br>9 brain<br>3 bone<br>1 lung<br>6 lymph node<br>1 unknown |
| **Clinical and pathological features of the human TMAs cohorts** |  |  |
| Median age | 61 ± 5 (n=90/94, 4 NA) | 63.5 ± 7 (n=46/47, 1 NA) |
| Pretreatment PSA (ng/mL ±SD) | 8.9 ± 3.2 (n=85/94, 9 NA) | 11.12 ± 5.45 (n=4/47, 43 NA) |
| **Pathology T stage** |  |  |
| T2 | 39 | 4 |
| T3 | 46 | 10 |
| T4 | 5 | 0 |
| NA | 4 | 33 |
| **Clinical T stage** | na | na |
| **Pathology stage N** |  |  |
| N0 | 82 | 6 |
| N1 | 3 | 7 |
| NX | 5 | 1 |
| NA | 4 | 33 |
| **Pathology Gleason Score** |  |  |
| ≤5 | 0 | 0 |
| 6 | 24 | 1 |
| 7 (3+4) | 25 | 8 |
| (4+3) | 16 | 4. |
| ≥8 | 16 | 19 |
| NA | 0 | 0 |
| **Paired samples** |  |  |
| Adjacent normal and Primary tumor | 47 | 13 |
| Primary tumor and metastasis | 0 | 3 |
| Adjacent normal, Primary tumor and metastasis | 0 | 1 |
| **BCR median (months ± MAD)** | 110.1 ± 48.25 (n=81/94, 13 NA) | na |
| **Median overall survival (months ± MAD)** | 174.37 ± 77.12 (n=86/94, 8 NA) | na |
| Clinical data were compiled as available; all median values (unless specific n counts); MAD, median absolute deviation; na, information was either not available or not applicable |  |  |

**Table S7:** Clinical characteristics of patients for the analysis of CTCs. [Related to Figures 6 and S9].

| Patient ID | Age (years) | Race | Smoking status | Histology | Stage | Metastatic sites | Gleason score | Prior Treatments (n) | Last treatment | Treatment-to-Blood Draw Interval (days) | Disease status |
| --- | --- | --- | --- | --- | --- | --- | --- | --- | --- | --- | --- |
| PRO-CTC-001 | 85 | Caucasian | Former smoker | Prostate adenocarcinoma | IVB | Bone and lymph nodes | 9 | 4 | TVB-2640 + Enzalutamide | 58 | PD |
| PRO-CTC-002 | 59 | Asian | Non smoker | Prostate adenocarcinoma | IVB | Bone and lymph nodes | 4 + 5 | 7 | Carboplatin/ Cabazitaxel | 6 | SD |
| PRO-CTC-003 | 70 | Caucasian | Former smoker | Prostate adenocarcinoma | IVB | Bone and lymph nodes | 8 | 4 | ARX517 (anti-PSMA) | 63 | PD |
| PRO-CTC-004 | 63 | Caucasian | Non smoker | Prostate adenocarcinoma | IVA | Bone, lymph nodes and lungs | 4 + 5 | 2 | Degarelix + Enzalutamide | 28 | PD |
| PRO-CTC-005 | 66 | Caucasian | Former smoker | Prostate adenocarcinoma | IVB | Bone, lymph nodes and bone marrow | 4 + 5 | 4 | <sup>67</sup>Cu-SAR-bisPSMA | 54 | PD |

*PD = progressing disease; PR =partial response; SD = stable disease*

## Materials and Methods

### Genetically engineered mouse models (GEMMs)

#### Description of GEMMs

All experiments using animals were performed according to protocols approved by the Institutional Animal Care and Use Committee (IACUC) at Columbia University Irving Medical Center (CUIMC). The genetically engineered mouse models (GEMMs) used in this study have been described previously (*35–37, 39*) and the multi-allelic strains are available in the Jackson laboratory (Resource Tables R1 and R2): *(i) NP* mice, for *Nkx3.1^CreERT2/+^;Pten^flox/flox^; R26R-CAG-^LSL-EYFP/+^* (*39*); *(ii) NPM* mice, for *Nkx3.1^CreERT2/+^;Pten^flox/flox^; Hi-MYC; R26R-CAG-^LSL-EYFP/+^* (*35*); *(iii) NPp53* mice, for *Nkx3.1^CreERT2/+^; Pten^flox/flox^; Trp53^flox/flox^; R26R-CAG-^LSL-EYFP/+^* (*37*); and *(iv) NPK* mice, for *Nkx3.1^CreERT2/+^; Pten^flox/flox^; Kras^lsl-G12D/+^; R26R-CAG-^LSL-EYFP/+^* (*36*). These mice utilize a tamoxifen-inducible allele, *Nkx3.1^CreERT2^*^/+^, to achieve spatial and temporal regulation of Cre-mediated recombination specifically in luminal prostatic epithelial cells (*40*), and also contain a conditionally-activatable fluorescent reporter allele, *Rosa-CAG-LSL-EYFP-WPRE* (*44*), for lineage marking of tumor cells and their descendants (*36, 37*). A detailed description of all mice used in this study is provided in Dataset 1A.

#### Tumor induction and analysis

All studies were done using littermates that were genotyped prior to tumor induction; since the focus of this study is prostate cancer, only male mice were used. At 2-3 months of age, mice were induced to form tumors by administration of tamoxifen (100 mg/kg in corn oil; Sigma-Aldrich T5648, St. Louis, MO, USA) via oral gavage once daily for 4 consecutive days. Following tumor induction, mice in the end-stage group, including the *NP, NPM*, *NPp53* and *NPK* mice, were monitored regularly and euthanized when their body condition score was <2.0 (*78*), or when they experienced body weight loss ≥ 20% or signs of distress, such as difficulty breathing or bladder obstruction (at 5-8 months), or by one year post-tumor induction. A separate group of *NPK* early-stage mice were euthanized a specific time-point (3-4 months) following tumor induction, prior to the time when these mice develop extensive metastasis. At the time of euthanasia, all mice underwent full necropsy and were analyzed by bright field and fluorescence imaging to visualize tumors and metastases, and for semi-quantitative analyses of metastases, as described below. Blood was collected for analysis of circulating tumor cells (CTCs), tissues were collected and fixed for histological analyses, or fresh tissues were collected for generation of organoids or single cell RNA (scRNA) sequencing.

#### Analysis of metastasis

At the time of sacrifice, YFP-lineage marked prostate tumors and metastases were visualized and photo-documented by *ex vivo* fluorescence, as described in (*36*), using an Olympus SZX16 microscope (Ex 490–500/Em 510–560 filter). Metastases identified via visual detection were confirmed by histopathological analyses (as per below). For detection of bone metastases, muscle and connective tissue surrounding the vertebrae, pelvis, femurs, tibiae, humeri, ulnae, radii and calvariae were removed prior to *ex vivo* fluorescence imaging. A semi-quantitative metastasis score was defined for each mouse by visualization of metastases and micro-metastases using epifluorescence imaging as described in (*36*). Briefly, each mouse was given a metastasis score, assigned from 0 to 4, based on the presence of local versus distant metastases and their relative prevalence and sizes: 0 = no metastases observed; 1 = metastases to lymph nodes only; 2 = micro-metastases to distant organs; 3 = visible metastases to distant organs; 4 = extensive metastases to distant organs.

#### Histopathological analysis

At the time of sacrifice, tissues were fixed in 10% formalin (Fisher Scientific, Hampton NH, USA) and embedded in paraffin. Bones were fixed in 10% formalin for 48 hours followed by decalcification in 15% ethylenediaminetetraacetic acid disodium salt dihydrate (EDTA) (Sigma-Aldrich) solution (pH 7.0) for 3-5 days. Hematoxylin and eosin (H&E) and immunostaining were done using 3 μm paraffin sections as described (*37*). Briefly, sections were deparaffinized in xylene, followed by antigen-retrieval by boiling in Citrate acid-based Antigen Unmasking Solution at pH 7.0 (Vector Labs, Newark, CA, USA). Slides were blocked in 10% normal goat serum, then incubated with primary antibodies overnight at 4 °C, followed by secondary antibodies for 1 hour at room temperature. For immuno-staining, the signal was enhanced using the Vectastain ABC system and visualized with the NovaRed Substrate Kit (Vector Labs, Newark, CA, USA). Slides were counterstained with Hematoxylin and mounted with Permount Mounting medium (Fischer Scientific, Hampton, NH, USA). Immunostained images were captured using an Olympus BX43 microscope. For immunofluorescence staining, slides were incubated with secondary antibodies labeled with Alexa Fluor 488, 555, or 647 (Invitrogen, Thermo Fisher Scientific, Waltham, MA, USA). Sections were counterstained with DAPI (1:1000) (Invitrogen, Thermo Fisher Scientific, Waltham, MA, USA) to visualize nuclei and mounted with Vectashield antifade mounting medium (Vector Labs, Newark, CA, USA). Immunofluorescence images were captured using a Leica Stellaris 5 Lightning spectral confocal microscope (Leica Microsystems, Wetzlar, Germany). All antibodies used in this study are described in Resource Tables R1 and R3).

### Single cell RNA sequencing of mouse prostate tumors

Single-cell RNA-sequencing (scRNA-seq) was performed on prostate tumors from the *NP*, *NPM*, *NPp53*, *NPK* early-stage and *NPK* end-stage mice using a 10X Genomics Chromium platform as described (*36*). Briefly, freshly dissected tissues were enzymatically digested for 15 minutes at 37 °C in a cocktail of pre-warmed 1X collagenase/hyaluronidase, 0.5 U/mL Dispase II and 0.1 mg/mL DNase 1 (STEMCELL Technologies, Vancouver, BC, Canada) in Dulbecco’s Modified Eagle Medium/Nutrient Mixture F-12 (DMEM-F12, Gibco, Thermo Fisher Scientific Waltham, MA, USA), followed by addition of 0.25% Trypsin/EDTA (Gibco, Thermo Fisher Scientific Waltham, MA, USA) for 15 minutes. Cells were resuspended in ice cold DMEM-F12 supplemented with 10% fetal bovine serum (FBS) (Gibco, Thermo Fisher Scientific Waltham, MA, USA), filtered through a 40-μm cell strainer (Falcon, Fisher Scientific, Hampton, NH, USA) and centrifuged at 350 g in an Eppendorf 5810R refrigerated centrifuge for 5 min at 4 °C. The pelleted cells were resuspended in chilled 1X Red Blood Cell Lysis Buffer (Invitrogen eBioscience^TM^, Thermo Fisher Scientific Waltham, MA, USA), incubated for 5 min, and diluted fourfold with cold PBS. Cells were centrifuged as before and resuspended in Dulbecco’s Modified Eagle Medium (DMEM; Gibco) with 10% FBS. Cells were counted using a Countess II Automated Cell Counter (Thermo Fisher Scientific Waltham, MA, USA) and 10,000 cells with a viability greater than 70% were loaded into a 10X Genomics Chromium Controller for cell capture following the 10X Genomics Single Cell Protocol, as per the manufacturer (10X Genomics).

RNA sequencing was performed using Illumina NovaSeq6000. Reads were mapped to the mouse mm10 genome and processed with the CellRanger v.3.1.0 pipeline. Unsupervised clustering was performed based on the Cell Ranger count matrices using the PhenoGraph implementation of Louvain community detection (*79*) after identifying highly variable genes using the drop-out curve, as described previously (*53*). Cells belonging to clusters with high expression of the *eYFP* reporter transcript were defined as tumor cells, as described in (*36*).

To identify gene signatures that distinguish prostate tumors among the GEMMs, we applied probabilistic matrix factorization using a consensus implementation of single-cell hierarchical Poisson factorization (consensus-scHPF) (*53, 80*). The resulting cell score matrix was embedded in two-dimensions using diffusion component analysis (DMAPS; https://github.com/hsidky/dmaps) with Euclidean distance and a kernel bandwidth of 50. To interpret the scHPF-derived gene signatures, enrichment analyses of the top 100 weighted genes for each factor were performed against the MSigDB Hallmark 2020 signature database with Enrichr (*81*). The scRNA-seq data have been deposited in GEO (GSE311017) and lists of differentially-expressed genes and pathways are provided in Dataset 2.

### Isolation and characterization of circulating tumor cells (CTCs)

#### Isolation of circulating tumor cells

At the time of sacrifice, YFP lineage-marked circulating tumor cells (CTCs) were isolated from blood of *NP*, *NPM*, *NPp53*, *NPK* end-stage and *NPK* early-stage mice. To minimize variability associated with circadian rhythm (*82*), the majority of the mice were dissected early in the light phase (between 8:00 AM and 12:00 noon). Immediately postmortem, blood was collected via intracardiac puncture using a 26g needle attached to a 1 ml syringe inserted into the left ventricle. Alternatively, blood was collected from the submandibular vein by performing a shallow puncture with a lancet (Goldenrod animal lancet, Braintree Scientific). Blood samples (200-900 μL) were collected into ethylenediaminetetraacetic acid (EDTA) blood collection tubes (BD Vacutainer, Becton Dickinson, Franklin Lakes, NJ) and incubated for 5 minutes with 10 volumes (10:1 v/v) of 1X RBC Lysis Buffer (Invitrogen eBioscience^TM,^ Thermo Fisher Scientific, Waltham, MA, USA). Lysis of erythrocytes was stopped by addition of 2 volumes (2:1 v/v) phosphate buffered saline (PBS) (Gibco, Thermo Fisher Scientific Waltham, MA, USA) and samples were centrifuged at 400 g in an Eppendorf 5810R refrigerated centrifuge for 5 min at 4° C. The resulting RBC-depleted cell pellets, containing the YFP-lineage marked CTCs and the non-fluorescently labeled white blood cells, were used for all subsequent analyses, unless otherwise indicated, and are referred to as “CTC samples”.

#### Quantification of CTCs

For enumeration, CTC samples were resuspended in PBS (100 μL); 10 μL (10% of the total volume) was transferred to a Countess^TM^ Cell Counting chamber slide (Thermo Fisher Scientific Waltham, MA, USA) and the number of YFP-labeled CTCs in each sample was determined by visual counting of fluorescent cells under the microscope. Representative fluorescence images were acquired with an Olympus IX51 Inverted Microscope (Ex540/Em605 TRITC filter cube) with a XCITE Fluorescence Illumination system (EXFO). For samples with low-levels of YFP-labeled CTCs, CTC samples were resuspended in 100 μl PBS and the samples were transferred in their entirety to a 24-well plate for counting. The total number of CTCs in each sample was determined by multiplying the number of cells by the dilution (e.g., 10X) and reported as number of CTCs/mL.

Alternatively, YFP-lineage marked CTCs were subjected to fluorescence-activated cell sorting (FACS), with gating of the YFP cell population using the APC/FITC (Allophycocyanin/Fluorescein isothiocyanate) channels. Cells were acquired and counted using a BD Influx Cell sorter at the Flow Cytometry Core of the Columbia Center for Translational Immunology (CCTI) and the Herbert Irving Comprehensive Cancer Center (HICCC). The number of CTCs/mL for each mouse were compared to their corresponding tumor weights (in grams) and their metastatic phenotype (metastasis score), and the data were plotted on a log10-transformed x-axis using R (with transformed 0 values plotted as half-points on the y-axis). Spearman correlation analysis of CTC cell number versus tumor weight or metastasis score was computed using the cor.test function from the R *stats* package (v 4.2.2). *P* values were calculated from a two-sided permutation test of 100,000 permutations.

#### Cytospin analysis

RBC-depleted cells were resuspended and counted as above and diluted to 500,000 cells/mL in 1% Bovine Serum Albumin (BSA) in PBS. Cells were pelleted at 400 g for 5 min as above and fixed for 10 minutes in 1 mL of Paraformaldehyde solution 4% in PBS (Santa Cruz Biotechnology, Dallas, TX, USA). Fixed cells were resuspended in 1% BSA in PBS, and 150 μL (75,000 cells) were spun onto Superfrost Plus Microscope slides (Fisher Scientific) using a Thermo Scientific Cystospin 4 Centrifuge. For immunofluorescence staining, the slides were washed with PBS, and the cells were permeabilized with 0.1% Triton X-100 (Sigma-Aldrich, St. Louis, MO, USA) in PBS for 10 min, washed again twice with PBS, and blocked with 10% normal goat serum (Abcam, Cambridge, UK) in PBS for 1 hour. Immunofluorescence staining was done as above.

#### Transplantation of CTCs in vivo

For transplantation studies, a minimum of 200 μL of blood was collected from individual mice. The number of CTCs was determined from 100 μL of the blood sample using the method described above. The remaining 100 μL of the blood sample was processed as above, and the resulting CTC samples were resuspended in 100 μL PBS and injected percutaneously into the left heart ventricle of immunodeficient NCr *nude* (male, Envigo) host mice. Host mice were monitored twice weekly and euthanized when their body condition score was <1.5, or when they experienced body weight loss ≥ 20% or signs of distress, or by 4 months post-implantation. Following euthanasia, the metastatic phenotypes of the host mice were determined by semi-quantitative analyses of *ex vivo* YFP fluorescence with confirmation by subsequent histological analyses, as described above.

### Generation and analyses of CTC-derived organoids

#### Generation of organoids

For generation of organoids from CTCs, RBC-depleted cells were resuspended in organoid culture media (*55*) [hepatocyte culture medium (Corning), 5% heat-inactivated charcoal-stripped fetal bovine serum (CS-FBS) (Gibco), 1X GlutaMAX (Gibco, Thermo Fisher Scientific, Waltham, MA, USA), 10 ng/mL epidermal growth factor (EGF, Corning), 10 µM Y-27632 ROCK inhibitor (STEMCELL Technologies), 100 nM dihydrotestosterone (DHT) (Sigma-Aldrich), 5% Matrigel (Corning) and 1X antibiotic-antimycotic (Gibco)]. The samples were plated onto ultralow-attachment 96-well plates (Corning, 3474) and monitored for organoid-forming efficiency via fluorescence imaging for 10 to 14 days. Alternatively, prior to suspension in organoid media, the CTC samples were depleted of white blood cells using an EasySep Mouse Biotin selection kit (STEMCELL Technologies, Vancouver, BC, Canada), according to the manufacturer’s instructions. Briefly, the CTC samples were labeled with a biotinylated anti-mouse CD45 antibody (BioLegend, San Diego, CA, USA) and CD45-expressing white blood cells were magnetically depleted to enrich for YFP-lineage marked CTCs. To assess the overall incidence of CTC-organoids observed across GEMMs, we calculated the percentage of cases with detectable CTCs that also produced CTC-organoids within each cohort. Organoid-forming efficiency was calculated for each blood sample by counting the number of organoids formed relative to the total number of CTCs 10 days after seeding.

#### Establishment of CTC organoid lines

For passaging, CTC organoids were dissociated by digestion in 1 mL 0.25% Trypsin-EDTA (Gibco) for 10 min at 37°C; digestion was stopped by addition of 2 mL modified Hank’s Balanced Salt Solution (HBSS; STEMCELL Technologies) supplemented with 2% FBS. The dissociated cells were collected by centrifugation at 350g for 5 min at 4°C and resuspended in organoid culture medium, as above. Cells were counted using a Countess II Automated Cell Counter (Thermo Fisher Scientific Waltham, MA, USA) and plated at a seeding density of 5000 cells/well onto ultralow-attachment 96-well plates (Corning). CTC organoids were passaged every four days or when approaching confluence. CTC organoid lines were established following five consecutive passages; a minimum of two CTC organoid lines was established for each of the GEMMs.

For histopathological characterization, CTC organoid lines were harvested by centrifugation at 250g X 5 min at 4°C, fixed in 1 mL 10% formalin at 4 °C overnight (Fisher Scientific, Fair Lawn, NJ), and placed in 80 μL HistoGel (Epredia, Portsmouth, NH, USA) before embedding. Paraffin-embedded organoids were sectioned (3μ) and subjected to histopathological analyses and imaging, as above.

#### scRNA-seq of CTC organoid lines

For scRNA sequencing, CTC organoid lines (passage 10) were dissociated into single cell suspensions and collected as described above. Cells were resuspended in modified Hank’s Balanced Salt Solution (HBSS; STEMCELL Technologies) supplemented with 2% FBS, filtered using a Cell Strainer Snap Cap (35 µm; Falcon) and collected by centrifugation at 300 g in an Eppendorf 5810R centrifuge for 5 min at 4°C. Cells were washed twice with 1 mL of FACS Staining buffer [HBSS supplemented with 1% BSA, 2mM EDTA (Sigma-Aldrich), 10mM HEPES (Sigma-Aldrich)] and cell viability and numbers were assessed using a Countess II Automated Cell Counter (Thermo Fisher Scientific Waltham, MA, USA).

For each sample, 1x10^6^ viable cells were resuspended in 100 µl of FACS staining buffer; 1 µL (0.5 μg) of TotalSeq anti-mouse hashtag antibody (Biolegend, San Diego, CA, USA) (Resource Tables R1 and R3) was added and incubated for 30 min at 4 °C. Cells were washed with 900 µL of FACS staining buffer, collected by centrifugation at 300g x 5 min at 4 °C and resuspended in 1 mL of PBS supplemented with 10% FBS. Cells were counted and an equal number of viable cells (20,000 cells) for each sample, each stained with a different hashed antibody, was combined to a final concentration of 1,000 cells/µL in a total volume of 100 µL PBS with 10% FBS. Cells were subjected to a final cell viability analysis, and 10,000 cells with > 90% viability were loaded into a 10X Genomics Chromium and sequenced as above.

The hashtags were demultiplexed using HashSolo with default settings (*83*). Differential expression analysis was performed as described previously (*84*). Briefly, we randomly subsampled each group to the same number of cells and randomly downsampled the resulting count matrices to the same average number of counts per cell. Differential expression analysis was then performed using the two-sided Mann-Whitney U-test (*mannwhitneyu* function in SciPy) and the resulting p-values were corrected for false discovery using the Benjamini-Hochberg method (*multipletests* function in Statsmodels). Comparison of the differentially expressed genes in CTC organoids with primary tumors was done using gene set enrichment analyses (GSEA) (*85*) with 1,000 permutations (*85*). The scRNA-seq data are deposited in GEO (GSE313119); a list of differentially expressed genes and pathways is provided in Dataset 3.

#### Functional analyses of CTC organoid lines

Tumor growth was determined by subcutaneous implantation of CTC organoid lines into the flank of immunodeficient NCr nude (male, Taconic) host mice. For engraftment, CTC organoids were cultured as above and collected without dissociation. Organoids, corresponding to the equivalent of 2 X 10^5^ cells, were diluted to 100 μL with a 1:1 mixture of Matrigel and PBS and injected subcutaneously into the flank of the host using a 25G 5/8-inch needle. Tumor growth was monitored twice per week and harvested when they reached 2000 mm^3^ or earlier if the body condition score of the host mice was <2.0 or if they exhibit signs of distress; all mice were euthanized by 4 months. Tissues were collected and fixed in 10% formalin (Fisher Scientific, Fair Lawn, NJ) and processed for histopathological analyses as described above.

The metastatic potential of CTC organoid lines was determined following intracardiac injection into immunocompetent C57BL/6 host mice. CTC organoid lines were enzymatically digested at 37°C in 0.25% Trypsin/EDTA (Gibco) for 10 min, resuspended in modified Hank’s Balanced Salt Solution (HBSS; STEMCELL Technologies) supplemented with 2% FBS, and filtered through a 35-µm strainer cap (Corning, Corning, NY,USA), followed by centrifugation at 350 g for 5 min at 4 °C. Cells were resuspended in PBS and counted as above using a Countess II Automated Cell Counter (Thermo Fisher Scientific Waltham, MA, USA). 1 X 10^5^ cells were resuspended in 100 μL PBS and injected percutaneously into the left heart ventricle of the host mice. Mice were euthanized 14 days after injection or sooner if the body condition score was <2.0 (see also above). At the time of sacrifice, metastases were visualized and semi-quantified by ex vivo fluorescence as above. Tissues were fixed in 10% formalin (Fisher Scientific, Fair Lawn, NJ) and metastases were confirmed by histopathological analysis.

### Analyses of individual CTCs (*i*CTCs) and individual CTC-organoids(*i*Organoids)

#### Isolation and sequencing of iCTCs

CTC samples were resuspended in 500 μL of HBSS supplemented with 2% FBS and subjected to FACS using a BD Influx cell sorter (BD Biosciences, Franklin Lakes, NJ) equipped with a 100-μm nozzle and operated at a sheath pressure of 20 psi. The YFP-marked cells were FACS sorted with the BV421/FITC channels and collected at a density of one-cell per well onto 96-well plates for sequencing. The 96-well plates contained 7.5 μL of lysis buffer [0.2 % Triton X-100 (Sigma-Aldrich), SUPERaseIN (1 U/μL) (Thermo Fisher Scientific), 2 mM deoxyribonucleotides (dNTPs) (Thermo Fisher Scientific), and 2 μM reverse transcriptase (RT) primer (Integrated DNA Technologies)] (*86*). As a control, the matched primary tumor and metastases from lung were dissociated into single cells (as described above) and individual YFP-marked cells were FACS sorted in parallel with the *i*CTCs.

#### Generation and sequencing of iOrganoids

To generate organoids from individual CTCs (*i*Organoids), CTCs were FACS sorted as above and collected at a density of one-cell per well onto ultralow-attachment 96-well plates (Corning, 3474) in organoid culture medium, as above. For controls, matched primary tumor and lung metastases were dissociated into single cells and grown as *i*Organoids in parallel with the CTC *i*Organoids. Growth of the *i*Organoids was documented for 8 days by acquiring bright-field and fluorescence images with a 20X objective on an Olympus IX51 inverted Microscope (Ex540/Em605 TRITC filter cube) equipped with an XCITE Fluorescence Illumination system (EXFO). *i*Organoid-forming efficiency was determined by counting visible organoids in each well of the 96-well plate after seven days. Following growth for one week, the *i*Organoids were transferred onto a new 96-well plate containing lysis buffer, as above.

#### Plate sequencing

The 96-well plates containing the *i*CTCs or *i*Organoids were subjected to plate sequencing as described (*86*). For library preparation, we performed template-switching reverse transcription with barcoded oligo(dT) primers followed by full-length cDNA amplification with PCR. We then constructed 3’-end sequencing libraries by tagmentation using an Illumina Nextera kit and sequenced the resulting libraries on an Illumina NextSeq 500 sequencer. Reads were aligned to the mouse reference sequence Gencode m13 (GRCm38.p5) using STAR algorithm (v2.5.3a). The plate seq data are deposited in GEO (GSE313118); a list of genes and pathways is provided in Dataset 4.

For analyses of the *i*Organoid plate seq data, the scHPF algorithm was used to identify factors from the *i*Organoid count matrix as described above. The resulting cell score matrix was embedded using uniform manifold approximation and projection (UMAP) (*87*). Projection of *i*CTC scRNA-seq data into the UMAP embedding for the *i*Organoids was performed by first projecting the *i*CTC count matrix into the scHPF model of the *i*Organoid data using the *project* function in scHPF and then projecting the resulting cell score matrix into the *i*Organoid UMAP embedding using the *transform* function in UMAP.

To identify genes that are correlated with *i*Organoid heterogeneity, inferred based on differences in organoid size, we computed the Spearman correlation coefficient between *i*Organoid area (measured from bright field images of each organoid using Analyze Particles in ImageJ) and expression of each gene (raw counts normalized by total counts for a given organoid). The resulting list of all detected genes ranked by their Spearman correlation with *i*Organoid size constitutes the “size signature” in our analyses (Dataset 4B). To identify pathways associated with *i*Organoid area, we used GSEA (*85*) with all genes ranked by their Spearman correlation coefficient with organoid area (as above) and using gene sets from Hallmarks 2020 in the MSigDB database (*88*).

### Identification of CTC master regulator modules

#### Pre-processing iCTC scRNA seq data for Protein Activity Inference Pipeline

Gene expression count matrices from the *i*CTC data were normalized and scaled in R version 4.4.2 as described above. We regressed out the effect of read count and cell cycle using Seurat SCTransform, with cell cycle annotated by Seurat CellCycleScoring function (*89*). Scaled data from each sample were batch-corrected by Seurat using the functions FindIntegrationAnchors and IntegrateData, with default parameters (*89*). The batch-corrected dataset was projected into its first 50 principal components using the RunPCA function in Seurat, and further reduced into a 2-dimensional visualization space using the RunUMAP function with method umap-learn and Pearson correlation as the distance metric between cells (*89*). Data were clustered using the Louvain algorithm with silhouette score resolution-optimization selecting the resolution with maximum bootstrapped silhouette score in the range of resolution values from 0.01 to 1.0 incremented by 0.01 (*90*).

#### Generation of regulatory networks

We generated a mouse prostate cancer regulatory network using the combined scRNA seq 10X Chromium dataset of mouse prostate tumor cells (described in section entitled, *Single cell RNA sequencing of mouse prostate tumors)*. MetaCells were computed within each gene expression-inferred cluster by summing SCTransform-corrected template counts for the 10 nearest neighbors of each cell by Pearson correlation distance. 200 metaCells per cluster were sampled to compute a regulatory network from each cluster. Regulatory networks were reverse engineered using the ARACNe algorithm exactly as in (*58*).

#### Protein Activity Inference (metaVIPER)

Protein activity was inferred for the iCTCs scRNA data by running the metaVIPER algorithm (*59*), using the ARACNe networks derived as described above. We used the cell cycle regressed iCTC gene signatures generated above as input for metaVIPER. The resulting VIPER matrices included 350 proteins with activity successfully inferred from ARACNe gene regulatory networks. These were loaded into a Seurat Object with CreateSeuratObject (*89*), then projected into the first 50 principal components using the RunPCA function. These were further reduced into a 2-dimensional visualization space using the RunUMAP function with method umap-learn and Pearson correlation as the distance metric between cells. Clustering was performed by resolution-optimized Louvain algorithm. Differential protein activity between clusters (defined as master regulator modules) was computed by t-test, and top proteins for each module were ranked by p-value. These are defined as the proteomic Master Regulators (MRs) (Dataset 5). Data were further visualized by supervised linear discriminant analysis dimensionality reduction using R MASS package lda function (*91*).

#### Enrichment with Size Signature

Based on the size annotations of the CTC organoid dataset (see above), correlation was computed between expression of each gene and annotated organoid size. The resulting vector of correlation values was defined as a gene signature for GSEA. Subsequently, for each cell in the *i*CTC dataset, a set of marker genes was defined as the 50 most-upregulated genes in that cell, ranked by z-score. Normalized Enrichment Score (NES) was computed by GSEA for each cell by enrichment of its marker gene set within the CTC organoid size signature. NES were determined by 1000 random permutations of gene ranking, with distribution of resulting NES values for each cell plotted by module and compared across modules by *t*-test.

### Validation of CTC MR activity on human prostate cancer patient cohorts

#### Human patient cohorts based on transcriptomic data

We used several independent publicly-available human prostate cancer cohorts comprised of primary tumors, metastases, and/or circulating tumor cells (CTCs). We used three independent cohorts of bulk RNA sequencing (bulk Seq) data including primary tumors and metastatic biopsies: The Cancer Genome Atlas (TCGA) cohort, which is comprised of treatment naïve primary tumors (n = 558) (*42*); the Stand Up to Cancer (SU2C) cohort, which is comprised of heavily treated metastatic biopsies (n = 270) (*43*); and the Glinsky cohort, which is comprised of primary tumors from radical prostatectomies in recurrent and non-recurrent patients (n = 79) (*61*). We also queried two independent cohorts of scRNA seq data from primary tumors and metastases: a cohort of metastatic castrate-sensitive prostate cancer (mCSPC) that included pre-treatment biopsies (n = 783 tumor cells from 8 patients)(*63*); and a cohort of primary prostate tumors, which included treatment-naïve prostatectomy specimens (n = 232 tumor cells from 7 patients) (*62*). We used two additional scRNA seq cohorts comprised of human CTC: one (n = 77 tumor cells) from 13 patients with metastatic prostate cancer and with control cells from 12 patients with primary tumors (*64*); and a second comprised of scRNA seq data from CTCs (*n* = 44 tumor cells) from five patients with metastatic prostate cancer (*65*). Details of the clinical and pathological characteristics of all patient cohorts are described in Table S5.

#### Validation of CTC MR modules

To define MR modules, we selected the set of all inferred MR proteins differentially upregulated per module using a Benjamini-Hochberg corrected p-value < 0.05 by Wilcoxon test. In each validation dataset, counts were normalized to log10(Transcripts Per Million), internally scaled by z-score for each, and VIPER protein activity inference was performed using the ARACNe network described above. For each dataset, enrichment of each CTC MR module was computed on a patient-by-patient level by GSEA, and enrichment of individual MR proteins was directly assessed on patient-by-patient level by VIPER NES for that protein.

For association to Gleason score, we used the TCGA cohort. The distribution of enrichment scores for each CTC MR module was plotted against Gleason score to assess for correlation between increased CTC MR module enrichment and Gleason score, by Pearson correlation test. To evaluate enrichment in primary tumor versus metastases or CTCs, we compared the enrichment scores in the relevant human clinical cohorts (as above). Cox regression against recurrence-free survival was performed for each CTC MR module enrichment score. NES were binarized to less than zero (low) or greater than zero (high), and Kaplan-Meier curves showing association with Recurrence-Free Survival in each cohort were generated along with the corresponding log-rank p-value. All quantitative and statistical analyses were performed using the R computational environment version 4.4.2 and survminer statistical package (*92*). Statistical comparisons of enrichment scores were performed by Wilcoxon test with Benjamini-Hochberg multiple testing correction where appropriate, and survival analyses were performed by log-rank test and cox regression. In all cases, statistical significance was defined as an adjusted p-value less than 0.05.

#### Validation of TMPO MR activity

To evaluate the expression and activity levels of TMPO in the above described human patient cohorts, we performed analyses as above using binarized TMPO gene expression and VIPER protein activity in each dataset, with low vs high cutoff selected by log-rank maximization. To evaluate the expression status of the TMPO regulon in these cohorts, TMPO gene expression and VIPER activity were compared between TCGA and SU2C by Wilcox test. Positively regulated TMPO targets from the ARACNe regulon were overlapped with statistically significant differentially expressed genes between TCGA and SU2C (by Wilcox test with Bonferroni-corrected p-value <0.05 and fold-change > 0.5) and visualized in a gene expression heatmap. Activity of TMPO was further visualized as violin plots, grouping CTCs and Primary vs Metastatic tumor cells in the datasets described above.

### Validation of TMPO protein expression on human tissue microarrays (TMAs)

#### Description of TMAs

All studies involving human tissue specimens were performed according to protocols approved by the Human Research Protection Office and Institutional Review Board at the Columbia University Irving Medical Center (CUIMC) and Weill Cornell Medicine (WCM). Only anonymized tissues were used, and patient consent was obtained. We used several TMAs for these studies. The HICCC TMA cohort includes all Gleason (scores 6 to 9) primary prostate tumors (n = 81) and adjacent benign tissues (n = 60) collected over a 10-year period from patients who underwent radical prostatectomy in the Department of Urology at CUIMC (*73, 93*). The WCM cohort consisted of primary tumor samples from various stages of prostate cancer, as well as metastatic samples, collected from 47 patients (*69–72*). The primary tumor samples (n = 32) include benign prostate tissue (n = 14), prostate adenocarcinoma (n = 17), castration-resistance prostate cancer (CRPC) (n = 8), and neuroendocrine prostate cancer (NEPC) (n = 7). The metastatic samples (n = 20) were biopsy specimens obtained from various anatomical sites, including bone (n = 3), brain (*n* = 9), lung (n = 1) and lymph nodes (n = 6). TMAs were constructed (Beecher Instruments, MD, USA) by punching triplicates cores of 1 mm for each sample (*73*). Details of the clinical and pathological features of the TMAs are summarized in Table S6.

#### Analyses of TMPO expression on TMAs

Immunohistochemistry was performed by the CUIMC Molecular Pathology Shared Resource using a rabbit antibody directed against the protein encoded by TMPO, namely LAP2 (Abcam). Briefly, the TMA slides were deparaffinized in xylene, followed by heat-induced antigen-retrieval in 0.01 M Citrate based buffer solution at pH 6.0 for 16 min. Slides were blocked in 10% normal goat serum for 25 min, then incubated with anti-LAP2 primary antibody (Abcam, Cambridge, MA, USA, cat# ab5162) at 1:4000 dilution in antibody diluent (Dako, cat#S0809, Agilent Technologies, Santa Clara, CA, USA) for two hours at room temperature. The slides were incubated with HRP (Horseradish Peroxidase)-labelled polymer anti-Rabbit (Dako, cat#K4003, Agilent Technologies, Santa Clara, CA, USA) for 45 min at room temperature. Slides were counterstained with Hematoxylin, dehydrated and mounted using Permount Mounting medium (Fisher Scientific, Hampton, NH, USA) Immunostaining was evaluated by two pathologists (G.N.F. and M.L.). Analysis of TMPO (LAP2) expression was performed on digitized whole-slide images acquired using a Leica GT450 high-throughput scanner (Leica Biosystems, Wetzlar, Germany). Image analysis was conducted with HALO (Indica Labs, v4.0) and HALO AI module. First, individual TMA cores were segmented, excluding those lacking epithelial components, containing artifacts, or with insufficient tissue. A trained epithelial–stromal classifier was then applied to distinguish tumor from non-tumor regions. Cell segmentation was performed on a cell-by-cell basis, with compartmentalization into nuclear, cytoplasmic, and membrane regions, and TMPO (LAP2) expression was quantified exclusively in the nuclear compartment of neoplastic epithelial cells. Raw data were exported, and for each core the percentage of TMPO (LAP2)-positive tumor cells was calculated. For each patient, mean values across replicate cores were obtained. Comparative analyses were subsequently conducted between benign, primary, and metastatic samples and further stratified into benign, adenocarcinoma, castration-resistant prostate cancer (CRPC), and neuroendocrine prostate cancer (NEPC) groups. Overall, TMPO (LAP2) expression was analyzed across 207 samples, including 74 benign, 113 primary tumors, and 20 metastases. Statistical analyses were performed in R (v 4.5.1), Data distribution was assessed using the Shapiro–Wilk test. For comparisons between more than two groups, Kruskal–Wallis tests followed by Dunn’s post-hoc multiple comparisons with Bonferroni correction were applied. For pairwise group comparisons, Mann–Whitney U tests were used. Results were considered statistically significant at *P* < 0.05.

### Analysis of Human CTCs

#### Description of patients

All studies involving human tissue specimens were performed according to protocols approved by the Human Research Protection Office and Institutional Review Board at WCM. Only anonymized tissues were used, and patient consent was obtained. We analyzed CTCs from five patients with metastatic CRPC. The clinical characteristics of the patients enrolled for the analysis of CTCs are described in Table S7.

#### Enrichment for CTCs

Peripheral blood samples (∼10mL) were collected from patients into EDTA tubes and processed for CTC isolation within 24 hr of collection. CTCs were enriched using a size-based microfluidic approach with the Genesis System with Celselect Slides (Genesis System, Bio-Rad), according to the manufacturer’s protocol. Briefly, 8 mL of blood was diluted 1:1 with PBS and loaded a cartridge pre-primed with 8 mL ethanol followed by 8 mL PBS. CTCs retained within the cartridge were automatically washed and flushed into the collection funnel with 4 mL of dilution buffer pre-warmed to room temperature. Collected cells were gently centrifuged at 300 x g (Eppendorf 5702 R) for 5 min at room temperature. The supernatant was discarded and the pellet was resuspended in 1 mL of PBS containing 2% FBS and kept on ice until further use. In parallel, peripheral blood mononuclear cells (PBMCs) were isolated from 2 mL of matching patient blood using density gradient centrifugation with Ficoll-Paque (GE Healthcare). Briefly, blood was diluted 1:1 with PBS, gently layered over Ficoll in a 15 mL conical tube, and centrifuged at 400 × g for 30 minutes at room temperature (no brake). The PBMC layer was collected, washed twice with PBS, and resuspended in PBS containing 2% FBS for downstream applications.

#### Immunofluorescence staining

CTC and PBMC suspensions were seeded onto 8 mm Cell-Tak-coated coverslips (Corning) prepared according to the manufactureŕs instructions. Coverslips, placed in a 48-well plate, were centrifuged at 200 x g (Eppendorf 5810 R) for 10 min at room temperature, and incubated at 37°C with 5% CO_2_ for 30 minutes. Cells were then fixed in 300 µL 1:1 formaldehyde: PHEM buffer (60 mM PIPES, 25 mM HEPES, 10 mM EGTA, 2 mM MgCl₂) for 15 min, and blocked overnight with 10% normal goat serum (Jackson ImmunoResearch, West Grove, PA, USA) and 6% Bovine Serum Albumin (BSA) in PBS (Corning, Corning, NY, USA), as previously described (*94, 95*). Cells were sequentially immuno-stained with antibodies againstCD45 (mouse anti-CD45, QDot800-conjugated; Invitrogen), Lap2α (TMPO, rabbit anti-Lap2α, Abcam, followed by Alexa Fluor 647–conjugated secondary), Lamin A/C (mouse anti–Lamin A/C, Santa Cruz, followed by Alexa Fluor 488–conjugated secondary), and either pan-cytokeratin (mouse anti–pan-cytokeratin clone C-11, BioLegend, directly conjugated with CF594 using Biotium Mix-n-Stain) or PSMA (rabbit anti-PSMA, abcam, followed by Alexa-Fluor 647-conjugated secondary). Nuclei were counterstained with DAPI (Thermo Fisher Scientific). Cells were incubated with primary or directly conjugated antibodies for 1 hour at room temperature, washed three times with PBS, and—when applicable—incubated with secondary antibodies for 30 minutes under the same conditions, followed by three additional PBS washes. Coverslips were mounted in Mowiol mounting medium (Sigma-Aldrich), placed on glass slides, and allowed to cure overnight before imaging. Details of primary and secondary antibodies used in this study are provided in Resource Tables R1 and R3.

#### CTC Imaging

CTCs were imaged on a Nikon Eclipse Ti2-E inverted microscope equipped with a Yokogawa CSU-W1 SoRa spinning-disk confocal system paired with two 3^rd^-generation sCMOS cameras (Hamamatsu ORCA-Fusion BT). The microscope was configured with Nikon Lambda D objective series at low (4x/10x/20x) and high (40x/60x) magnifications, enabling efficient sample screening without oversampling. CTCs were first imaged at 20X magnification across five fluorescent channels. Fluorescence intensity thresholds for each channel were determined from positive and negative controls included in each staining experiment. Images were analyzed with NIS-Elements GA3 software (Nikon Instruments). Each coverslip was scanned in its entirety at a single interpolated focal plane, generating ∼250 image tiles across five fluorescence channels. Nucleated cells were segmented using a global threshold of the DAPI channel to generate binary masks. Cells were scored as putative CTCs if they met the established DAPI⁺/CK-PSMA⁺/CD45⁻ phenotype. X–Y coordinates of candidate CTCs were automatically marked using the NIS-Elements GA3 software, and single CTCs were subsequently imaged at high resolution using a 60×/0.4 NA objective and the super-resolution 4× SoRa scanning mode.

### Functional validation of master regulators

#### Generation of CTC cell lines

Functional validation studies were done using the established 3-D CTC organoid lines, as described above, or 2-D cell lines derived from the established from CTC organoid lines. To generate the 2-D cell lines, CTC organoid lines (passage 5) were enzymatically digested at 37°C in 0.25% Trypsin/EDTA (Gibco) for 10 min, resuspended in modified Hank’s Balanced Salt Solution (HBSS; STEMCELL Technologies) supplemented with 2% FBS, and filtered through a 35-µm strainer cap (Corning, Corning, NY,USA), followed by centrifugation at 350 g for 5 min at 4 °C. The dissociated cells were plated directly onto 100 mm Corning^TM^ Primaria tissue culture dishes (Fisher Scientific, Hampton, NH, USA) in RPMI-1640 medium supplemented with 10% FBS and 1X antibiotic-antimycotic (Gibco) and propagated for five passages. To avoid clonal variation, pooled cell population was used in these studies. Each CTC cell line stock was established at passage five and used for experimental assays within three passages from thawing. Cell lines were tested using a Universal Mycoplasma Detection Kit (ATCC # 30-1012K). The tumor and metastatic phenotypes of the 2-D cell lines were compared with the parental CTC organoid lines, as described below.

#### Generation of the lentiviral vectors

The LT3GEPIR lentiviral backbone (Tet-ON miR-E (miR-30 variant)-based RNAi) was purchased from Addgene (Plasmid#111177) (*96*), and was modified by replacing the EGFP with the DsRed fluorescent marker from the donor backbone LT3REVIR (Addgene, Plamid#111176) by cloning into unique BamHI and XhoI sites in the vector. The modified LT3GEPIR-DsRed lentiviral vector maintains the shRNA targeting the Renilla Luciferase (Ren.713), drug selection (Puromycin) and other key features of the LT3GEPIR backbone (*96*), but expresses red fluorescence coupled with miR-E shRNAs from a Tet-responsive element promoter (T3G). For shRNA-mediated silencing of candidate genes, we selected the three top-ranked Sensor-based shRNAs predictions for each gene as published in (*96*). A complete list of the selected sensor-based shRNAs predictions is provided in Resource Table R4.

For cloning the miR-E shRNA cassettes into the recipient miR-E based LT3GEPIR-DsRed backbone, 97-mer oligonucleotides coding for the respective shRNAs sequence were PCR based amplified, followed by PCR purification using the using the QIAquick PCR purification kit (Qiagen, Hilden, Germany). The miR-E based LT3GEPIR-DsRed backbone was then digested with EcoRI/XhoI enzymes (New England BioLabs) and gel purified using the MinElute Gel extraction kit (Qiagen, Hilden, Germany). The amplified products were cloned into the corresponding EcoRI/XhoI sites of the miR-E LT3GEPIR-DsRed backbone. All final vectors were sequenced-confirmed using the miREseq primer 5’-TGTTTGAATGAGGCTTCAGTAC-3’. The sequences of the PCR primers used for cloning are provided in the Resource Table R5.

#### Production of virus

All procedures for using lentiviruses were done according to approved procedures by the Office of Environmental Health and Safety at CUIMC. The miR-E LT3GEPIR-DsRed lentiviruses, described above, were generated using second-generation packaging vectors (psPAX2 and pMD2.G, Addgene) produced in HEK-293T cells (ATCC), and concentrated using the Lenti-X Concentrator reagent (Clontech) (*36, 72*). Upon infection of cell lines or organoids with the miR-E LT3GEPIR-DsRed based lentiviruses, puromycin was added in the culture medium (5 µg/mL) and cells were selected for 4 days. Following removal of the puromycin, doxycycline (0.5 µg/mL) was added for 2 consecutive days to induce shRNA-mediated silencing. Cells were subjected to fluorescence-activated cell sorting (FACS) to enrich for the population of cells co-expressing RFP and YFP fluorescence using the PE-TR/FITC (Phycoerythrin-Texas Red/Fluorescein isothiocyanate) channels. Cells were acquired on a BD Influx Cell sorter at the Flow Cytometry Core of the Columbia Center for Translational Immunology (CCTI) and Herbert Irving Comprehensive Cancer Center (HICCC).

#### Quantitative real-time PCR

Quantitative real time PCR analysis was performed by using the QuantiTect SYBR Green PCR kit (Qiagen, Hilden, Germany) using mouse *Gapdh* as control. Relative expression levels were calculated using the 2^-ΔΔCT^ method, as described (*93*). Sequences of all primers used in this study are provided in Resource Table R5.

#### Western blot analysis

Immunoblot analysis was performed using whole protein extracts obtained with 1X radioimmunoprecipitation assay (RIPA) buffer (0.1 % SDS, 1 % deoxycolate sodium salt, 1.0 % Triton-X 100, 0.15 M NaCl, 10 mM Tris-HCl (pH 7.5), 1 mM EDTA supplemented with 1% protease inhibitor cocktail (Roche), 1 % phosphatase inhibitor (Sigma-Aldrich, St. Louis, MO, USA) and 0.5 % PMSF (Sigma-Aldrich, St. Lous, MO, USA). Protein lysates (5 μg per lane) were resolved by SDS-PAGE, followed by immunoblotting with the appropriate primary and secondary antibodies, and visualized using an ECL Plus Western Blotting Detection Kit (GE Healthcare/Amersham Biosciences, Little Chalfont, United Kingdom). Details of primary and secondary antibodies used in this study are provided in Resource Tables R1 and R3.

#### In vitro studies

Colony formation assays were performed by plating CTC cell lines (200 cells per well) in six-well tissue culture plates. Colonies were visualized by staining with crystal violet and quantified using ImageJ software (https://imagej.net/ij/index.html).

For migration and invasion assays, 2.5 X 10^4^ cells per well from CTC cell lines were seeded into either control inserts (Corning Biocoat Control Insert) or matrigel-coated inserts (Corning Biocoat^TM^ Matrigel^TM^ Invasion Chamber with Corning^TM^ Matrigel Matrix) in FBS-free RPMI-1640 medium, using 24-well cell culture plates. RPMI-1640 medium supplemented with 10% FBS was added to the lower chamber as a chemoattractant. Cells were incubated for 18 hours at 37^°^C in 5% CO_2._ Following incubation, inserts were stained with crystal violet. Five random fields/view per insert were imaged using an Olympus BX43 microscope and quantified using ImageJ software. All *in vitro* studies were performed in quadruplicate and with a minimum of two independent biological replicates.

For 3-(4,5-dimethylthiazol-2-yl)-2,5-diphenyltetrazolium bromide (MTT)-based proliferation assays, 5000 cells per well from CTC organoids were dissociated and seeded in quadruplicate in 96-well plates. Cells were grown for up to 72 hours using the Cell Proliferation kit I (MTT, Roche) according to the manufacturer’s instruction. MTT-based proliferation was measured using absorbance of A560 and quantified using the SpectraMax iD5 plate reader (Molecular Devices).

#### In vivo studies

Functional analyses were performed using either in the 3-D CTC organoid lines or 2-D CTC cell lines that were infected with the lentiviruses as described above. For tumor growth assays, 5 X 10^5^ of CTC cells were injected in the flank of a male NCr *nude* mice (Envigo). Alternatively, CTC organoids (not dissociated) were implanted into the flank as described above. To maintain shRNA expression, doxycycline was administered three times per week and provided in the drinking water at a concentration of 2 mg/mL. Tumors were monitored by caliper measurement twice weekly and tumor volumes were calculated using the formula [Volume = (width)^2^ x length/2]. Mice were sacrificed when the tumor size reached 2000 mm^3^ or if the body condition score of the host mice was <2.0 or if they exhibited signs of distress. At the time of sacrifice, blood samples were collected via intracardiac puncture for CTC enumeration (as detailed above). YFP-positive allograft tumors and distant metastases were visualized by *ex vivo* fluorescence using an Olympus SZX16 microscope (Ex490–500/Em510–560 filter).

The metastatic potential of CTC-organoids (dissociated) or CTC cell lines was determined following intracardiac injection into immunocompetent C57BL/6 host mice as described above. 1 X 10^5^ cells were resuspended in 100 μL PBS and injected percutaneously into the left heart ventricle of immunocompetent C57B/L 6 mice. Mice were euthanized 14 days after injection or sooner if the body condition score was <2.0. At the time of sacrifice, metastases were visualized and semi-quantified by *ex vivo* fluorescence, as above. Tissues were fixed in 10% formalin (Fisher Scientific, Fair Lawn, NJ) and metastases were confirmed by histopathological analysis, as above.

### Metabolic analyses

#### Seahorse assay

Oxygen consumption rate (OCR) was quantified using a Seahorse XF96 (Agilent Technologies, Santa Cruz, CA, USA). Briefly, CTC cell lines were seeded at a density of 2 x 10^4^ cells per well into XFe96/XF Pro Cell Culture Microplates (Agilent, Santa Clara, CA, USA, Cat# 103792-100). The culture medium supplemented with doxycycline (0.1 μg/mL) was replaced after 24 hours with bicarbonate-free assay medium (Agilent, Santa Clara, CA, USA, Cat# 103680-100) and incubated for one hour at 37°C in a CO_2_ free incubator. Oxygen consumption rate (OCR) was measured using an XF96 Extracellular Flux Analyzer (Seahorse Biosciences, Billerica, MA, USA) under basal conditions, followed by sequential injections of oligomycin (1 µM; Millipore Sigma cat#75351), carbonyl cyanide-4-(trifluoromethoxy) phenylhydrazone FCCP (1 µM; Millipore Sigma; Cat # C2920), and Rotenone (1 µM; Millipore Sigma, Cat # 557368), according to the manufacturer’s instructions.

#### Anoikis assay

To evaluate cell viability following detachment-induced stress (anoikis), CTC cell lines expressing inducible shRNAs were pre-treated with doxycycline (0.5 μg/ml) for 48hrs. Cells were detached using 0.05% Trypsin-EDTA (Gibco), pelleted, and counted using a hemocytometer. 1 X 10^6^ cells were resuspended in 5 mL of complete medium supplemented with doxycycline (0.5 μg/ml) and transferred into a 15-ml Falcon conical tube (Corning). Tubes were mounted on a Thermo Scientific Tube Revolver Rotator and incubated at 37°C to maintain cells in suspension with rotation. At the indicated time point (0 and 48 hours), 1 mL aliquots were collected for viability assessment using the CellTiter-Glo Luminescent Cell Viability Assay (Promega, Madison, WI, USA) according to the manufacturer’s instructions. Each condition was assayed in quintuplicate, and experiments were performed in three independent biological replicates. Luminescence was recorded using a SpectraMax iD5 plate reader (Molecular Devices, San Jose, CA, USA). For immunofluorescence analysis, the remaining suspension was cytospun onto Superfrost Plus Microscope slides (Fisher Scientific) using a Thermo Scientific Cystospin 4 Centrifuge. Cells were fixed for 10 minutes in paraformaldehyde solution 4% in PBS (Santa Cruz Biotechnology, Dallas, TX, USA) and then subjected to immunofluorescence staining as described above.

#### Hypoxia assay

To measure cell viability under hypoxic conditions, CTC cell lines were seeded at a density of 1000 cells per well in 96-well plates and cultured in a hypoxia chamber for 48 hours. Hypoxic culture was performed in an O₂ Control InVitro Glove Box (Coy Laboratory Products, Inc.) connected to N₂ and CO₂ tanks, maintained at 0.5% O₂, 5% CO₂, and 37 °C. Control cells were maintained under normoxic conditions (21% O_2_). Cell viability was assessed using the CellTiter-Glo Luminescent Cell Viability Assay (Promega, Madison, WI, USA) according to the manufacturer’s instructions. Luminescence was recorded using a SpectraMax iD5 plate reader (Molecular Devices, San Jose, CA, USA). Each time point was assayed in quintuplicate, and the experiment was performed in two independent biological replicates. For immunofluorescence analysis, cells were seeded onto Falcon 4-well culture slides (Corning) at 5000 cells per well and cultured either in the hypoxia chamber (0.5% O_2_) or maintained under normoxic conditions for 48 hours. After incubation, the cells were fixed in Paraformaldehyde solution 4% in PBS (Santa Cruz Biotechnology, Dallas, TX, USA). Immunofluorescence staining was done as above.

### Statistical Analysis

For in vivo studies, survival curves from mouse models were generated using Kaplan-Meier analysis, and statistical significance was calculated using a two tailed Log-rank test. The significance between differences in tumor weight across mouse models was calculated using a Kruskal-Wallis test with Dunn’s multiple comparisons correction for multiple-group comparisons or relative to control group. Statistical analyses of CTC enumeration and combined metastasis scores were performed using a non-parametric unpaired *t* test (Mann-Whitney test) for two-group comparisons, or the Kruskal-Wallis test with Dunn’s multiple comparisons correction for multiple-group comparisons, as indicated in each figure legend. Metastasis frequencies were compared using the Fisher’s exact test or chi-squared test, when appropriate. Correlation analyses were conducted using the Spearman correlation test. For in vitro functional studies, statistical analyses were performed using a non-parametric unpaired *t* test (Mann-Whitney test), one-way ANOVA with Dunnett’s multiple comparisons, two -way ANOVA test or Student’s t-test as specified in the figure legends. All statistical analyses and data visualizations were performed using GraphPad Prism software (Versions 9.5.0 and 10.6.0). For all bar-graphs and dot-plots, means are represented and error bars represent the standard error of the mean (SEM), unless otherwise noted. Statistical analyses of RNA sequencing data were provided above.

### Accession Numbers/ Data availability

All relevant data supporting this study are provided in this paper, and reagents will be made available upon request. All sequencing data have been deposited in GEO, including the 10X scRNA data from the mouse primary tumors (GSE311017); the plate-seq data from the mouse *i*CTC and *i*Organoids (GSE313118); the scRNA-seq data of mouse derived CTC organoids (*NPK* early-stage and end-stage and *shTmpo*-silenced organoids) (GSE313119). Source data are provided with this paper.

